# A generative algorithm for *de novo* design of proteins with diverse pocket structures

**DOI:** 10.1101/2020.03.23.003913

**Authors:** Benjamin Basanta, Matthew J Bick, Asim K Bera, Christoffer Norn, Cameron M Chow, Lauren P Carter, Inna Goreshnick, Frank Dimaio, David Baker

**Affiliations:** Institute for Protein Design, University of Washington, Seattle, WA 98195, USA; Biochemistry Department, School of Medicine, University of Washington, Seattle, WA 98195, USA; Department of Integrative Structural and Computational Biology, The Scripps Research Institute, La Jolla, CA 92037, USA; Lyell Immunopharma, Seattle, WA 98109, USA; Howard Hughes Medical Institute, Seattle, WA 98195, USA

**Author notes:** Corresponding author: David Baker.

## Abstract

To create new enzymes and biosensors from scratch, precise control over the structure of small molecule binding sites is of paramount importance, but systematically designing arbitrary protein pocket shapes and sizes remains an outstanding challenge. Using the NTF2-like structural superfamily as a model system, we developed a generative algorithm for creating a virtually unlimited number of *de novo* proteins supporting diverse pocket structures. The generative algorithm was tested and refined through feedback from two rounds of large scale experimental testing, involving in total, the assembly of synthetic genes encoding 7896 generated designs and assessment of their stability on the yeast cell surface, detailed biophysical characterization of 64 designs, and crystal structures of 5 designs. The refined algorithm generates proteins that remain folded at high temperatures and exhibit more pocket diversity than naturally occurring NTF2-like proteins. We expect this approach to transform the design of small molecule sensors and enzymes by enabling the creation of binding and active site geometries much more optimal for specific design challenges than is accessible by repurposing the limited number of naturally occurring NTF2-like proteins.

## Introduction

Proteins from the NTF2-like structural superfamily consist of an elongated β-sheet that, along with three helices, forms a cone-shaped structure with a pocket (Figure 1.A). This simple architecture is highly adaptable, as evidenced by the low sequence homology among its members, and the many different functions they carry out (1). Natural NTF2-like proteins have been repurposed for new functions through design (2–4), further showing the adaptability of this fold. General principles for designing proteins with curved beta sheets have been elucidated, and used to design several *de novo* NTF2-like proteins (5).

**Figure 1.**
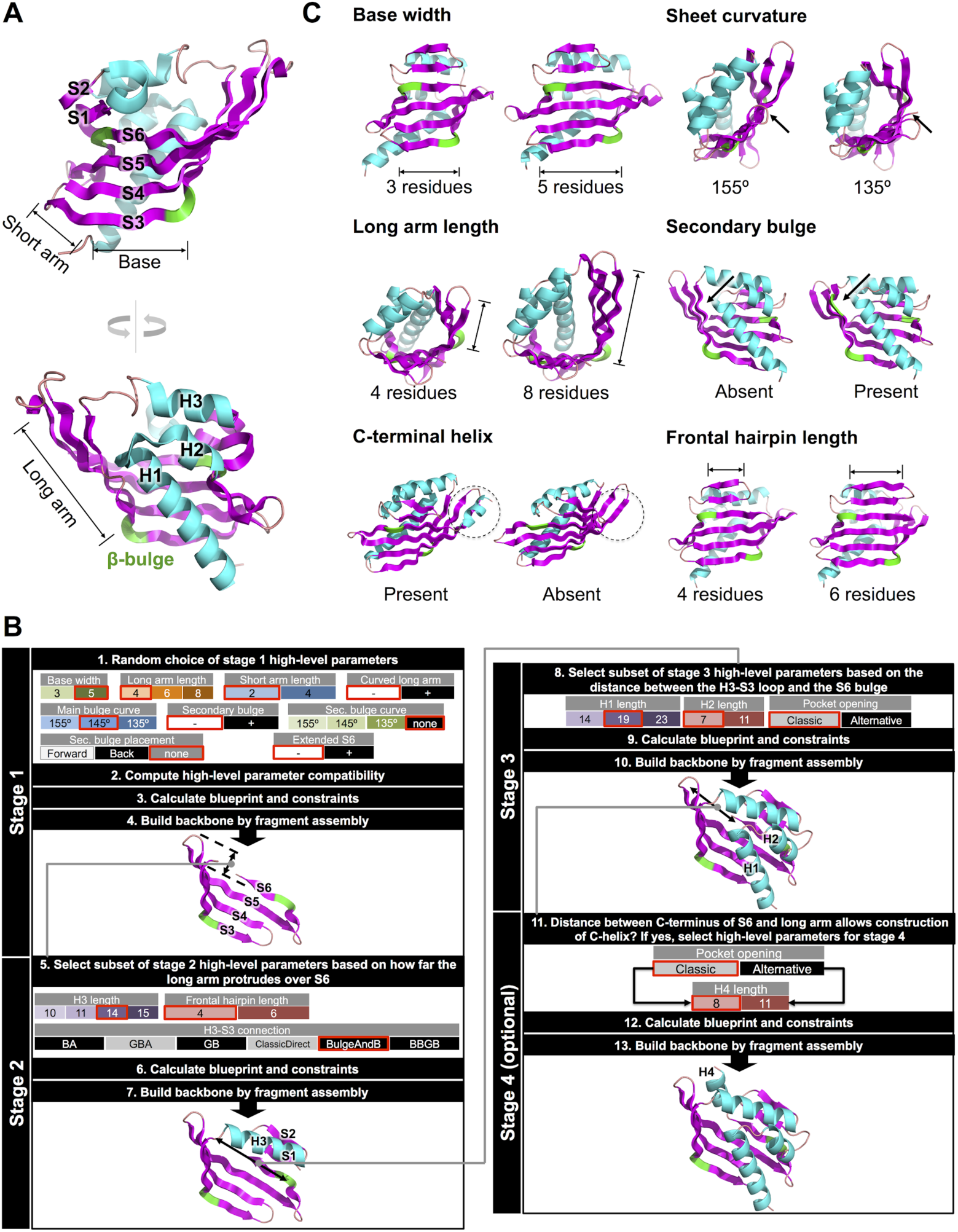
High-level description of the NTF2 generative algorithm. **A:** Canonical NTF2-like structural elements, labeled on the structure of scytalone dehydratase from *Magnaporthe grisea* (PDB 1IDP). **B:** Generative algorithm diagram, depicting hierarchical backbone assembly, and sampling of high-level parameters and local structure variation at each step. **C:** Examples of fold parameters sampled at the higher levels, and structures representing two extreme values for each.

*De novo* design of protein function starts with an abstract description of an ideal functional site geometry (for example, a catalytic active site), and seeks to identify a protein backbone conformation that can harbor the site. The extent to which the ideal site can be realized depends on the number and diversity of backbone conformations that can be searched (6, 7). A promise of *de novo* protein design is to generate a far larger and more diverse set of designable backbones for function than is available in the largest public protein structure database, the Protein Data Bank (PDB) (8, 9). This has been achieved for protein-protein binding due to the simplicity of small globular proteins (10). However, protein structures with pockets are considerably more complex, and since only a small number of *de novo* designed pocket-containing proteins have been characterized, this vision has not yet been realized for small molecule binder or enzyme design. Here we construct a generative algorithm for NTF2-like proteins that samples the structural space available to the fold systematically and widely, and show that the generated protein scaffolds surpass native NTF2-like proteins in pocket diversity.

*De novo* protein design is a two-step process: first, a protein backbone conformation is generated, and second, low energy amino acid sequences for this backbone are found by combinatorial side-chain packing calculations. In Rosetta (11, 12), new backbones can be constructed by Monte Carlo assembly of short peptide fragments based on a structure “blueprint”, which describes the length of the secondary structure elements, strand pairings, and backbone torsion ranges for each residue (13, 14). Because this process is stochastic, each structure generated is distinct. We previously showed that NTF2-like proteins can be designed from scratch using this approach (5), but the diversity and number of designs (on the order of tens) to date is too limited to provide pockets for arbitrary function design. For a given blueprint, the resulting set of structures is generally more homogeneous than that observed in naturally occurring proteins within a protein family, where differences in secondary structure lengths and tertiary structure give rise to considerable diversity. Hence while large numbers of backbones can be generated for a particular blueprint, for example, those previously used to design NTF2-like proteins, the overall structural diversity will be limited.

## Results

### The NTF2 generative algorithm

To access a much broader range of protein backbones, we sought to develop a generative algorithm which samples a wider diversity of structures than natural NTF2-like proteins by carrying out backbone sampling at two levels (Figure 1.B). At the top level, sampling is carried out in the space of high-level parameters that define the overall properties of the NTF2 fold: for example, the overall sheet length and curvature, the lengths of the helices that complement the sheet, the placement of the pocket opening and the presence or absence of C-terminal elements (Figure 1.C). We then convert each choice of high-level parameters into structure blueprint/constraints pairs (hereon referred to simply as blueprints), which guide backbone structure sampling at successive stages of fold assembly (next paragraphs; Figure 1.B). In total, there are 18 high-level fold parameters (Table S1), and each unique combination gives rise to a specific blueprint. At the lower level, backbone structures are generated according to these blueprints through Monte Carlo fragment assembly; the blueprints dictate the secondary structure and torsion angle bins of the fragments, as well as a number of key residue-residue distances (Fig. S1-4). In a final sequence design step, for each generated backbone, low energy sequences are identified through combinatorial sequence optimization using RosettaDesign.

We generate structure blueprints from the high level parameters using a hierarchical approach (Figure 1.B). First, the four main strands of the sheet are constructed, then helix 3 and the frontal hairpin, finally, the two N-terminal helices. If the backbone to be assembled has a C-terminal helix, it is added in a fourth step.

In the first step, the length and curvature of the sheet are the primary high-level parameters sampled (Fig 1C, top two rows). For each choice of high-level sheet length and curvature parameters, compatible sets of low-level parameters - secondary structure strings and angle and distance constraints - are generated to guide Rosetta fragment assembly. The translation from sheet length to secondary structure length is straightforward as longer strands generate longer sheets. To realize a specified sheet curvature, bulges are placed at specific positions on the edge strands, where they promote sheet bending (5, 15, 16). Bulges are specified by a residue with α-helical ɸ/ψ torsion values in the blueprint, leading to a backbone protuberance with two adjacent residues pointing in the same direction. As shown in figure 1A, there are always at least two bulges on the NTF2 sheet, delimiting the base and arms, and marking the axis around which the sheet curves. An additional bulge can exist on the long arm, further curving the sheet. To control the degree of curvature centered at the bending points, angle constraints are placed on C_α_ carbons on center strands, at positions adjacent to bulges (Fig. S1). Not all combinations of sheet length and curvature values are compatible with a closed pocket-containing structure, for example, long sheets with low curvature can not generate a cone-shaped structure. These incompatibilities are identified by attempting to directly construct sheet structures (as described above) across the full parameter space and then assessing the success in generating pocket containing structures; the region not sampled at the bottom left of Figure 3A reflects the incompatibility of long sheets and low curvature with the formation of a pocket (See Table S2 for the complete set of rules dictating high-level parameter combinations).

The range of possibilities for helix 3 and the frontal hairpin, which are generated next, is limited by the geometric properties of the sheet constructed in the first step. In order to determine which parameter combinations lead to folded proteins, we generated and evaluated backbone structures based on a wide variety of parameter combinations, and extracted the following rules, which are implemented in the generative algorithm. Sheets where the long arm does not protrude outwards over S6 require longer H3-S3 connections to place H3, S1 and S2 such that the correct strand pairing is realized (Fig. S2A). Conversely, sheets where the long arm protrudes outwards over S6 require shorter H3-S3 connections to avoid placing H3 too far from the rest of the structure (Fig. S2A). The length of H3 (possible lengths in residues: 10, 11, 14 and 15) is coupled with the torsional angles of the H3-S3 connection as the rigid S2-H3 connection limits the angle at which the last amino acid of H3 faces S3: lengths 10 and 14 are only paired with H3-S3 connections starting with a “B” torsion bin, and lengths 11 and 15 only with connections starting with a “G” torsion bin (Fig. S2B, Table S3). Independent from the H3 length and connection to the sheet, the length of the frontal hairpin strands has two possible values: 4 or 6 residues, with only 4 residues strands possible in narrow sheets (base length = 3), as S1 needs to be fully paired with S6 (Fig. S2C).

Stage 3, construction of the N-terminal helices, is likewise constrained by the geometric properties of the structure built so far. If the distance between the bulge on S6 and the H3-S3 connection is more than 25Å, then H1 and H2 are elongated by a full turn (4 amino acids) to close the cone described by the sheet (Fig. S3). The constraints that control the placement of H1 and H2 are adapted based on the shape of the current structure in order to position H1 and 2 such that good side chain packing is favored during sequence design, and occluding backbone polar atoms on the outward-facing edge of S3 is avoided (Fig. S3).

In cases where the backbone to be assembled has a C-terminal helix (has_cHelix = True), if the pocket opening is, like in most native NTF2-like proteins, between the frontal hairpin and H3 (Opening = Classic), the C-terminal helix is set to 8 residues long and rests against the long arm. If the opening is set to be between the termini of H1 and 2, and H3 (Opening = Alternative), then the C-terminal helix length is set to 11 residues long, and closes the space between H3 and the frontal hairpin (Figure 1B and S4).

### High-throughput characterization of the known de novo NTF2 structure space

The design of large pockets in *de novo* NTF2-like proteins is challenging and requires strategies to compensate for the loss of stabilizing core residues that would otherwise fill the space occupied by the pocket. Before setting out to experimentally sample the full range of structure space accessible to the generative algorithm, we chose to characterize the sequence and structure determinants of stability in the region of NTF2 space explored in our previous work (5), and its immediate vicinity. We generated 2709 new NTF2-like proteins belonging to the blueprints previously described, plus a few variations (9 different blueprints, see Fig. S5 and Table S4). We adapted a high throughput stability screen based on folding-induced protease resistance on the yeast cell surface, originally developed for small (< 43 amino acid) domains (17) to the much larger (105-120 residues) NTF2-like protein family. This required optimizing current methods (18) for efficiently splicing long oligonucleotides (230 bs) from oligonucleotide arrays to form longer genes by limiting pairing promiscuity and, therefore, the number of chimeric design combinations (see Methods).

A fifth (578, 21%) of the tested designs were stable (stability scores above 1), while only 2% of scrambled controls (randomly selected design sequences scrambled such that the hydrophobicity pattern is maintained) passed this stability threshold (Figure 2A). All tested blueprints had representatives among the stable sequences (Fig. S6). Analysis of the sequences and structures of the stable designs revealed several broad trends. There was a marked depletion of hydrophilic residues in positions oriented towards the protein core (Fig. S7), suggesting that the stable proteins identified in this first round experiment are likely folded as modeled, but may not be able to accommodate a pocket with polar amino acids, limiting their potential to be designed for general function. A logistic regression model trained to distinguish designs with stability score above or below 1.0 identified total sequence hydrophobicity (see “hydrophobicity” feature definition in SI methods), Rosetta energy (“score_res_betacart”) and local sequence-structure agreement (fragment quality, see “avAll”) as key determinants of stability (Figure 2.A and S8).

**Figure 2.**
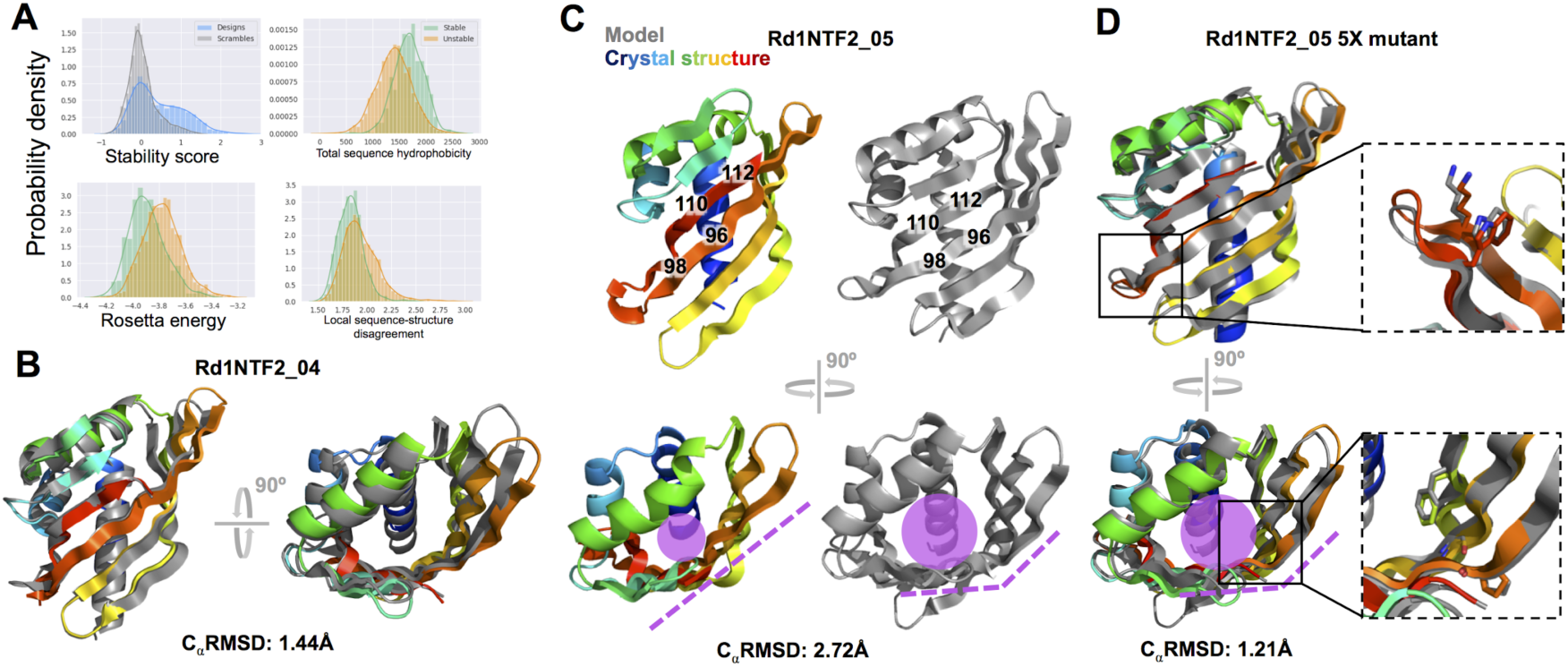
High-throughput screening and structural characterization of *de novo* NTF2-like proteins. **A:** (top left) Round 1 stability score distribution. Designs are more likely than scrambled sequences to have stability scores above 1.0. Remaining panels compare the distributions of sequence hydrophobicity, Rosetta energy, and local sequence-structure compatibility of stable and unstable (stability score < 1.0) designs. **B:** Crystal structure and computational model of design Rd1NTF2_04 (PDB ID 6W3G); the protein backbone is in very close agreement. **C:** Crystal structure and model of design Rd1NTF2_05 (PDB ID 6W3D), showing significant differences between model and structure. Strands 5 and 6 are shifted 2 residues relative to each other (bold numbers, left), resulting in a smaller space in the concave side of the flattened sheet (magenta sphere and dashed line, right). **D:** Crystal structure and model of design Rd1NTF2_05 5-fold mutant (PDB ID 6W3F), showing agreement between model and structure for backbone and mutated side-chains. As in C, a magenta circle and lines show how the concave side and sheet curvature fold as designed.

The importance of overall hydrophobicity is in agreement with the observed per-position amino acid enrichments, and suggests the composition or size of the designed protein cores is suboptimal. While Rosetta optimizes local sequence-structure agreement at single positions (p_aa_pp and rama_prepro energy function terms (19)), overall secondary structure propensities depend on stretches of several residues and cannot be decomposed in pairwise or single body energies. The detection of local sequence-structure agreement as a feature of stable designs suggests the first round design protocol produces sequences with suboptimal local sequence-structure relationship. These observations provide the basis for improving design methods, leading to more stable proteins in subsequent *de novo* NTF2 libraries.

We selected 17 designs with a stability score above 1 for more thorough biophysical characterization (See SI Methods). Seven of these expressed solubly in *E coli*, and all seven of them were folded, most remained folded up to 95°C, and had 2-state unfolding transitions in guanidine hydrochloride denaturation experiments (Fig. S9, S10 and Table S5). The remaining 10 designs did not express, or formed higher-order oligomers (Table S5), indicating stability score values above those of most scrambles are no guarantee of soluble expression and folding in *E. coli* cytoplasm.

We obtained crystal structures for two of the above-mentioned hyperstable proteins, with *de novo* NTF2 blueprints not characterized before (Figure 2.B and C, S11.A). The crystal structure and model of design Rd1NTF2_04 are in close agreement both in terms of C_α_ atom positions and most core side-chain rotamers (Figure 2.B and S11.B). In contrast, the structure of design Rd1NTF2_05 shows a two-residue register shift between strands 5 and 6 relative to the design (Figure 2.C), which results in a flatter sheet and a smaller core, a shorter strand 5, and longer strand 6. While the overall shape of the structure and the relative orientations of the hydrophobic residues in strand 5 and 6 are preserved (Figure 2.C), the structure deviations would be significant for a designed functional pocket. The identification of a design that is stable but has a structure different from its model provides an opportunity to discover determinants of structural specificity not captured by the design method.

We hypothesized that the disagreement between model and structure for design Rd1NTF2_05 originates from a lack of core interactions favoring the modeled high sheet curvature around residue 94, as well as from lack of consideration of negative design in the sequence choice for the 5-6 strand hairpin, which allows the shortening of strand 5. We identified several mutations that could favor the modeled sheet curvature and strand register. Mutations D101K and L106W near the strand 5-6 connection make favorable interactions in the context of the designed conformation, and replace leucine 106 by a large tryptophan side-chain, which would not fit in the context of the observed crystal structure (Fig. S12). Mutation A80G, at the most curved position of strand 4, favors bending by removing steric hindrance between the alanine 80 side-chain and the backbone at position 66, but leaves a void in the core, which modeling suggests should be rescued by I64F (Fig. S12, (6, 20)). A phenylalanine side-chain at position 64 makes favorable interactions in the designed conformation, and is likely to not fit in the core and be exposed in the observed conformation. Finally, the rigidity imparted by proline in position 94 limits the Ramachandran angles to those compatible with the designed conformation, as well as preventing strand 5 and 6 pairing beyond residue 92.

Experimental characterization of the Rd1NTF2_05 design 5-fold mutant showed a higher ΔG of unfolding than the original design (Fig. S13), and its crystal structure is in close agreement with the model (Figure 2.D). The side-chains at the five mutated positions were in the exact designed conformation, supporting our structural hypothesis and the incorporation of negative design to increase structural specificity (Figure 2.D, right). The 5-fold mutant also displays a large cavity, present in the design, the first example of a *de novo* designed monomeric NTF2 with a large pocket that does not require additional stabilizing features such as a disulfide bond or a dimeric interface (Fig. S14). The rationale used to change the structure of design Rd1NTF2_05 can be widely applied to ensure subsequent designs fold as modeled.

### High-throughput characterization of new regions of structure space explored by the generative algorithm

Armed with the insights from high-throughput characterization of known *de novo* NTF2 structural space, we set out to design proteins from hundreds of backbone blueprints created using the generative algorithm that explore a much larger structure space. We incorporated the lessons learned in the sequence design stage, with the goal of generating more stable and diverse designs that fold as modeled. To address the low sequence hydrophobicity, we added an amino acid composition term to the Rosetta energy function to favor sequences with 30% non-alanine hydrophobic amino acids on average, with different hydrophobicity targets for core, interface and surface positions. We also increased amino acid sampling to increase sequence-structure agreement (Fig. S15). Finally, guided by the experience with design Rd1NTF2_05, we incorporated steps in the design process that detect strand curvature ranges that require glycine placement to reduce strain. We used this improved method to generate a second round of designs exploring a much larger set of 1503 blueprints. These designs span a wide range of pocket volumes that are modulated by sheet length and curvature (Figure 3A x and y axes). There are two main modes by which the specification of sheet structure by the high level parameters modulates pocket volume. First, as sheets of similar length curve, the concave face collapses resulting in smaller pocket volumes (Fig. S16). Second, as sheets with similar curvature elongate, they wrap around the concave face and extend the pocket outwards (Fig. S16).

**Figure 3:**
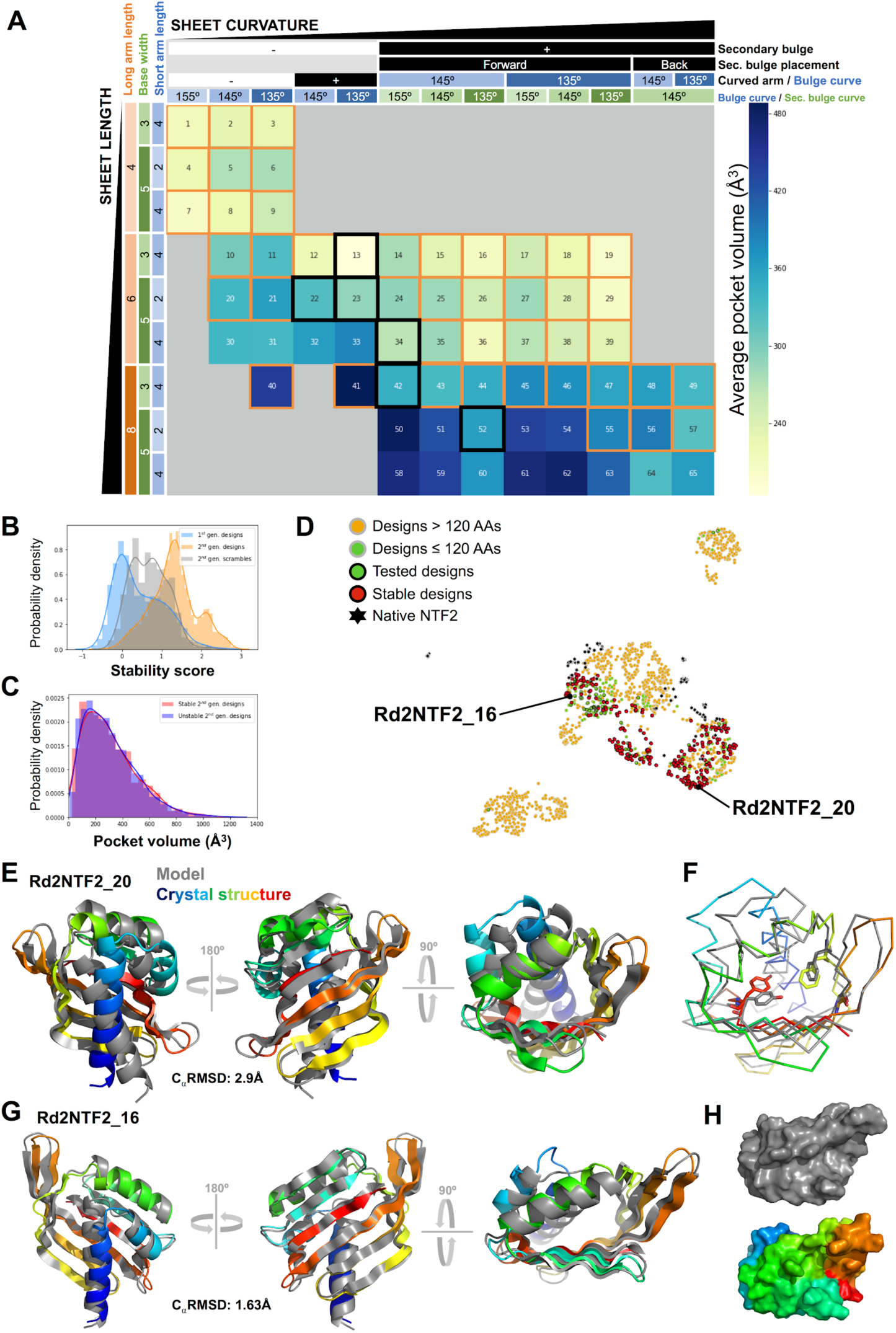
Biochemical characterization of round2 designs. **A:** *De novo* NTF2 designs sorted by sheet structure and ordered by sheet curvature and length. Each quadrant is colored by the average pocket volume of designs belonging to it. Orange frames denote quadrants for which stable designs were identified. Black frames denote designs were tested, but no stable design was identified. **B:** Stability score of algorithm designs (orange), compared to controls (grey) and designs from the initial screening (blue). **C:** Volume distribution of stable and unstable designs. **D:** UMAP embedding of NTF2 structural space, using 1-TMscore as pairwise distances. Each dot is a single structure. In the case of *de novo* designs, each unique parameter combination is represented by a single structure randomly selected from that combination. Structurally similar proteins are closer together. **E:** Crystal structure of stable design Rd2NTF2_20 (PDB ID 6W3W), which features a new, elongated helix 3-strand 3 connection. Despite significant differences between the model and structure in the N-terminal helices, the new loop and the sheet are well recapitulated. **F:** Core rotamers of Rd2NTF2_20. TYR101 (red, sticks) shows a significant deviation from the model, and enables the change in location of helix 1. In contrast, PHE61 and GLY77 interact as modeled, showing the glycine rescue feature can be designed from scratch. **G:** Crystal structure of stable design Rd2NTF2_16 (PDB ID 6W40), which features a secondary bulge and an elongated frontal hairpin, features not designed before. Both of these features are recapitulated in the crystal structure. As in Rd2NTF2_20, but not as dramatic, the Rd2NTF2_16 crystal structure presents significant deviations from the model in the N-terminal helices. **H:** Surface rendering of the model and crystal structure of Rd2NTF2_16, showing the shallow pocket formed by the long arm and the frontal hairpin is recapitulated by the crystal structure.

Due to gene length limitations, we were able to test designs for 323 unique parameter combinations out of the possible 1503 -- these yield proteins of 120 amino acids or less in length. We synthesized genes for 5188 proteins generated from these 323 blueprints, and subjected the designed proteins and scrambled versions to the protease stability screen. The protease resistance of the scrambled sequences was greater than in the first high-throughout experiment, likely due to the increased sequence hydrophobicity (Fig. S17). Roughly one third (29%) of the designs had stability values above those of most scrambled sequences (Figure 3B, 98% of all scrambles have stability score <1.55), a larger fraction than the 21% of stable designs in the initial screen, increasing our dataset of stable NTF2-like designs from a total of 578 to 2077. These stable designs belong to 236 parameter combinations, a very large increase over the 9 combinations in the previous round, with most of the missing combinations having less than 10 initial samples (Fig. S18). The new parameter combinations include structural features not sampled before, such as a secondary bulge on the long arm, new H3-S3 connections and elongated frontal hairpins. The pocket volume distribution of stable designs is very similar to the distribution for all tested designs (Figure 3C), suggesting that pocket volume is not a limiting factor, and spans most of the native NTF2 range (Fig. S19). Furthermore, per-position amino acid identities in stable designs show much lower levels of general enrichment and depletion than those in the first round of high-throughput screening (Fig. S20). In particular, polar amino acids are not depleted in core positions (Fig. S20), suggesting that polar residues are likely better tolerated in pocket positions, perhaps due to the improved core packing resulting from the optimized sequence design protocol.

With the large increase in diversity in the second round, the stable designs created by the generative algorithm span a very wide range of structures. To visualize the space spanned by our generated structures compared to native NTF2 structures, we used the UMAP algorithm (21) to project similarity in backbone structure (TM-score, (22)) into two dimensions (Figure 3D, see Fig. S21 for map generated using different UMAP hyperparameters). The grouping of structures with similar features in different map regions provides an indication of which generation parameters lead to novel NTF2 structures (Fig. S22). Inspection of the map shows that our algorithm samples most of the native space, as well as completely uncharted regions. The subset of designs tested by high-throughput screening sample a wide range of structures within the accessible protein length, and stable representatives from the 236 unique NTF2 parameter combinations are found across the sampled space (Figure 3A and D). Overall, the number and diversity of *de novo* designed NTF2-like structures is considerably larger than that of the NTF2 structures in the PDB. Native structures appear in small clumps in NTF2 space, as they fall into groups with highly similar members. In contrast, *de novo* NTF2-like proteins sample large areas more uniformly, providing fine-grained sampling of the structural space, and hence more optimal starting points for designing new functions requiring new pocket geometries. Most native proteins are close to *de novo* groups, reflecting overall structural similarity, but are peripheral to them. This structural distinction likely reflects differences in loop structure: native NTF2-like proteins often have long loops, but our designs tend to have short loops.

A logistic regression model trained on stability of second-round designs suggests the lessons from the first round of high-throughput screening proved effective, and provides new suggestions for improvement (See SI text and Fig. S23). Furthermore, features based on the high-level parameters of the generative algorithm (e.g., H3 length, sheet curvatures, sheet length and hairpin length) did not contribute significantly to stability prediction, suggesting stable proteins can be designed across all the considered structural space (See SI text).

We biochemically characterized 37 stable designs from the second round of high-throughput screening. Less than half of them (43%, similar to the 41% in round 1) expressed solubly in *E. coli* and were folded. Most of these folded designs remained folded above 95°C (Fig. S24-25 and Table S6). The length of helix 3 in two of these second-round stable designs, Rd2NTF2_06 and Rd2NTF2_19, is the longest of the values we allowed, supporting the designability of this feature despite it being slightly disfavored by the stability model (SI text, S23). The remaining 20 second-round designs did not express, and a few formed higher-order oligomers (Table S6).

More than half of the designs we attempted to express in *E. coli* did not express or formed soluble aggregates, indicating high stability score does not necessarily translate to folding in *E. coli* cytoplasm. While stability score has no significant correlation to ΔG_unfolding_ for these larger proteins, it has some capacity to discriminate between designs that fold from those that do not (Fig. S26C). Furthermore, 9 out of the 9 designs with low stability score we attempted to express in *E. coli* did not fold, supporting the use of stability score as a metric to improve the design of these pocket-containing proteins. In an attempt to improve the power of the stability score to predict folding and stability of proteins expressed in *E. coli*, we trained an alternative unfolded state protease resistance model based on the protease resistance of scrambled sequences (see SI Methods and Fig. S26D-E). As expected, this model predicts NTF2 scrambled sequence stability better than the published unfolded-state model, but using it to recalculate stability scores does not lead to better prediction of ΔG_unfolding_ or folding in *E. coli* (Fig. S26B,C).

For two of the folded and hyperstable designs (Rd2NTF2_20 and Rd2NTF2_16), we obtained high-resolution crystal structures, and found that they are in close agreement with the models (Figure 3E-H). Both designs feature structural elements designed for the first time in *de novo* NTF2-like proteins by the generative algorithm. Rd2NTF2_20 has an extended connection between H3 and S3, recapitulated in the crystal structure (Figure 3E), which enables the use of a short helix 3. Rd2NTF2_16 features two new structural elements, a bulge on the long arm (in addition to the ones flanking the base), and an extended frontal hairpin, both recapitulated in the crystal structure (Figure 3G). The additional bulge enables higher curvature on the long arm, contributing significant diversity to long arm structure, which can be further increased by allowing different bulge placements. The extended hairpin, which is only designable when the base is long enough, extends the pocket outwards, thereby increasing its volume. In the case of Rd2NTF2_16, the combination of these features yields a protein with a shallow groove instead of a pocket (Figure 3H); the ability to generate proteins with shallow grooves with two open ends should enable design of binding sites for polymers such as peptides or polysaccharides.

The accuracy of the Rd2NTF2_20 and Rd2NTF2_16 computational models indicated by their close agreement with the experimental crystal structures follows directly from the insights gained in the first large scale design round. Both proteins feature a glycine on strand 4, enabling high curvature between the base and the long arm, as described for the design Rd1NTF2_05 5-fold mutant, and consequently incorporated in the generative algorithm. In order to implement the glycine placement on strand 4 as generally as possible, the design protocol searches for large hydrophobic side chains to fill the void left by the glycine. In Rd2NTF2_20, this is achieved by a phenylalanine in the same conformation as the one observed in the design 0589 5-fold mutant, while in Rd2NTF2_16 a void is left in the core. Unlike design 0589, in the Rd2NTF2_20 and Rd2NTF2_16 crystal structures the highly curved sheet conformation is in close agreement with the model. In addition to generally supporting the models created by the generative algorithm, the two crystal structures provide information to improve the design method (See SI text and Fig. S27). The ability to design and properly model the sheet in *de novo* NTF2-like proteins is of great importance, as this structural element is the most involved in pocket structure.

Most of the 1503 possible high-level parameter combinations yield proteins that are too long to be encoded by assembling two 240 base-pair oligonucleotides (the current limit in what can be synthesized at very large scale). To explore the parameter space that generates these longer proteins, we characterized 10 designs that are predicted to be stable by a logistic regression model trained on the second high-throughput screening experiment data, and have large pockets (500 to 1200Å^3^). Two of the ten were monomeric and remained folded above 95°C, a success rate similar to that of the biochemical characterization of designs identified in the second high-throughput experiment, suggesting that *de novo* NTF2-like proteins longer than 120 amino acids with large pockets are also designable using the generative algorithm (Fig. S28-S29, Table S7).

### Suitability of designed scaffolds for harboring small molecule binding sites

To probe the capability of the designed proteins to host binding sites, we docked 50 ligands (See Fig. S30 and Methods) from the PDB in all *de novo* and native NTF2 structures with pockets larger than 30Å^3^ (862 and 64, respectively), and optimized the surrounding sequence to interact with the ligand. We then evaluated whether the pocket with the most favorable interactions was based on a *de novo* or native protein, for each ligand. This test provides a conservative estimate of the relative ability of the designs to scaffold binding sites, as they were not constructed to bind any specific molecule, we only used the subset of stable *de novo* proteins that already had a pocket, and limited design to positions within that pocket. Despite these disadvantages, *de novo* proteins provide a better (lower ligand interaction energy) pocket for 80% of all tested ligands (40 out of 50), without obvious biases in ligand molecular weight, charge, chemical groups, or hydrophobicity (Figure 4 and S31). The *de novo* scaffold with the largest number of top ranking docks is Rd2NTF2_03, one of the designs found to be folded and highly stable (Fig. S32). As controls for this docking test, we included two small molecules in the ligand set that are bound by the native scaffolds (PDB ligand codes EQU and AKV, bound by 1OH0 and 2F99 respectively) and found that native-like poses are recovered when the bound ligand conformer found in the crystal structure is used (Fig. S33). The observed advantage in binding site scaffolding should increase with the number of *de novo* designed structures generated, while the rate of growth of the native set is limited to what has been sampled by evolution.

**Figure 4:**
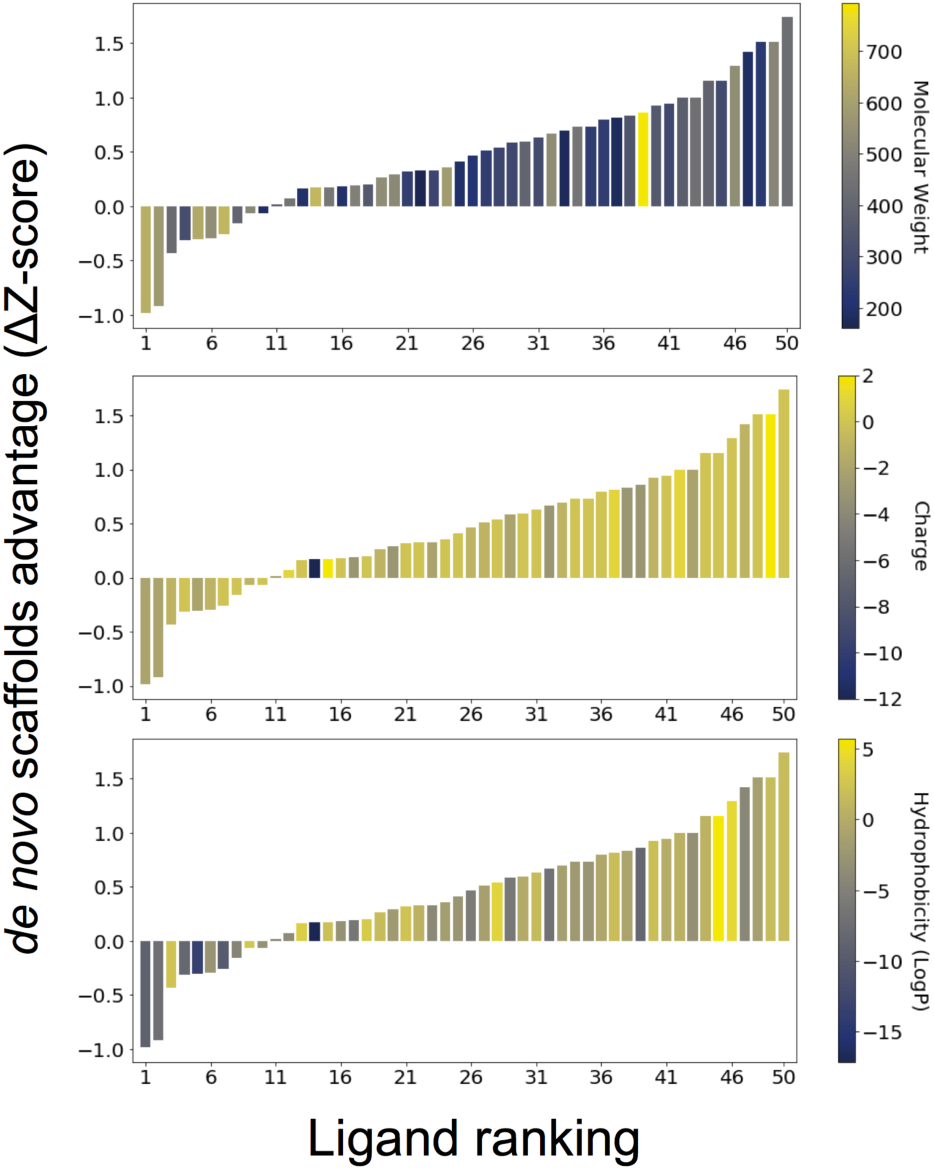
Comparison of *de novo* designs to native structures for ligand docking and design. Ligands are ranked by how well *de novo* scaffolds accommodate each of them in comparison to native structures (higher ΔZ-score values mean higher advantage of best *de novo* scaffold over best native). To calculate ΔZ-score for each ligand, we obtain the protein-ligand interaction energy for all docked conformations of that ligand, and calculate the Z-scores, then subtract the Z-score of the lowest (most favorable) *de novo* designed protein dock, from the lowest native protein dock. In each panel the same ranking is colored by different ligand properties: from top to bottom: Molecular weight (Da), charge at pH 7.5, and hydrophobicity (LogP).

As the overarching goal of this work is to expand the set of available protein structures with pockets, we generated a final set of scaffolds that incorporates all the lessons from previous experiments. Improvements in the generative algorithm, both in sequence design and backbone generation resulted in increased diversity (1619 unique parameter combinations) and improved stability-related metrics (see Fig. S34 and SI methods). We have made this set of 32380 scaffolds (20 models with different sequences per parameter combination) available for general use as starting points for ligand binding and enzyme design.

## Discussion

Our generative algorithm may be viewed as encoding the “platonic ideal” of the NTF2-like structural superfamily along with a method for essentially unlimited sampling structures belonging to it, in a fashion directly tied to pocket structure. In terms of SCOPe categories, each combination of top-level parameters can be thought of as a protein family, and the set of all combinations, the *de novo* NTF2-like structural superfamily. Whereas in our previous work 4 NTF2 structure blueprints were manually constructed, the new generative algorithm samples through over 1600 unique blueprints that result in well-formed backbones. This represents a qualitative jump in the structural diversity that can be achieved for complex folds by *de novo* protein design. The generative approaches to *de novo* protein structure design so far described in the literature, rule- or model-based, either focus exclusively on helical structures (23–25), are not geared towards atomic-detail modeling and design (26), or sacrifice fine-grained structural control for structural diversity (27). Machine-learning based generative models show considerable promise (27, 28), but have not yet been applied to the direct generation of full atomic structures with specific features of interest, as we do here for scaffolds containing a varied geometry of binding pockets.

Up to now, protein design for a specific function has relied either on searching through the scaffolds in the PDB, or generating small variations of a limited set of *de novo* scaffolds. Our approach now enables going far beyond both approaches by searching through an essentially unlimited set of generated scaffolds. The experimental characterization of the designs shows that the generative algorithm samples a wide range of feasible structural space, and that designs usually fold as modeled. The insights we gained in order to produce these diverse proteins can be harnessed to improve the success rate in future protein design efforts. Furthermore, our approach could be implemented for other protein folds to expand structural diversity even further. In combination with existing docking and design methods, the generative algorithm here presented should open the door to design of novel functions by eliminating the limitations imposed by current protein structural databases, and enabling scaffold generation custom-tailored to function.

## Materials and Methods

### Generative algorithm for proteins from the NTF2-like superfamily

All code can be downloaded from GitHub at: https://github.com/basantab/NTF2Gen

The NTF2Gen repository contains all the tools for *de novo* design of NTF2-like proteins. The main script is CreateBeNTF2_backbone.py, which manages the construction of NTF2 backbones, followed by DesignBeNTF2.py, which designs sequence on a given backbone generated by the previous script. To generate backbones from a specific set of parameters, use CreateBeNTF2PDBFromDict.py. The fundamental building blocks of the backbone generation protocol are Rosetta XML protocols (included in the repository) that are specialized instances of the BlueprintBDRMover Rosetta fragment assembly mover. All checks and filters mentioned in the result section previous to design are implemented either in the XML files or the python scripts. Additional backbone quality controls are ran after each step (See SI methods). The design script is also based on a set of XML protocols, one for each of the described stages. The glycine placement in highly curved strand positions and the selection of pocket positions are managed by DesignBeNTF2.py (BeNTF2seq/Nonbinding). Pocket positions are selected by placing a virtual atom in the midpoint between the H3-S3 connection and the S6 bulge, and choosing all positions whose C_α_-C_β_ vector is pointing towards the virtual atom (the V_atom_-C_α_-C_β_ angle is smaller than 90°), and their C_α_ is closer than 8Å.

### De novo NTF2 backbone generation and sequence design for the first round of high-throughput screening

Backbones were constructed as described in (5). For families not described in said paper (i.e., BBM2nHm* designs), the same backbone construction algorithms were used, but parameters were changed accordingly. Scripts for producing all these backbones can be found at https://github.com/basantab/NTF2Analysis, NewSubfamiliesGeneration. The sequence design protocol for the first round of designs can be found in the above-mentioned GitHub repository. Briefly: The design protocol begins by generating 4 different possible sequences using the Rosetta FastDesign mover in core, interface and surface layers separately. Then random mutations are tested, accepting only those that improve secondary structure prediction without worsening score, introducing Ramachandran outliers or worsening the shape complementarity between helices and the rest of the protein.

### Design of gene fragments for multiplex gene assembly

In order to obtain full-length genes from fragments synthetized in DNA microarrays, they must be assembled from halves, as described in (18). To generate highly orthogonal overlaps, we generated DNA sequences using DNAWorks (29), then split the gene in half and altered the composition of the around 20 overlapping nucleotides to have as low homology as possible with other halves in the pool, while maintaining an adequate melting temperature and GC content, and staying below the maximum oligonucleotide length (230 nucleotides). An optimized version of the algorithm described in (18) can be found at https://github.com/basantab/OligoOverlapOpt.

### Protease-based high-throughput stability screening

The protease-based high-throughput stability screening was carried out as described in (17). Briefly: genes encoding for thousands of different *de novo* NTF2 sequences cloned in the pETCON2 vector, which has the protein of interest expressed as a chimera of the extracellular wall yeast protein AgaII, on its C-terminus, connected by a “GS” linker of alternating glycine and serine. The protein of interest is followed by a myc-tag (EQKLISEEDL). This library is transformed in yeast for surface display in a one-pot fashion using electroporation. Different aliquots of the yeast culture are then subject to increasing concentrations of trypsin and chymotrypsin, and labeled with an anti-myc tag antibody conjugated to fluorescein. Cells still displaying full proteins (myc-tag-labeled) after this treatment are then isolated by Fluorescence-Activated Cell Sorting (FACS). Deep-sequencing of the sorted populations reveals which sequences are protease resistant and to what degree, providing an estimate for folding free energy. The metric reported by this assay is the stability score, an estimate of how much protease is necessary to degrade a protein over that expected if the protein was completely unfolded. A stability score of 0 indicates that the protein is degraded by the same amount of protease as expected if it was unfolded, i.e., it is likely completely unfolded. A stability score of 1 indicates that 10 times more protease is required to degrade the protein, than expected if it was completely unfolded.

### LASSO logistic regression model training on stability data

To identify features that predict stability, we trained LASSO logistic regression models (30) using the features described in the previous section, after normalization. A logistic regression model predicts the probability of a binary outcome using a logistic function that depends on a weighted summation of features. By sampling a series of L1 regularization values, we obtain models with varying degrees of parsimony, and for each of those L1 values we also generate different random partitions of our dataset. This way, for each L1 value we obtain models with a spread on accuracy, which we use for selecting an L1 regularization value that maximizes accuracy and minimizes complexity - i.e., the number of features with weight different from 0. The simplest measure of the importance of each feature is its assigned coefficient.

The data and code for analysis of data derived from the first high-throughput experiment can be found at:

https://github.com/basantab/NTF2analysis, ProteaseAnalysisExp1/LassoLogisticRegression.ipynb

Analysis of data from the second high-throughput experiment can be found at:

https://github.com/basantab/NTF2analysis, ProteaseAnalysisExp2/LassoLogisticRegression_new_version.ipynb

### Crystallography data collection and analysis metrics

To prepare protein samples for X-ray crystallography, the buffer of choice was 25 mM Tris, 50 mM NaCl, pH 8.0. Proteins were expressed from pET29b+ constructs to cleave the 6xHis tag with TeV. Proteins were incubated with TeV (1:100 dilution) overnight at room temperature and cleaved samples were loaded to a Ni-NTA column pre-equilibrated in PBS+30mM Imidazole. Flow-through was collected and washed with 1-2 column volumes. Proteins were further purified by FPLC as described above and specific cleavage of the 6xHis tag was verified by SDS-PAGE.

Purified proteins were concentrated to approximately 10-20 mg/ml for screening crystallization conditions. Commercially available crystallization screens were tested in 96-well sitting or hanging drops with different protein:precipitant ratios (1:1, 1:2 and 2:1) using a mosquito robot. When possible, initial crystal hits were grown in larger 24-well hanging drops. Obtained crystals were flash-frozen in liquid nitrogen. X-ray diffraction data sets were collected at the Advanced Light Source (ALS). Crystal structures were solved by molecular replacement with Phaser (31) using the design models as the initial search models. The structures were built and refined using Phenix (32, 33) and Coot (34). Crystallization conditions and data collection and refinement statistics can be found in the SI methods and Table S16.

### UMAP embedding of NTF2 designs

Uniform Manifold Approximation and Projection (UMAP) (21) is a dimension reduction technique widely used for visualization of high-dimensional data. We obtained the code for running UMAP by following instructions in https://umap-learn.readthedocs.io/en/latest/. For generating the embedding, UMAP requires a distance measure between points, for which we provided 1-TMscore between all samples. We ran UMAP in a Jupyter notebook with different metaparameter combinations and verified that the general cluster structure was conserved among all of them, and that structural features were reflected in the groupings. The code and files necessary for generating the UMAP-related figures can be found in the GitHub repository https://github.com/basantab/NTF2analysis, UMAP_embedding.

### Ligand in silico docking test

The goal of the ligand *in silico* docking test is to provide an estimate of how *de novo* NTF2-like proteins compare to native ones in terms of their ability to harbor arbitrary binding sites. We used RIFDOCK (6) for simultaneous docking and design based on a set *de novo* and native protein backbones. As RIFDOCK only uses backbone coordinates and a list of pocket positions to dock the ligand and design a binding site around it, it can be used in a sequence-agnostic way. We selected and prepared (see ligand preparation methods above) a subset of 50 ligands from all non-polymeric PDB ligands (Ligand Expo - http://ligand-expo.rcsb.org) using k-means clustering on physical and chemical features (See S30, and the 50_ligand_table.html file at https://github.com/basantab/NTF2analysis/tree/master/ligandInSilicoDockingTest). The number of ligands tested was limited to 50 for computational tractability, as RIFDOCK uses a significant amount of resources per ligand and scaffold: >3hs in 32 cores and 64GB of RAM on average per ligand, to generate the initial rotamer interaction field (RIF), and ∼2hs in 32 cores using >20GB of RAM, per ligand for docking in a subset of 12 scaffolds. As NTF2-like native representatives, we selected 64 structures with pockets (pockets detected and defined as described above) from the SCOPe2.05 database (described above). In order to provide a conservative estimate of pocket diversity and aid computational tractability, we limited the set of *de novo* designs used for docking to those stable (stability score > 1.55) and with detectable pockets in the concave side of the sheet (>25% overlap between CLIPPERS-detected pocket and backbone-based pocket sets, and >30Å^3^ volume), resulting in 862 different *de novo* sequences (See https://github.com/basantab/NTF2analysis “ligandInSilicoDockingTest” for relevant files). Pocket residues were detected using CLIPPERS, as described above, and only positions originally lining the pocket of the scaffolds were considered for binding site design by RIFDOCK. We designed five designs per scaffold, per ligand, and sorted them by “packscore”, a measure of favorable Van der Waals interactions and hydrogen bonds, with bonuses for bidentate (one side chain contacting two hydrogen-bonding ligand atoms) interactions. We measured the capacity of de novo scaffolds to accommodate binding sites batter by natives by subtracting the best native packscore Z-score from the best *de novo* packscore Z-score.

### Data and code availability

In order to facilitate reproducibility, improvement, further analysis and use of the models and information in this work, we have made all relevant data and code publicly available on basantab/NTF2Analysis and basantab/NTF2Gen GitHub repositories. All sequences, PDB models, analysis scripts and data tables for the first high-throughput experiments can be found in the ProteaseAnalysisExp1 folder of NTF2Analysis, and ProteaseAnalysisExp2 for the second high-throughput experiment. The set of 32380 scaffolds, available for general use as starting points for ligand binding and enzyme design, is available in the BeNTF2seq/design_with_PSSM/final_set folder in the basantab/NTF2Gen GitHub repository.

## Author Contributions

B.B. and D.B. designed research; B.B., M.J.B., A.K.B., C.N., C.M.C., L.P.C. and F.D. analyzed data; B.B., C.N. and D.B. wrote the paper; and B.B., C.M.C., L.P.C. and I.G. performed experiments.

## Acknowledgments

We thank current and former Baker laboratory members, in particular, Anastassia Vorobieva, Gabriel Rocklin, Mark Lajoie, Enrique Marcos, Ivan Anishchanka, Brian Coventry, William Sheffler, Hahnbeom Park and Sergey Ovchinnikov for insightful conversations and suggestions. This work was funded by DARPA Synergistic Discovery and Design (SD2) HR0011835403 contract FA8750-17-C-0219, Eric and Wendy Schmidt by recommendation of the Schmidt Futures program and the IPD Directors Fund. CN is supported by Novo Nordisk Foundation grant NNF17OC0030446. This work was facilitated through the use of advanced computational, storage, and networking infrastructure provided by the Hyak supercomputer system at the University of Washington.

## Supplementary Information Text

### Logistic regression model of stability trained on the second round of high-throughput experiments

As the proteins designed by the generative algorithm sample uncharted structural space, we investigated whether there were substantial differences in stability in different regions of the space. We included the high-level parameters used in the generative algorithm in a logistic regression model of stability (designs with stability score >1.55 are labeled as stable), and found that most have zero weight, suggesting that proteins can be equally well designed throughout structural space. The only high-level parameter feature with a non-zero weight is the length of helix 3; the negative weight indicates that designs with shorter helices tend to be more stable (the relatively small weight of this feature suggests that this is only a small bias; Fig. S23). The logistic regression model also detects a signal (positive weight) from core hydrophobic packing, but not raw sequence hydrophobicity (Fig. S23), suggesting that the contribution of overall sequence hydrophobicity was fully saturated going from the first round to second round of designs, and that the increase in the proportion of stable designs is not solely due to increased hydrophobicity. The model also shows a shift in importance from local sequence-structure compatibility to tertiary structure-sequence compatibility (Fig. S23), likely due to the extensive secondary structure propensity optimization in the second round of designs.

### Lessons from design Rd2NTF2_20 and Rd2NTF2_16 crystal structures

The N-terminal helices of both designs display significant deviations from the model. While this could be attributed to the extensive crystal contacts in both structures (Fig. S27), the alternative conformations of the side chains near the N-terminal helices suggest that better core packing could have prevented the backbone rearrangements. In the case of Rd2NTF2_20, four core side chains, N11, T75, T92, Y101 form a hydrogen bond network with an alternative conformation from the design, following the displacement of the helix. In particular, T92 is completely buried in the model, with a single polar interaction towards the backbone of residue 73, and it is possible that this interaction is not as favorable as the one with a water molecule observed in the crystal structure (Fig. S27). This highlights the importance of ensuring all polar interactions among buried side chains are highly favorable. A similar displacement of the N-terminal helix is observed in Rd2NTF2_16. In this case, the helix is again involved in crystal contacts, with significant deviations in core packing, but most side chains involved are hydrophobic. Both Rd2NTF2_20 and Rd2NTF2_16 feature glycines at the points of highest sheet curvature, but unlike Rd2NTF2_20, Rd2NTF2_16 has a cavity above it, as Rosetta was unable to find a favorable side-chain placement to fill it (Fig. S27). It is possible the destabilizing effect of this large void leads to the displacement of the N-terminal helix, which further illustrates the need of compensating the packing interactions lost by placing glycine on inward-facing strand positions.

## Supplementary methods

### De novo NTF2 backbone quality control

The generative algorithm includes quality control steps throughout the backbone construction process and sequence assignment to reduce deviation from ideal atomic geometry. At every stage, output that has Ramachandran outliers or unlikely bond angle and length values are discarded. Furthermore, sheets and helices where deformation leads to backbone hydrogen bonds with higher energy than average are discarded. The backbone assembly process is guided by constraints that impart the unique structural features for a given parameter combination, and structures falling beyond tolerable limits of some of those constraints are discarded. Finally, since the sequence assignment process can change the backbone slightly, filters are also applied at the end of this step, especially those that ensure a compact structure. All the previously mentioned filters are implemented as Rosetta filters within each Rosetta script (*.xml files).

### Quantification of de novo NTF2-like proteins families

As previously described, we define *de novo* NTF2-like families as all different combinations of high-level parameters that lead to well-formed backbones (See above, “*De novo* NTF2 backbone quality control”). To quantify the number of possible parameter combinations, we ran the backbone generation algorithm without requesting any specific combination, i.e., randomly choosing parameters at each stage, and obtaining only backbones passing all quality controls (See https://github.com/basantab/NTF2Gen, CreateBeNTF2_backbone.py). We then assign a sequence to the backbones. We quantify the number of different parameter combinations by reading the dictionary stored in each output, marked with the keyword “NTF2DICT” at the beginning of the line, which stores the parameters used to generate it (See https://github.com/basantab/NTF2Gen, PrintUniqueBeNTF2_file_input.py). The process of generating backbones from random parameter combinations leads to an uneven combination distribution – not all combinations are equally sampled, and some backbones are harder to assemble than others. To compensate for this, we take all combinations with a number of representatives lower than required, and generate more backbones with the same parameters (See https://github.com/basantab/NTF2Gen, CreateNewBeNTF2PDBFromDict.py), until the required number is met. For the final set of backbones, this number is 20.

### Pocket structure analysis using CLIPPERS

We created pocket inventories for each protein of interest using CLIPPERS (Coleman and Sharp, 2010) with default options. We then scanned through these inventories searching for the largest pockets using travel depth to define their boundaries: We trimmed the pocket tree (done by starting with the deepest, group=1, and walking back with through parents, capping it at group # 120) using a mean_TD cutoff defined as: pocket mean_TD ∼ max_TD - (max_TD - lowest_mean_TD) * X, with X = 0.75. The python code for this (pocketDetect_lines_TD_CLIPPERS.py) can be found at https://github.com/basantab/NTF2analysis. After detecting pockets, structures where the pocket was not in the canonical location (sheet concave side) or spanned micro-pockets on the surface, were discarded.

### Analysis of pocket volume in proteins designed by the generative algorithm

To produce the table in figure 3A, we measured the pocket volumes of all models generated in preparation for the second high-throughput experiment (including those finally tested) using CLIPPERS, as described above. Only proteins whose pockets could be detected by CLIPPERS were included in the analysis (13126 of 22853). See https://github.com/basantab/NTF2Analysis, ProteaseAnalysisExp2/Figure3_pocket_volume/Create_heatmap.ipynb. Figure 3C histograms were produced using the same data.

### Protein-protein alignment by TM-align

For each alignment, TM-align optimizes and reports TM-score, a measure of the distance between C_α_ carbons of aligned residues in target and template, normalized by protein length. The optimization algorithm used by TM-align results in alignments where superposition of segments with similar local structure is optimized over superposition of segments with disparate local structure. Because TM-score is normalized by target length, and we align proteins with similar, but not equal, lengths, for any given alignment, the TM-score we report is the average between two values.

### Generation of patterned scrambled sequences for control

In order to produce control sequences that retain the overall amino-acid composition, but are not optimized for folding, we took a subset of the design sequences, and scrambled all amino-acid identities, except for P and G, while keeping the hydrophobicity pattern. The code for this can be found in the GitHub repository https://github.com/basantab/NTF2analysis, create_patterned_scramble.py

### Features calculated for de novo NTF2 design stability prediction

Scripts for extracting design features used in logistic regression model training can be found in the public GitHub repository: https://github.com/basantab/NTF2analysis in the feature_extraction folder. For features described in (1) extracted with specialized code, refer to the supplementary material of that publication.

The features calculated using Rosetta filters and score function ref2015 (when dependent on score function) can be found on table S8. Features calculated using Rosetta filters and beta_nov16 score function (when dependent on score function) can be found in Table S9. Features calculated using Rosetta filters related to burial of unsatisfied polar atoms can be found in Table S10. Features calculated using CLIPPERS (Coleman and Sharp, 2010) pocket detection and inventory software can be found in Table S11.

Overall protein fragment metrics calculated for protein fragments with similar sequence and secondary structures to 9-mer sequence stretches (protein length-9) in the target protein in Table S12. For each 9-mer, 200 structure fragments are derived, as described in (2, 3).

Overall protein TERM metrics are calculated based on the output of the scripts provided with (4). TERM-based metrics were calculated based on the per-positions abundance_50, design_score_50 and structural score. These metrics can be found in Table S13. To obtain insight regarding specific parts of the proteins, we divided the protein in continuous sequence stretches that form local structures (sometimes with overlapping positions, Table S15), and calculated different fragment and TERM features in each of them, these can found in table S15. For each of the above stretches, TERM and fragment metrics were calculated, and the final name of the features calculated this way are <stretch name>_<metric>.

Features Tminus1_netq, Tend_netq, T1_absq, Tminus1_absq, Tend_absq, abego_res_profile, abego_res_profile_penalty, largest_hphob_cluster, n_hphob_clusters, hphob_sc_contacts, hphob_sc_degree, n_charged, hydrophobicity, contig_not_hp_internal_max, contig_not_hp_avg, contig_not_hp_avg_norm, tryp_cut_sites, chymo_cut_sites, chymo_with_LM_cut_sites, nearest_chymo_cut_to_Nterm, nearest_chymo_cut_to_Cterm, nearest_tryp_cut_to_Nterm, nearest_tryp_cut_to_Cterm, nearest_tryp_cut_to_term and nearest_chymo_cut_to_term, were calculated using the enchance_score_file.py script provided with (1), and are thoroughly explained in their supplementary materials.

### Hydrophobicity enrichment sequence profile

Designs were split between stable and unstable depending on the threshold selected for each experiment (see Results), and the enrichment was calculated based on the whole population frequencies vs. the frequencies in the stable population. Code for these calculations, figures and derivation of sequence data from designs on the first high-throughput experiment can be found at https://github.com/basantab/NTF2analysis, Exp1_SeqProfile

For designs tested on the second experiment: https://github.com/basantab/NTF2analysis, Exp2_SeqProfile.

### Experimental characterization of designs

#### Protein expression and purification in E. coli

Genes encoding the designed protein sequences were obtained from IDT already cloned in pET29b+ or pET21b+ (with N-terminal 6xHis tag followed by a TeV cut-site) expression vectors. Plasmids were transformed into chemically competent *Escherichia coli* Lemo21 cells from Invitrogen. Starter cultures were grown at 37°C in Luria-Bertani (LB) medium overnight with antibiotic (50 µg/ml carbenicillin for pET21b+ expression or 30 µg/ml kanamycin for pET-28b+ expression). For expression, overnight 5mL LB cultures were used to inoculate 500 mL of Auto-induction medium supplemented with antibiotic, at 25°C, for 18 hours (5). After overnight expression, cells were collected by centrifugation (at 4 °C and 4400 r.p.m for 10 minutes) and resuspended in 25 ml of lysis buffer (30 mM imidazole and phosphate buffered saline, PBS - 137 mM NaCl, 12 mM Phosphate, 2.7 mM KCl, pH 7.4).

Resuspended cells were lysed by sonication or microfluidizer in the presence of lysozyme, DNAse and protease inhibitors. Lysates were centrifuged at 4 °C and 20,000 r.c.f. for 30 minutes; and the supernatant was filtered and loaded to a nickel affinity gravity column pre-equilibrated in lysis buffer for purification. The column was washed with three column volumes of PBS+30 mM imidazole and the purified protein was eluted with three column volumes of PBS+300 mM imidazole. The eluted protein solution was dialyzed against PBS buffer overnight. The expression of purified proteins was assessed by SDS-polyacrylamide gel electrophoresis; and protein concentrations were determined from the absorbance at 280 nm measured on a NanoDrop spectrophotometer (ThermoScientific) with extinction coefficients predicted from the amino acid sequences. Proteins were further purified by FPLC size-exclusion chromatography using a Superdex 75 10/300 GL (GE Healthcare) column.

#### Circular dichroism (CD)

Far-ultraviolet CD measurements were carried out with an AVIV spectrometer, model 420. Wavelength scans were measured from 260 to 200 nm at temperatures between 25 and 95 °C. For wavelength scans and temperature melts a protein solution in PBS buffer (pH 7.4) of concentration 0.2-0.4 mg/ml was used in a 1 mm path-length cuvette, or 10 times more dilute for 1cm path-length cells.

Chemical denaturation experiments with guanidine hychloride were done with an automatic titrator using a protein concentration of 0.02-0.04 mg/ml and a 1 cm path-length cuvette with stir bar. PBS buffer (pH 7.4) was used for the cuvette solution and PBS+GdmCl for the titrant solution at the same protein concentration. GdmCl concentration was determined by refractive index. The denaturation process monitored absorption signal at 222 nm in steps of 0.1 or 0.2 M GdmCl with 1 min mixing time for each step and at 25 °C. The denaturation curves were fitted by non-linear regression to a two-state unfolding model to extract six parameters: slope and intercept for pre- and post-transition baselines, *m* value and the folding free energy (ΔG_H2O_) (6, 7).

#### Size exclusion chromatography combined with multiple angle light scattering (SEC-MALS)

To evaluate protein quaternary structure, SEC-MALS experiments were performed using a Superdex 75 10/300 GL (GE Healthcare) column, except for samples Rd2NTF2_10, Rd2NTF2_12 and Rd2NTF2_11, for which an Superdex 200 10/300 GL (GE Healthcare) column was used. Then combined with a miniDAWN TREOS multi-angle static light scattering detector and an Optilab T-rEX refractometer (Wyatt Technology). One hundred microliter protein samples of 1-3 mg/ml were injected to the column equilibrated with PBS (pH 7.4) or TBS (pH 8.0) buffer at a flow rate of 0.5 ml/min. The collected data was analyzed with ASTRA software (Wyatt Technology) to estimate the molecular weight of the eluted species.

### Crystallization conditions for solved crystal structures

*Rd1NTF2_05 (PDB ID 6W3D):* (His-tag not cleaved): Protein solution concentration: 56mg/L 1:1 dilution in 0.09M Sodium fluoride; 0.09M Sodium bromide; 0.09M Sodium iodide, 0.1M Tris/BICINE pH 8.5, 50% v/v of 40% v/v PEG 500 MME; 20 % w/v PEG 20000. (Morpheus-HT96 B9 (Gorrec, 2009))

*Rd1NTF2_05_I64F_A80G_T94P_D101K_L106W (PDB ID 6W3F):* (His-tag cleaved): Protein solution concentration: 7.7mg/mL

1:1 dilution in 0.09M Sodium nitrate, 0.09 Sodium phosphate dibasic, 0.09M Ammonium sulfate, pH 6.5 0.1M Imidazole/MES monohydrate (acid), %50v/v of 40% v/v PEG 500 MME; 20 % w/v PEG 20000 (Morpheus-HT96 C1 (8))

*Rd1NTF2_04 (PDB ID 6W3G):* (His-tag cleaved): Protein solution concentration: 48.7mg/mL 1:1 dilution in 0.12M 1,6-Hexanediol; 0.12M 1-Butanol; 0.12M 1,2-Propanediol; 0.12M 2-Propanol; 0.12M 1,4-Butanediol; 0.12M 1,3-Propanediol, 0.1M Imidazole/MES monohydrate (acid), pH 6.5, and 50% v/v of 40% v/v PEG 500 MME; 20 % w/v PEG 20000 (Morpheus-HT96 D1 (8)).

*Rd2NTF2_16 (PDB ID 6W40):* (His-tag cleaved): Protein solution concentration: 24.6mg/mL

1:1 dilution in 0.1 M Sodium Acetate pH 4.5 and 1.0 M di-Ammonium phosphate. Crystals flush freeze with 20% glycerol as cryo protector (JCSG4 H5 -(9)).

*Rd2NTF2_20 (PDB ID 6W3W):* (His-tag cleaved): Protein solution concentration: 22.5mg/mL

1:1 dilution in 0.09 M Sodium nitrate, 0.09 Sodium phosphate dibasic, 0.09 M Ammonium sulfate; 0.1 M Tris (base) & BICINE pH 8.5; 12.5 % v/v MPD; 12.5% PEG 1000; 12.5% w/v PEG 3350 (Morpheus-HT96 C12 (8)).

### Selection of designs for second high-throughput experiment

Using the logistic regression model trained on data derived from the first high-throughput experiment, we predicted the probability of being stable of 11548 models designed by the new generative algorithm, and selected a subset of 10073 with chance > 0.75. A mistake in the calculation of these values resulted in the selection of designs not being strictly above 0.75, but biased towards higher values. In parallel, we clustered all designs by their fold features, and for each cluster we searched for a representative in the 10073 “stable” designs subset. All code and values can be found in the GitHub repository https://github.com/basantab/NTF2analysis, in the Exp2_selection folder.

### Non-redundant set of native NTF2-like domain structures

A non-redundant (<95% sequence identity) set of domain crystal structures was downloaded from the SCOPe database from http://scop.berkeley.edu/astral/pdbstyle/ver=2.05 on September 2015. From this set, NTF2-like domains (d.17.4 SCOPe v2.05 superfamily) were extracted by selecting only *.ent files where the domain record line matched d.17.4 at least partially. For docking, the energy of these structures was minimized using the Rosetta FastRelax protocol with constraints to avoid large structural changes.

### Ligand model preparation in silico docking test

In order to perform ligand docking and binding site design in Rosetta, ligand atomic models need to be processed. For this, we generated ligand conformers using RDKirt (“RDKit: Open-source cheminformatics; http://www.rdkit.org”) and *.mol2 obtained by converting SDF files from the PDB Ligand Expo database using Open Babel using pH 7 for protonation states. Partial charged were assigned using ANTECHAMBER ((10), $AMBERHOME/bin/antechamber -i [input.mol2] -fi mol2 - o [output.mol2] -fo mol2 -c bcc -nc [netcharge]), and net charge calculated from the *.mol2 file.

Rosetta *.param files were obtained using the latest *.mol2 files, by running the scripts provided by RosettaCommons to that end (11). Finally, RDKit conformers were minimized in Rosetta.

### Misfolded-state model for predicting baseline protease stability of de novo NTF2-like protein sequences

As in previous work (1), we calculated the stability score as the difference between observed protease stability and predicted protease stability the design would obtain if misfolded.

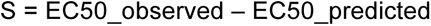

In previous work, the misfolded-state model was trained on stability measurements of sequences expected to be misfolded due to scrambling of the amino acid sequence or introduction of buried charges. The structure of the misfolded-state model enabled it to learn the protease specificities by fitting a 9-residue-length position-specific scoring matrix (PSSM) for each protein, which when applied across the length of the protein, would sum up the cut-rate, assuming the protein was entirely unfolded. With that model and for small proteins, protease stability of misfolded sequences could be predicted with high accuracy (R^2^=0.60 for trypsin, R^2^=0.48 for chymotrypsin). However, for NTF2-like proteins, the same misfolded-state model did not capture protease stability of scrambled sequences (R^2^=0.0), and underestimated the actual protease stability, in some cases, by approximately 10 fold, suggesting that cut-sites are being obscured by partial folding. Expecting that a globular collapse of misfolded sequences would be largely driven by the amino acid composition and length, we fitted a new misfolded-state model by minimizing the squared logarithmic error between observed and predicted EC_50_ over the set of 2694 scrambled sequences. The model predicted EC50 values as a weighted sum of amino acid counts and weights were fitted using linear regression and L1 and L2 regularization (ElasticNet regression) and 5-fold cross validation as implemented in python via scipy (12). The fitted weights largely reflect amino acid hydrophobicity for both trypsin and chymotrypsin (Fig. S26D-E), further supporting that the misfolded-state for NTF2-like proteins is partially collapsed. Across both design rounds, when trained using 5-fold cross validation, the model predicted R^2^=45% (25%) of the trypsin (chymotrypsin) stability variation of pattern-scrambled sequences on an independent holdout dataset (Fig. S26D-E). Including the EC_50_ predictions from the former misfolded-state (9-residues length PSSM) model as a feature in the new model did not improve the performance of the model. Model code, training and testing sets can be found in the GitHub repository https://github.com/basantab/NTF2Gen (BeNTF2seq/molten_globule_model).

### A global PSSM to explain the stability variation of the de novo NTF2 superfamily

To understand position-specific design rules of NTF2-like protein stability, we trained a position-specific scoring matrix (PSSM) model using 6882 designs from both rounds of protease stability screening with EC50 values determined with the updated misfolded-state model described above. The PSSM-model relies on a custom sequence-alignment method associated with the generative-algorithm, which, by stratifying the sequences into 183 independent positions, guarantees structural equivalence between all amino acids in one column of the sequence alignment. Across all positions, weights were then fitted using the same linear regression with L1 and L2 regularization as described above. While the resulting model only explained a fraction of the observed stability variation (R^2^=0.13), upon visual inspection, the PSSM appeared to reflect amino acid preferences correlated with the naturally occurring NTF2-like protein sequence tolerability. Model code, training and testing sets can be found in the GitHub repository https://github.com/basantab/NTF2Gen (BeNTF2seq/fit_PSSM_model).

### Generative algorithm improvement for last round of designs

Stability improvements from the first round to the second round came mainly from biasing the sequence towards higher number of hydrophobic amino acids. A LASSO logistic regression model of stability trained on the second round of designs indicates raw hydrophobicity is not as predictive as in the first round, instead, hydrophobic interactions and sequence-structure agreement (Design score as described in TERMS (4)) are most predictive. To improve these features, we applied the fitted PSSM weights described above as an independent score-term of the energy function. We also incorporated updates in the sequence design Rosetta mover “FastDesign” and Rosetta scoring, that improve designs’ final score and reduce buried unsatisfied polar atoms. This design protocol takes ∼1hr to run in an Intel E5 2680 processor, much faster than previous versions, which took >3hrs due to the extensive optimization trajectory based on point mutations. All files containing the full description of this sequence design protocol and necessary to run the calculations can be found in the GitHub repository https://github.com/basantab/NTF2Gen (BeNTF2seq/design_with_PSSM), along with the final set of 32380 protein models designed with this method and the input backbones.

The increase in parameter combination (backbone) diversity in the last round of designs comes from small changes in the backbone construction code that make fragment assembly more efficient, mainly by tweaking constraint strength and distance values. A history of code changes and their rationale is saved in the GitHub repository https://github.com/basantab/NTF2Gen.

### Design naming

In order to aid clarity, we gave short names to specific designs referenced in this work. Table S17 maps the short names to those used in gene orders. Designs longer than 120 amino acids were not part of the high-throughput experiments; therefore their names (BBLPX) were given at the time of specific gene order. Other short names sometimes used during analysis are written between parentheses.

---

**Fig. S1.**
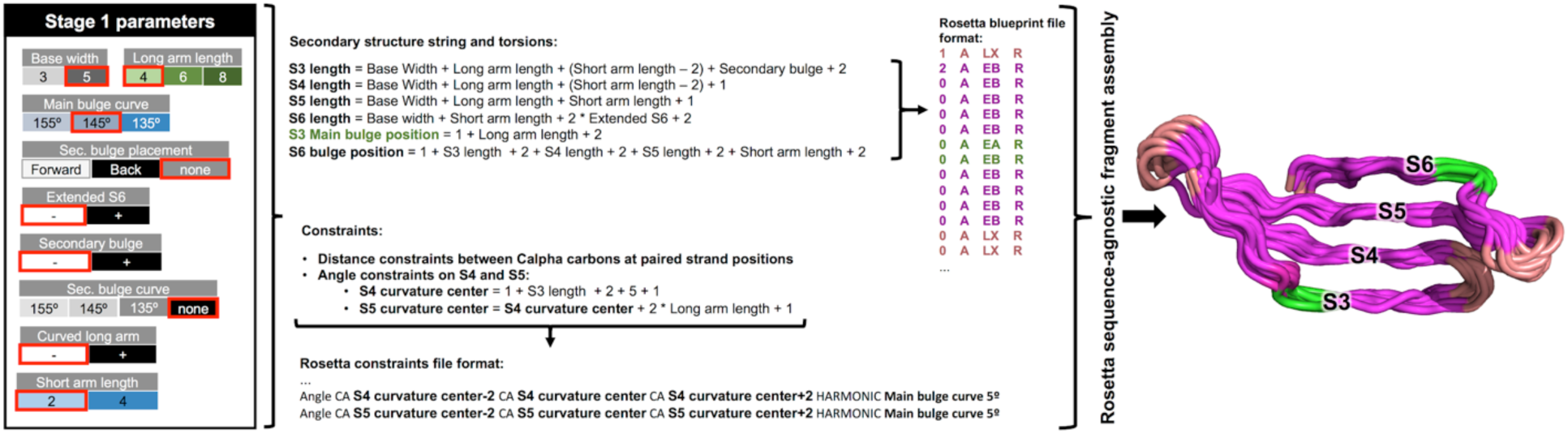
Example of translation of high-level parameters to Rosetta-readable file formats (blueprint and constraint files) for sheet (S3-6) fragment assembly (Stage 1). High-level parameters values selected are in red boxes (left). Examples of some of the conversions from high-level parameters to low-level descriptions are shown in the middle, with the resulting file formats. Fragment assembly produces local structure variations (right).

**Fig. S2.**
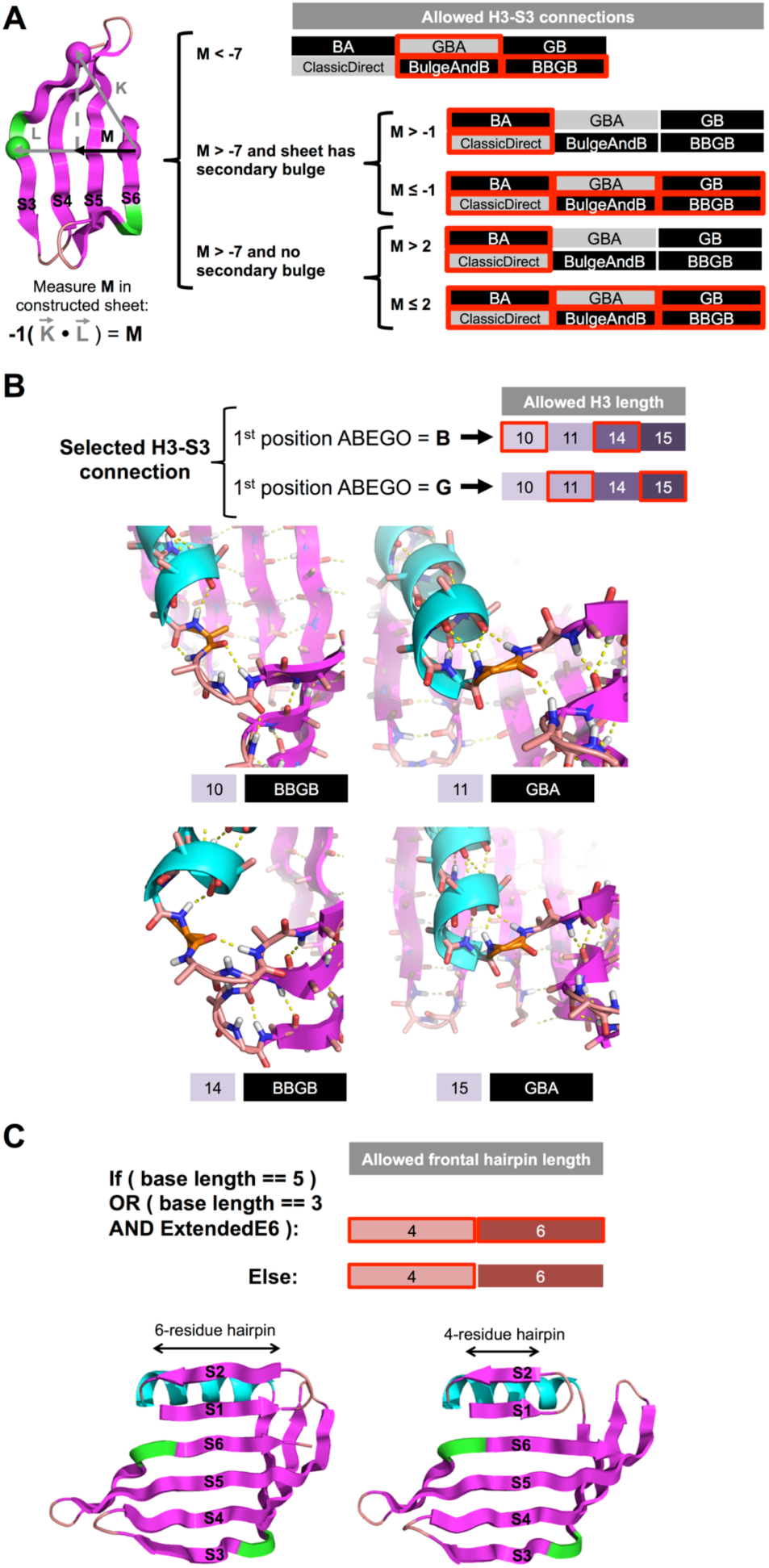
Dependence of stage 2 parameters on the strands built in stage 1. **A:** The protrusion value, calculated from vectors radiating from the S6 end, to the long arm tip and S3 bulge, determines what types of connections can be used between H3 and S3. Low protrusion values (Å) eliminate short connections that would place the C-terminus of H3 too near the center of the sheet, leading to low pocket depth. High protrusion values eliminate long H3-S3 connections that would place the C-terminus of H3 too far from the sheet and limit packing interactions that lead to folding and structured pockets. Intermediate protrusion values allow all types of connections. **B:** The length of H3 is dictated by the H3-S3 connection first positions’ torsion bin. The rigidity of the S2-H3 connection places the C-terminus of H3 in a specific angle to S3, to properly connect H3 to S3, the connection must have a suitable torsion that leads to favorable hydrogen-bond interactions (bottom cartoon examples). **C:** The selection of hairping length (4- or 6-residue S1 and S2) is independent of H3 length and H3-S3 connection, but depends on the length of S6, as it has to fully pair with it.

**Fig. S3.**
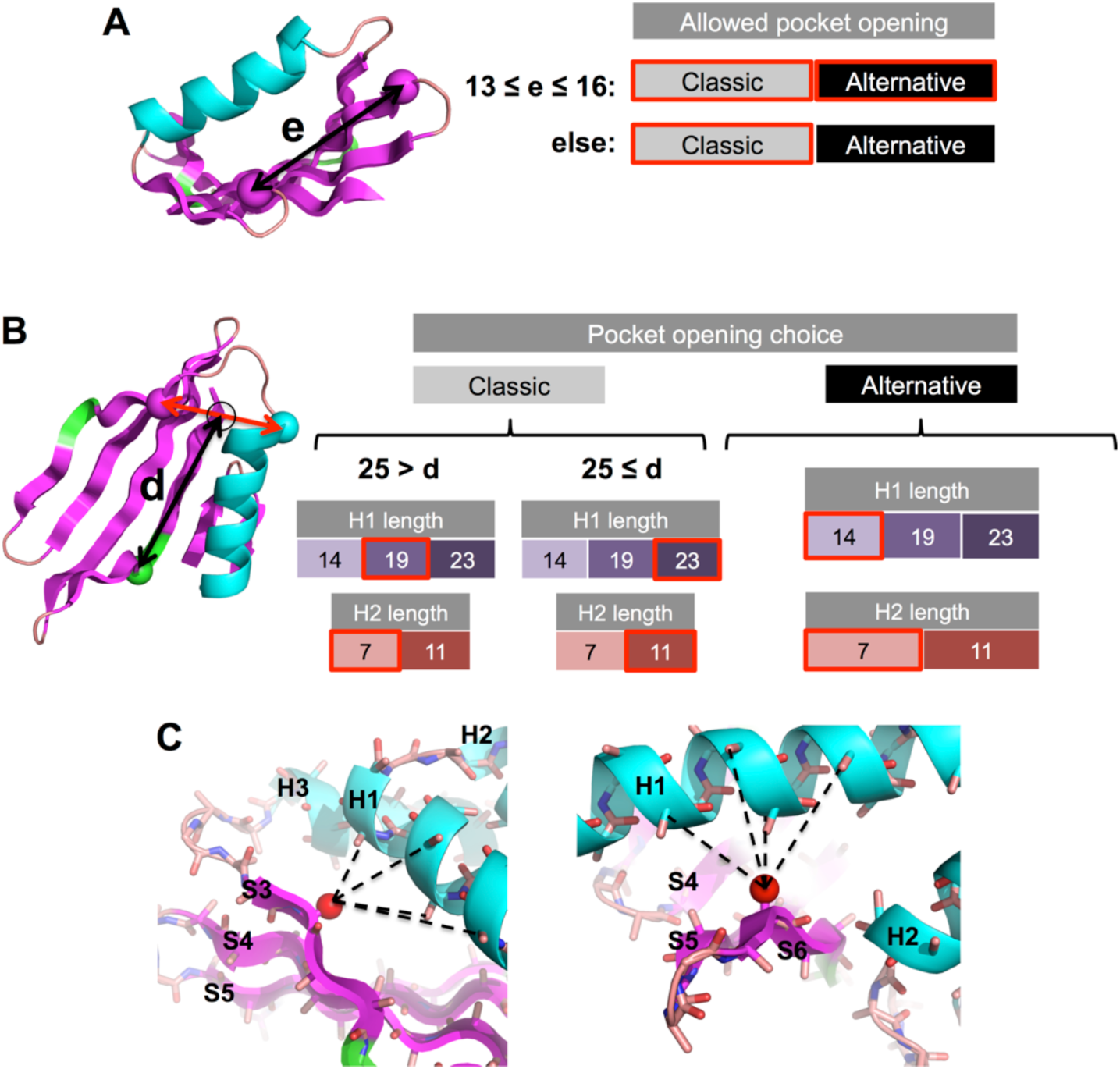
Dependence of stage 3 parameters on the strands built in stage 2. **A:** The distance “e” (Å) between the tip of the long arm and the frontal hairpin dictates wether the pocket opening can be placed between H3 and the H1-H2 loop (Alternative placement), or not. **B:** Once the opening placement is selected, this and the ditance between the S6 bulge and the H3-S3 connection dictate the lengths of helices 1 and 2. For alternative openings, H2 and H1 are as short as possible, to leave a space between the H1-H2 loop and H3. If the opening placement is Classic, then the distance between the S6 bulge and the H3-S3 loop distates the lengths of H1 and H2: if distance “d” is longer than 25Å, then H1 and H2 are both extended by a full turn (4 residues), to span the surface between the N-terminus of S3 and the short arm. **C:** In order to steer the placement of H1 and H2 such that energetically favorable side-chain packing can be achieved, constraints (dashed lines) are used to tie the N and C termini of H1 to the sheet. The N-terminus of H1 is tied to the N-terminus of S6 such that the C_β_ (red spheres) of this position points at the splace left by side-chains on the side of H1 (C, right). The same is done between the C-terminus of H1 and the N-terminus of S3 (C, left).

**Fig. S4.**
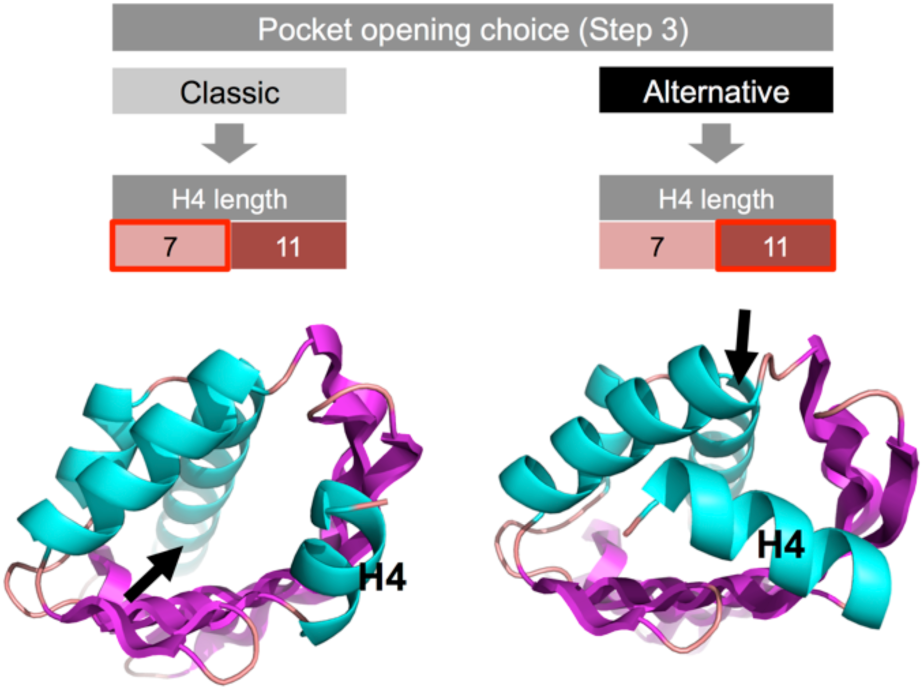
Dependence of stage 4 parameters on the strands built in stage 2. In case a C-terminal helix is constructed, the type of pocket opening (black arrows) determines its length and placement. A shorter H4 interacts with the long arm and extends the pocket outwards in pockets with Classic openning. In pockets with Alternative opening, H4 takes the space between the frontal hairpin and the long arm. In both cases, constrinats are placed between the C-terminus of H4 and the rest of the structure to steer fragment assembly and achive the described H4 placement.

**Fig. S5.**
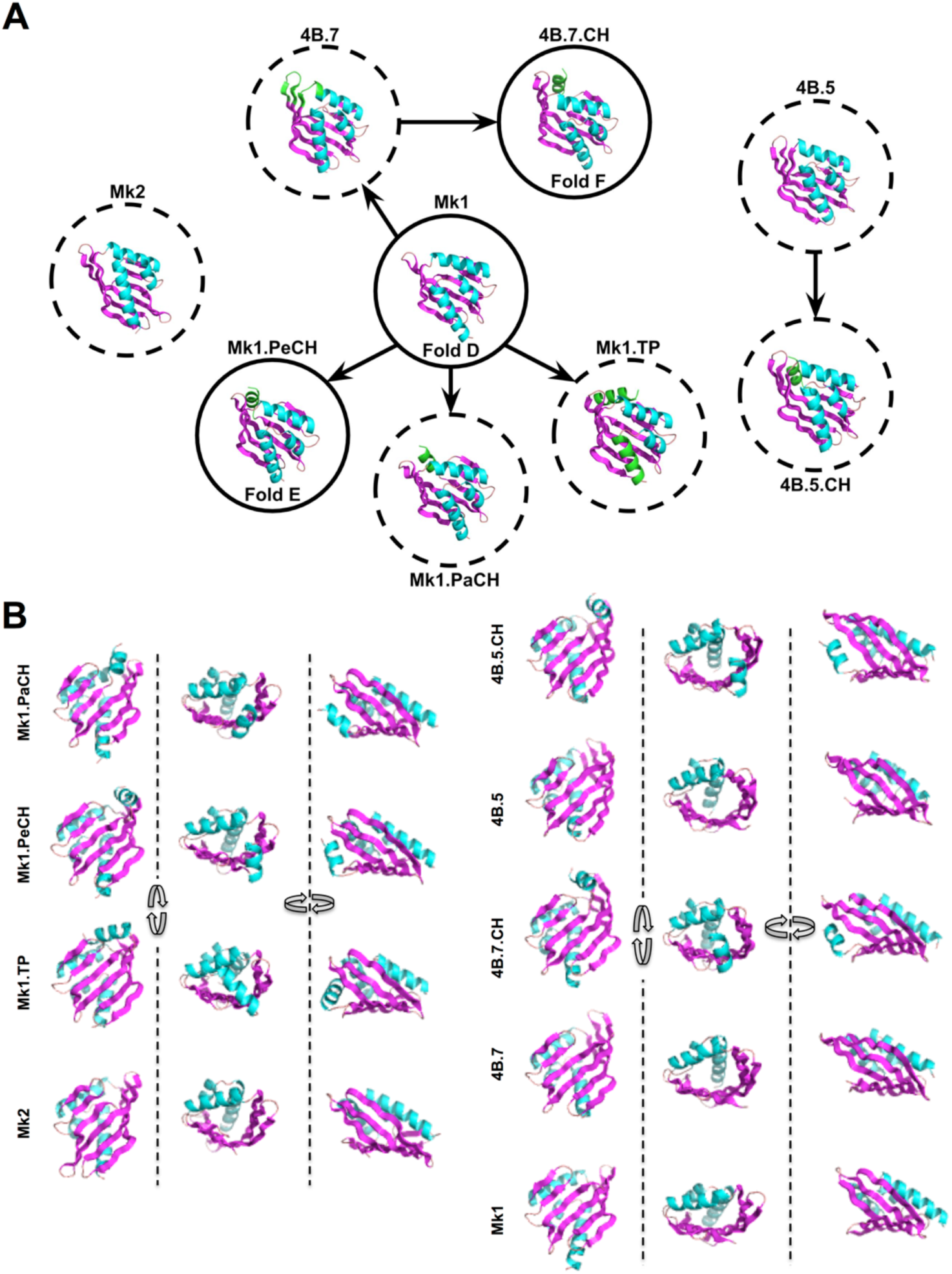
*De novo* NTF2-like designs tested in the first high-throughput experiment. **A:** Diagrams of the different blueprints tested, and how they map to previously published blueprints (solid circles), or not (dotted circles). Arrows indicate single changes that convert one blueprint to the other. **B:** Alternative visualization of models in figure A.

**Fig. S6.**
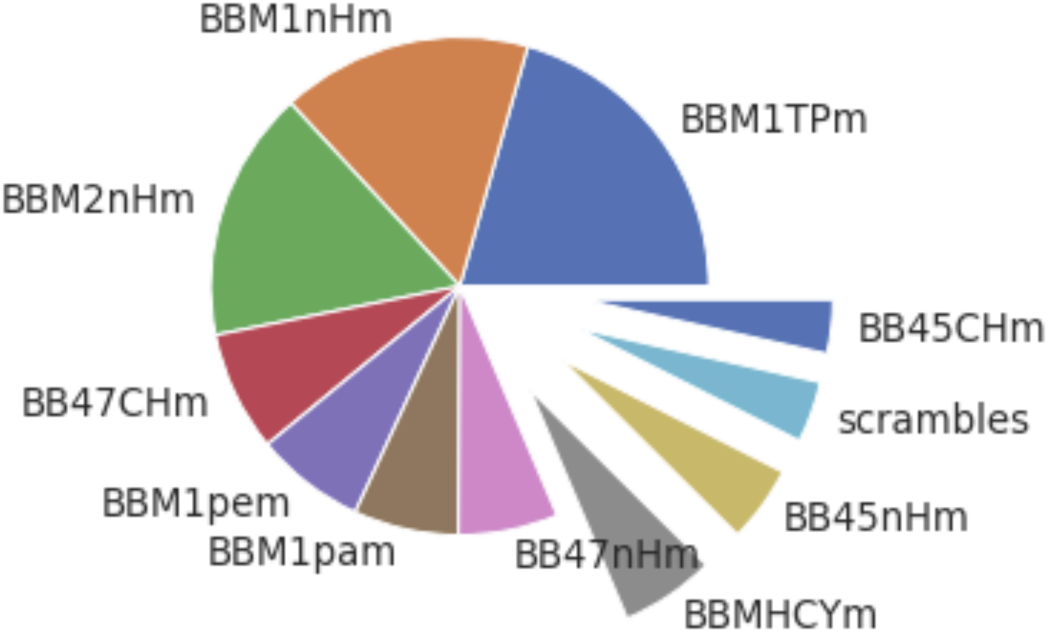
Population of stable designs, separated by family.

**Fig. S7.**
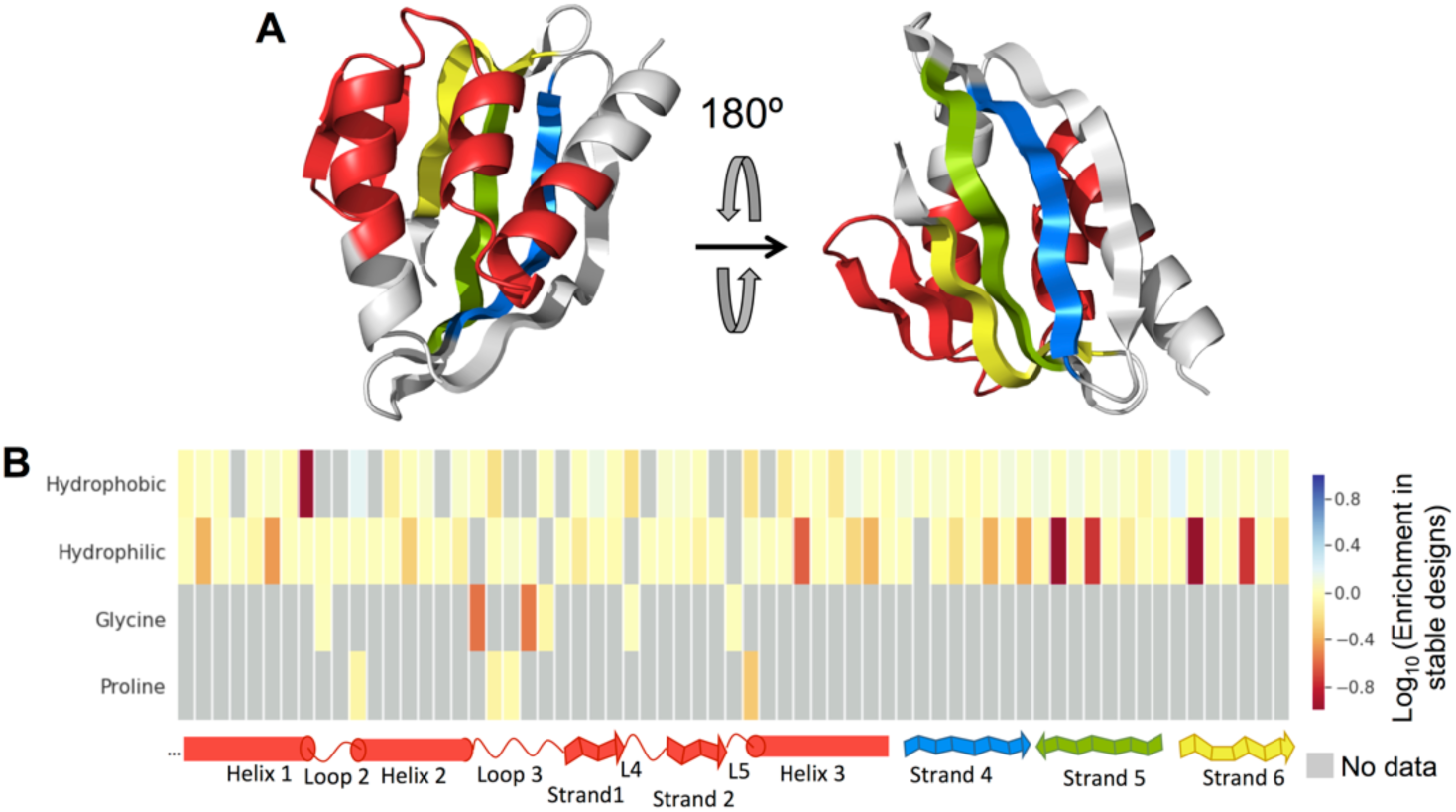
Enrichment pattern for first-round designs. **A:** Structural legend of analyzed postions, presented in the bottom of B as a continuous stretch, each color represents a separate sequence stretch. **B:** Enrichment heatmap. Each column is a structuraly homologous position in all tested designs. Grey cells indicate that either the amino-acid was not sampled at that postion, or the enrichment was not statistically significant.

**Fig. S8.**
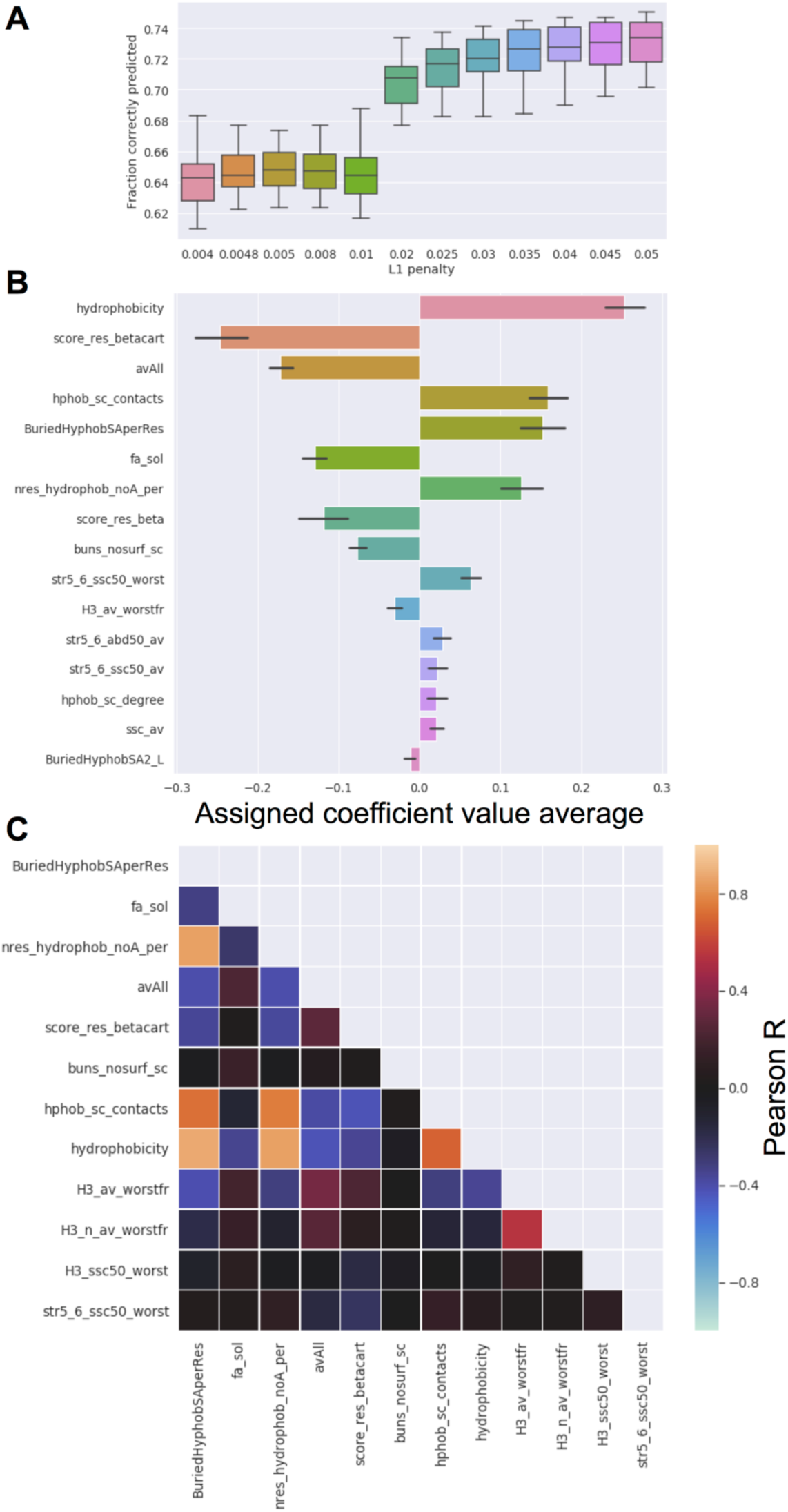
Logistic regression model on first-round designs. **A**. Boxplot of model accuracy on test set for different values of L1 penalty. Each box represents 40 different random partitions of the dataset (with replacement), with one third of it as test set in each case. **B**. Absolute weights of the 16 features with the highest average weights in the 40 dataset partitions. **C**. Correlation matrix for the top 12 features.

**Fig. S9.**
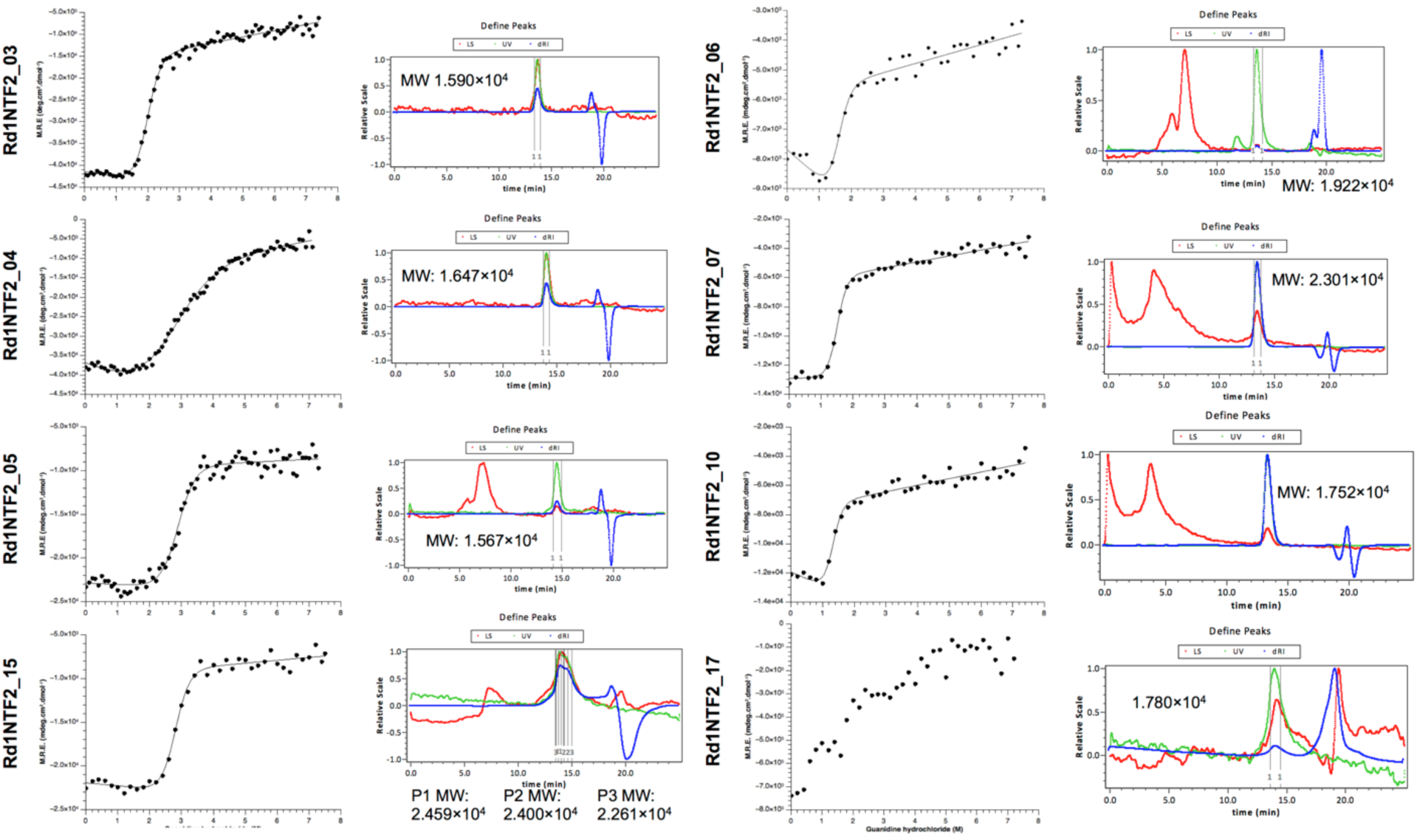
Experimental characterization of first-round designs. Column 1: Guanidinium chloride titrations following circular dichroism at 222nm, circular dichroism spectra, and size-exclussion chromatography followed by multi-angle light scattering.

**Fig. S10:**
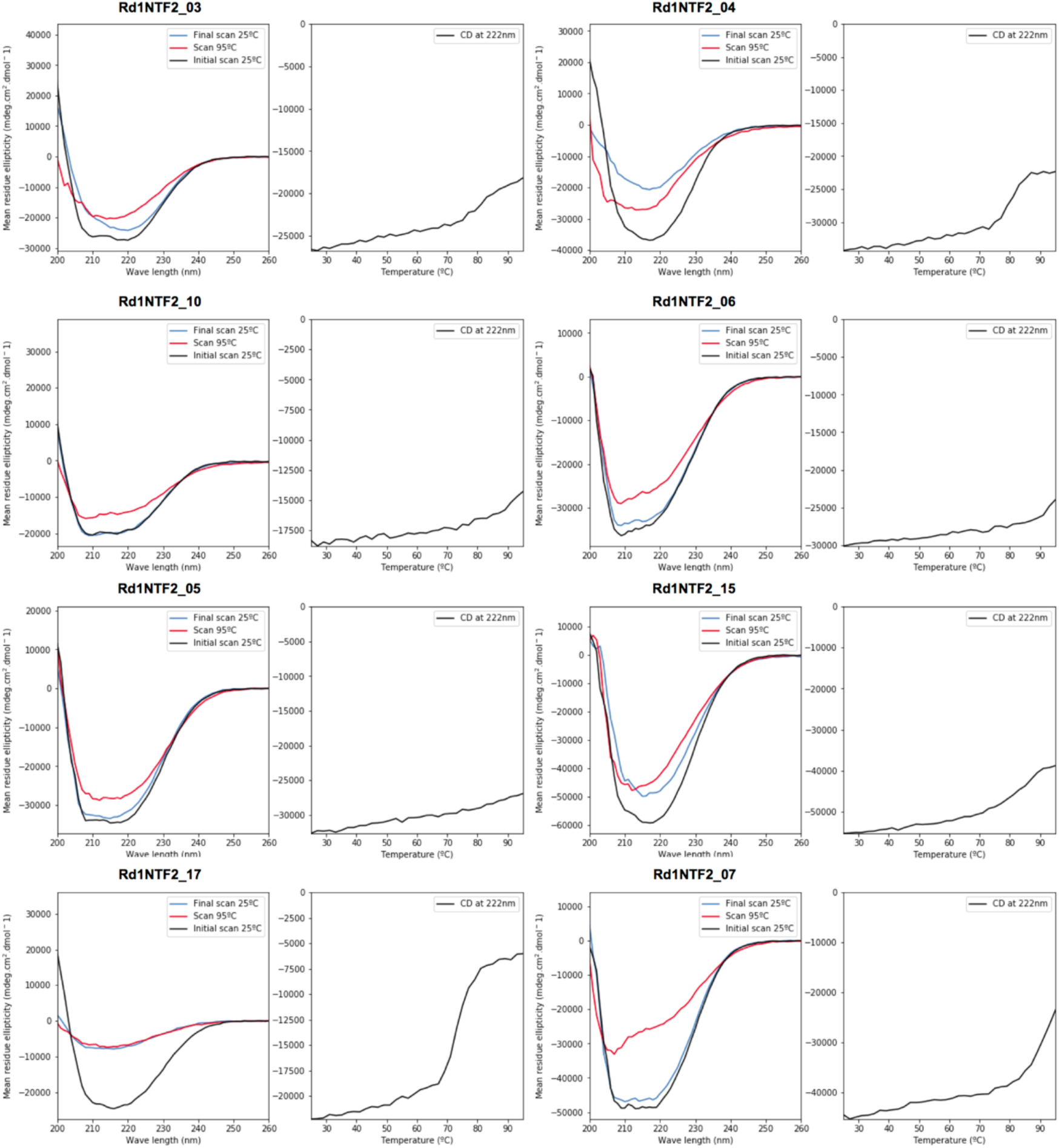
Experimental characterization of first-round designs. Circular dichroism spectra at 25°C and 95°C and temperature curves following circular dichroism at 222nm.

**Fig. S11.**
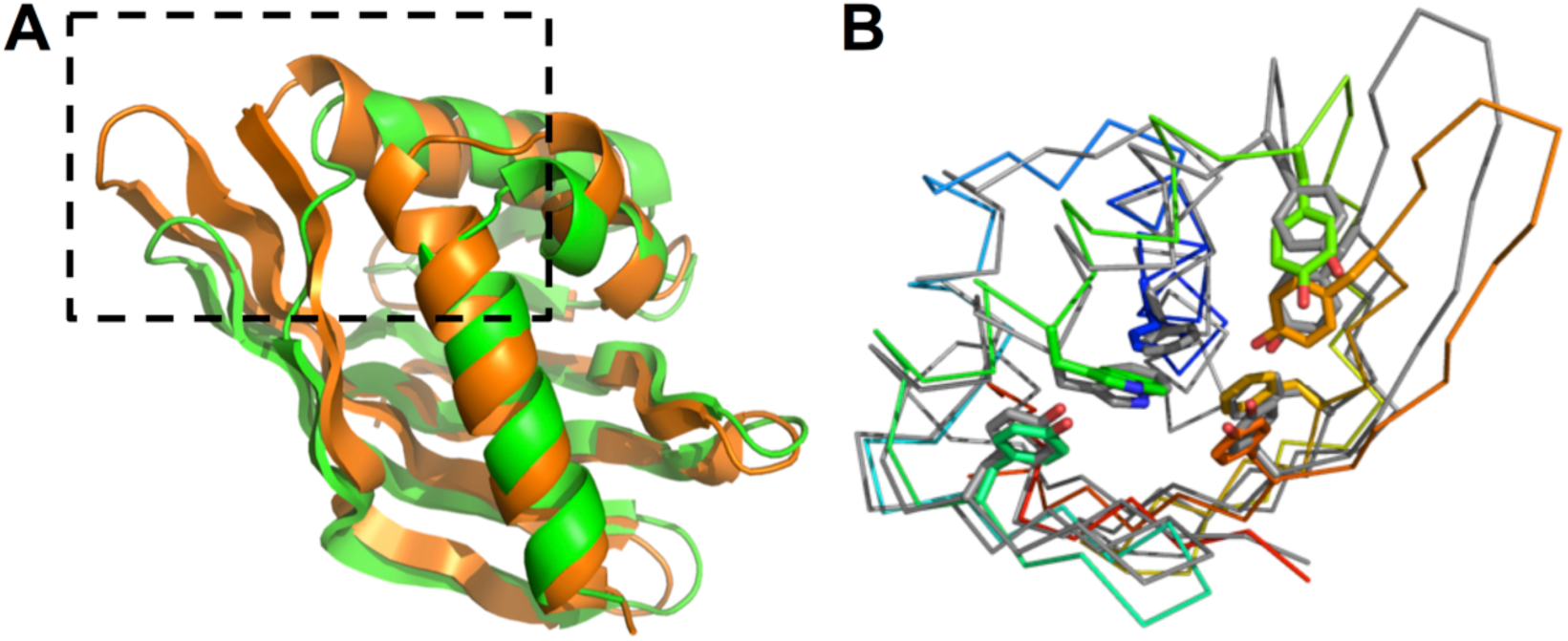
Rd1NTF2_04 crystal structure. **A:** Comparison between the Rd1NTF2_04 structure (orange) and the most similar structure from Marcos et al. (Fold C, green), highlighting with a box the structural elements that extend the concave sheet space outwards in Rd1NTF2_04. **B:** Core side-chain conformations in the Rd1NTF2_04model (gray) and structure (rainbow).

**Fig. S12.**
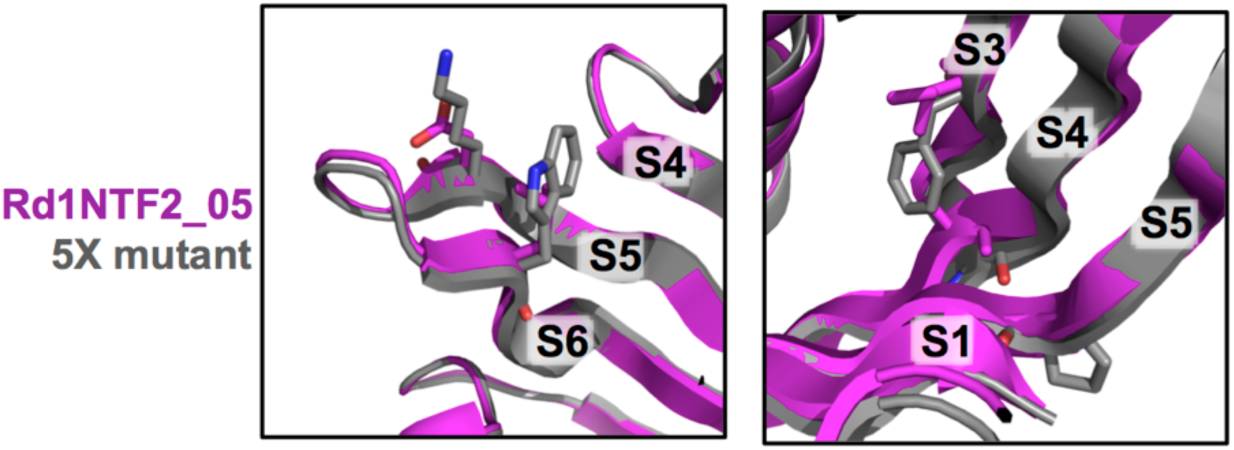
Rd1NTF2_05 mutations to correct structure. Left pannel: Muatations D101K and L106W on strand 5 and 6, respectively. Right pannel: Mutations I64F, A80G and T94P on strands 3, 4 and 5 respectively. Note how after changing A 80 to G the backbone structure relaxes into a deeper arch, and F64 takes the space left by the alanine side chain.

**Fig. S13:**
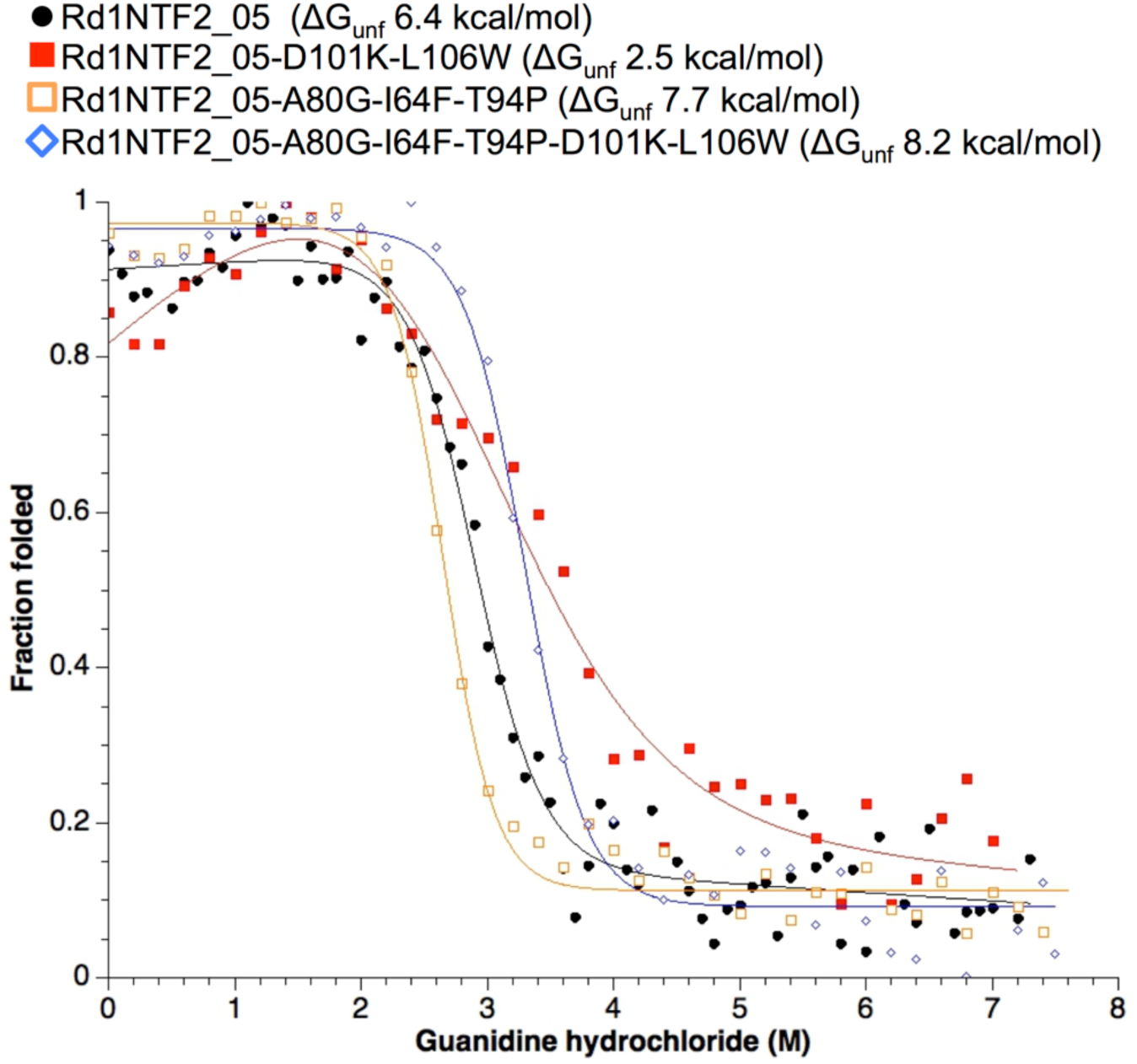
Guanidinium titration of Rd1NTF2_05 mutants.

**Fig. S14.**
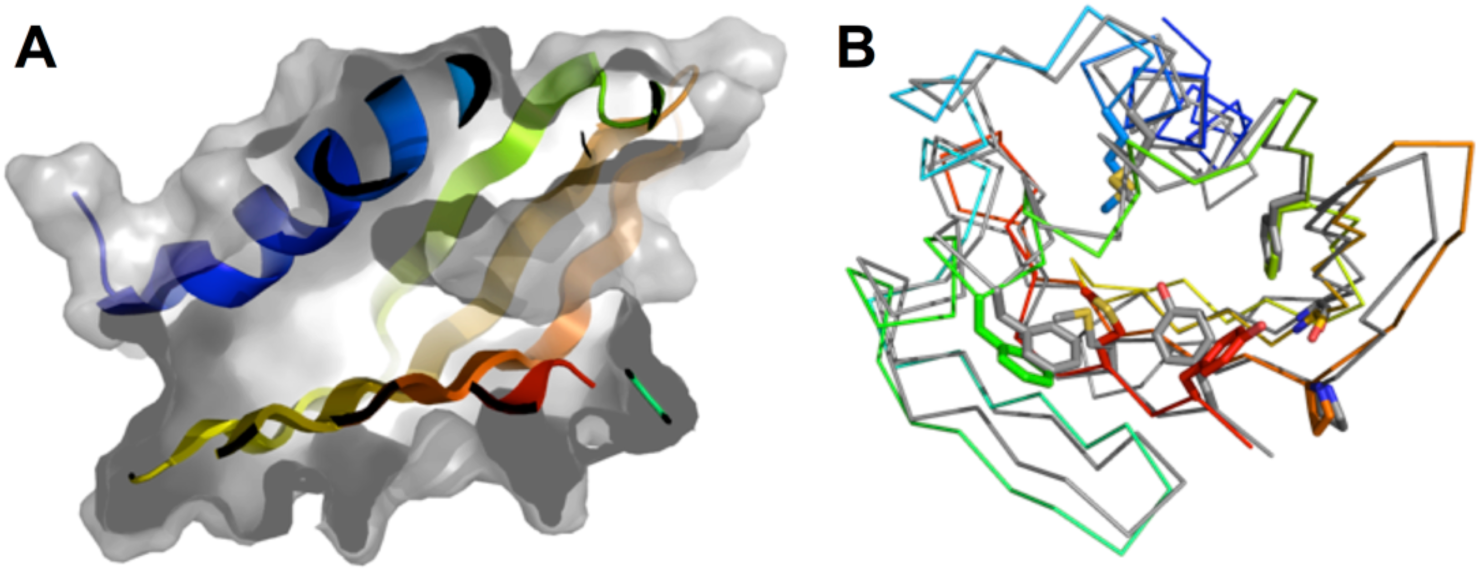
Rd1NTF2_05 5-fold mutant structure. A: Pocket in the Rd1NTF2_05 5-fold mutant structure. B: Comparison of side-chain rotamers between crystal structure (rainbow) and model (gray).

**Fig. S15:**
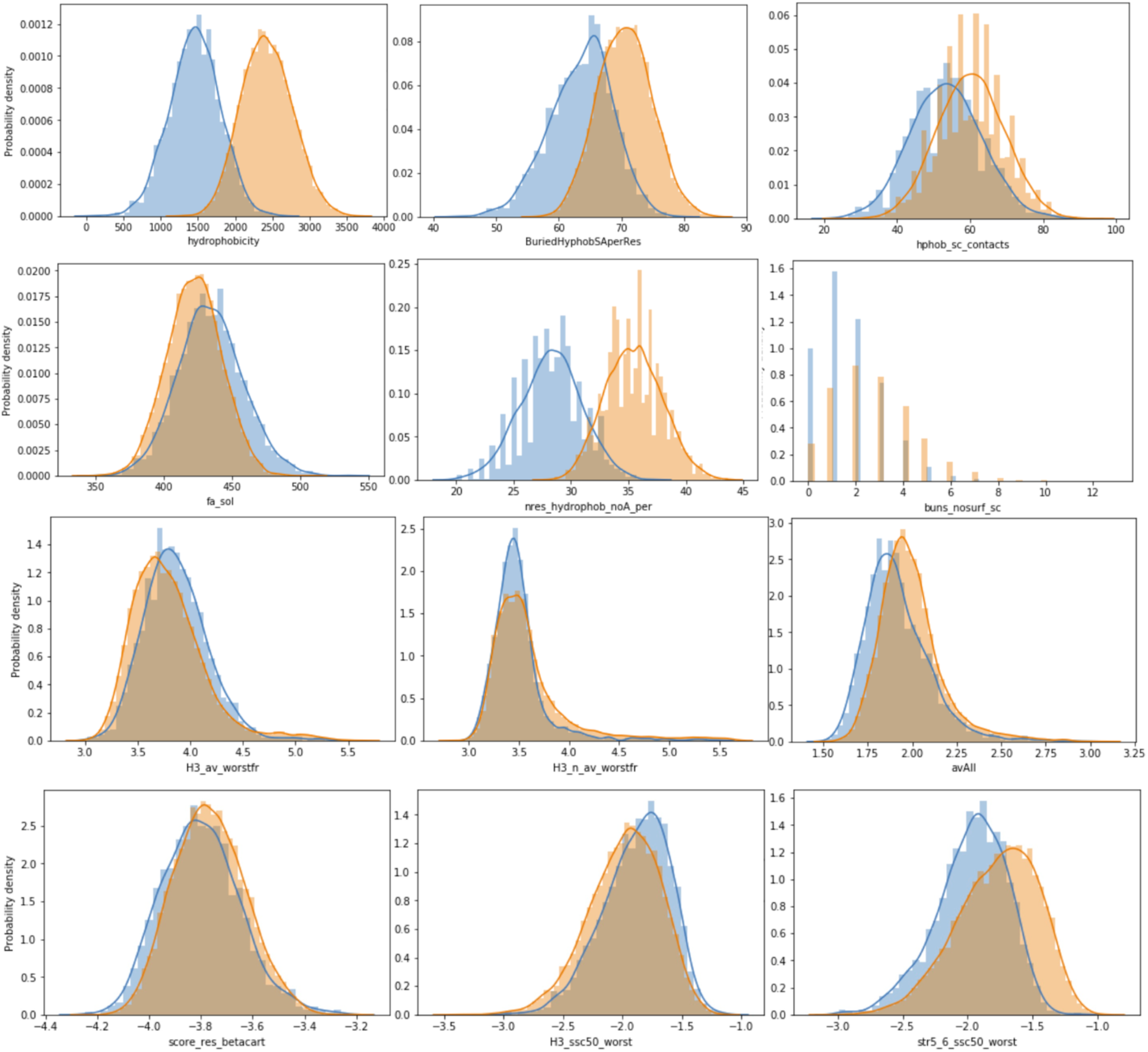
Feature distributions for the first (blue) and second (orange) high-throughput experiments. Here we present only the top features by importance according the the linear model trained using data from the first high-throughput experiment.

**Fig. S16:**
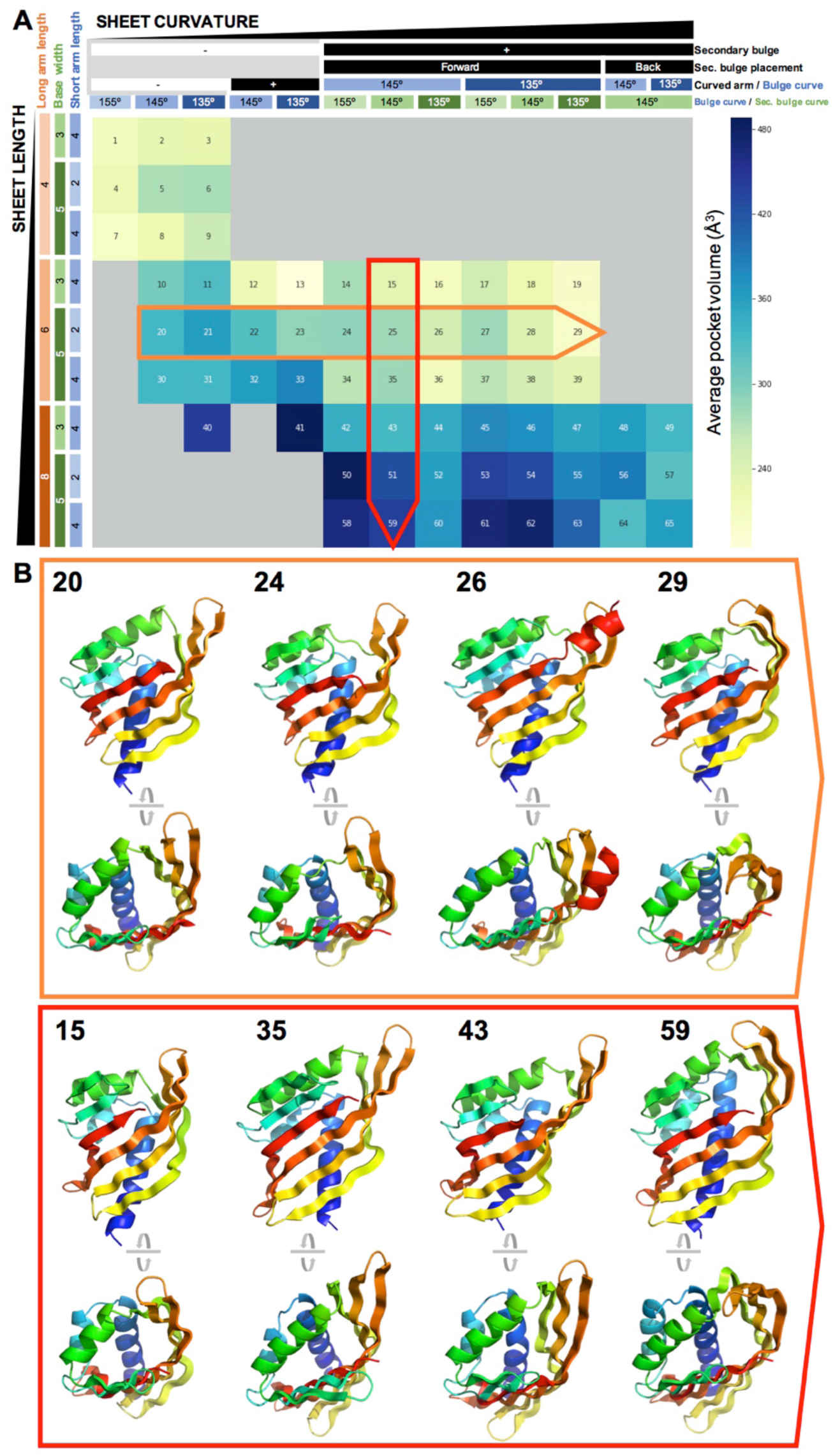
Modes of pocket modulation by sheet structure. **A:** Sheet parameter table, with y axis mapping to sheet length and x axis mapping to curvature. Colored cells are sampled parameter combinations, and the tone indicates average pocket volume after sequence design. Two traversals, one through the y axis (red), and one through the x axis (orange), show how sheet curvature and length modulate pocket volume. **B:** Exemplar structures with pocket volumes close to average, through each of the two above traversals.

**Fig. S17.**
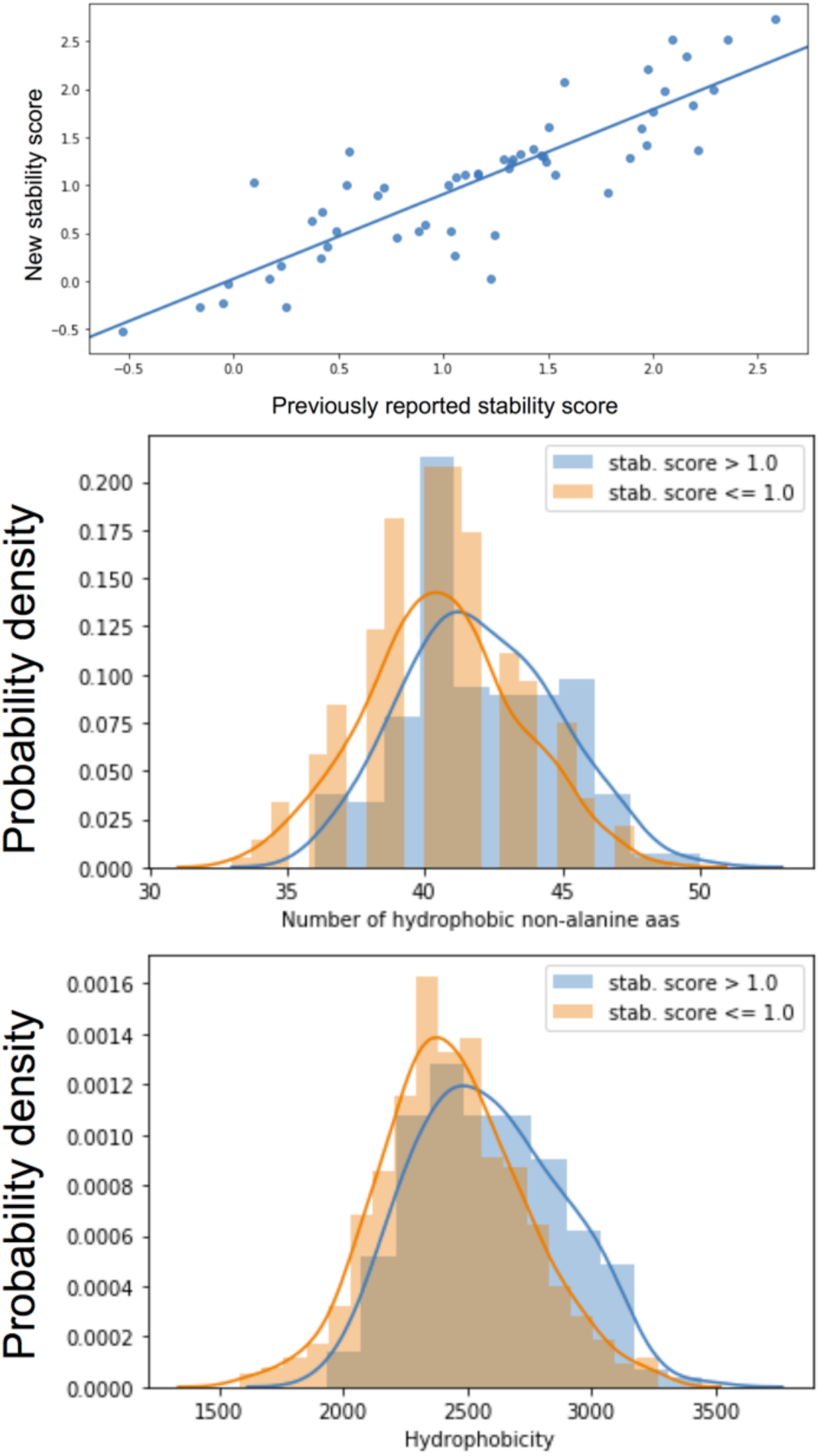
Top: Stability scores of designs tested in both rounds as controls. The linear fit suggests the stability score values are the same between assays. Middle and bottom: Sequence features that best predict stability in second-round scrambled controls: Hydrophobicity and the number of non-alanine hydrophobic residues. These features have the highest weights in the most parsimonious logistic models of stability.

**Fig. S18.**
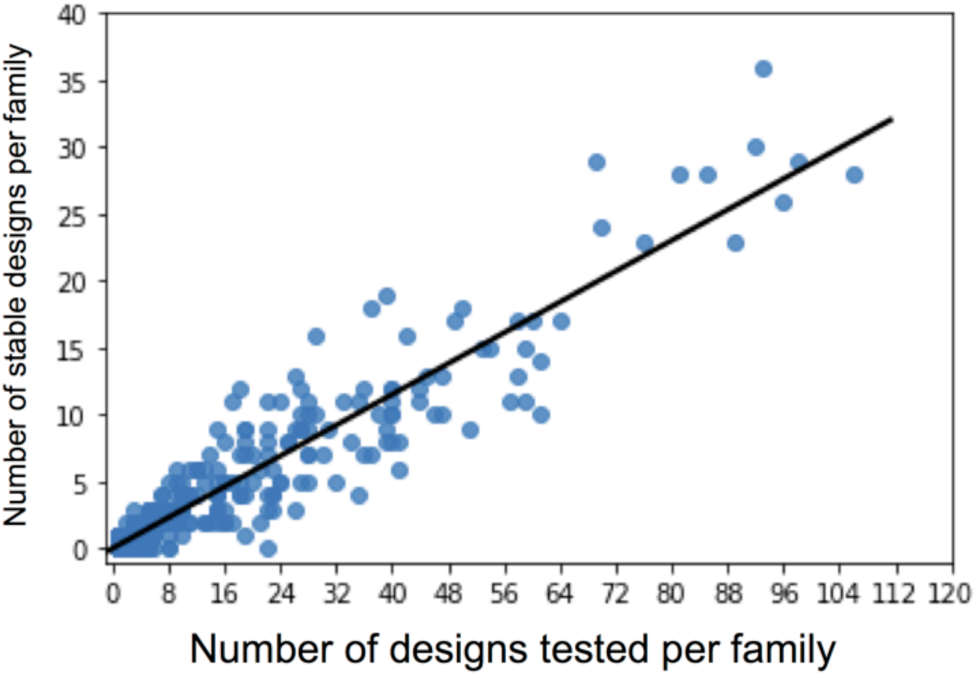
The number stable representatives for each tested family is similar to the population average. The black line is a linear fit, and has a slope of approximately 1/3. Most tested families without stable representatives had less than 10 tested members.

**Fig. S19.**
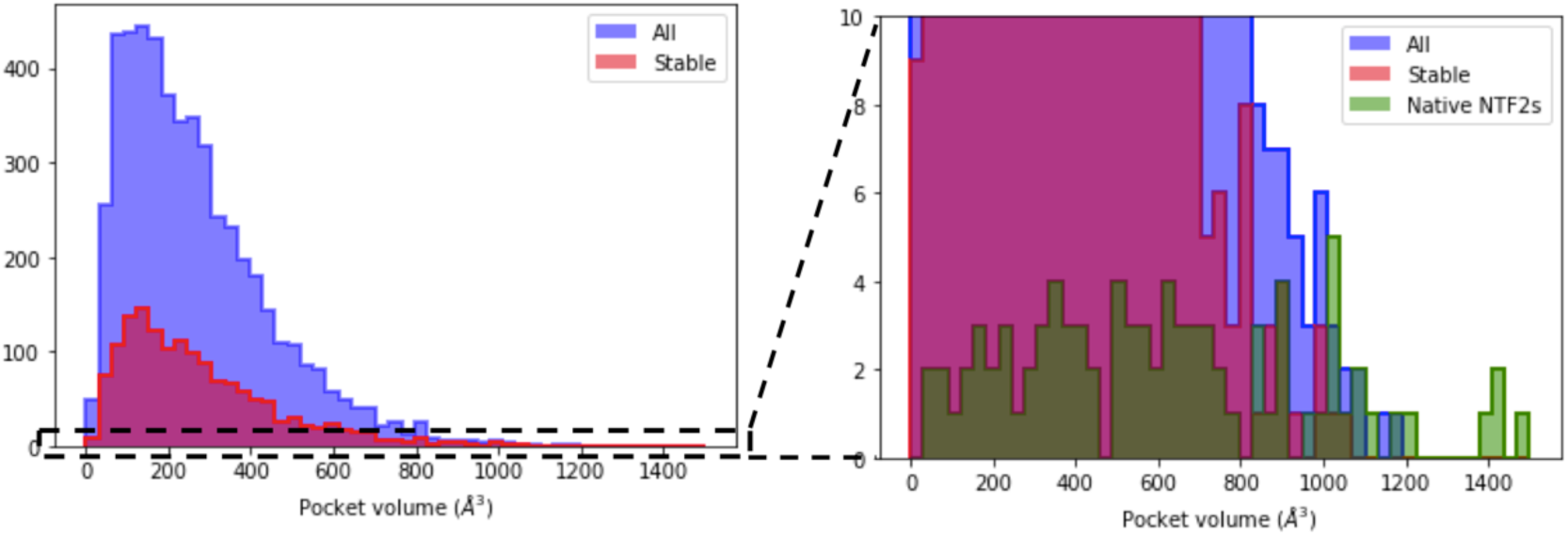
Stable second-round design cover most of the pocket volume range of native NTF2-like proteins, and for most of that range, there are more *de novo* designs.

**Fig. S20.**
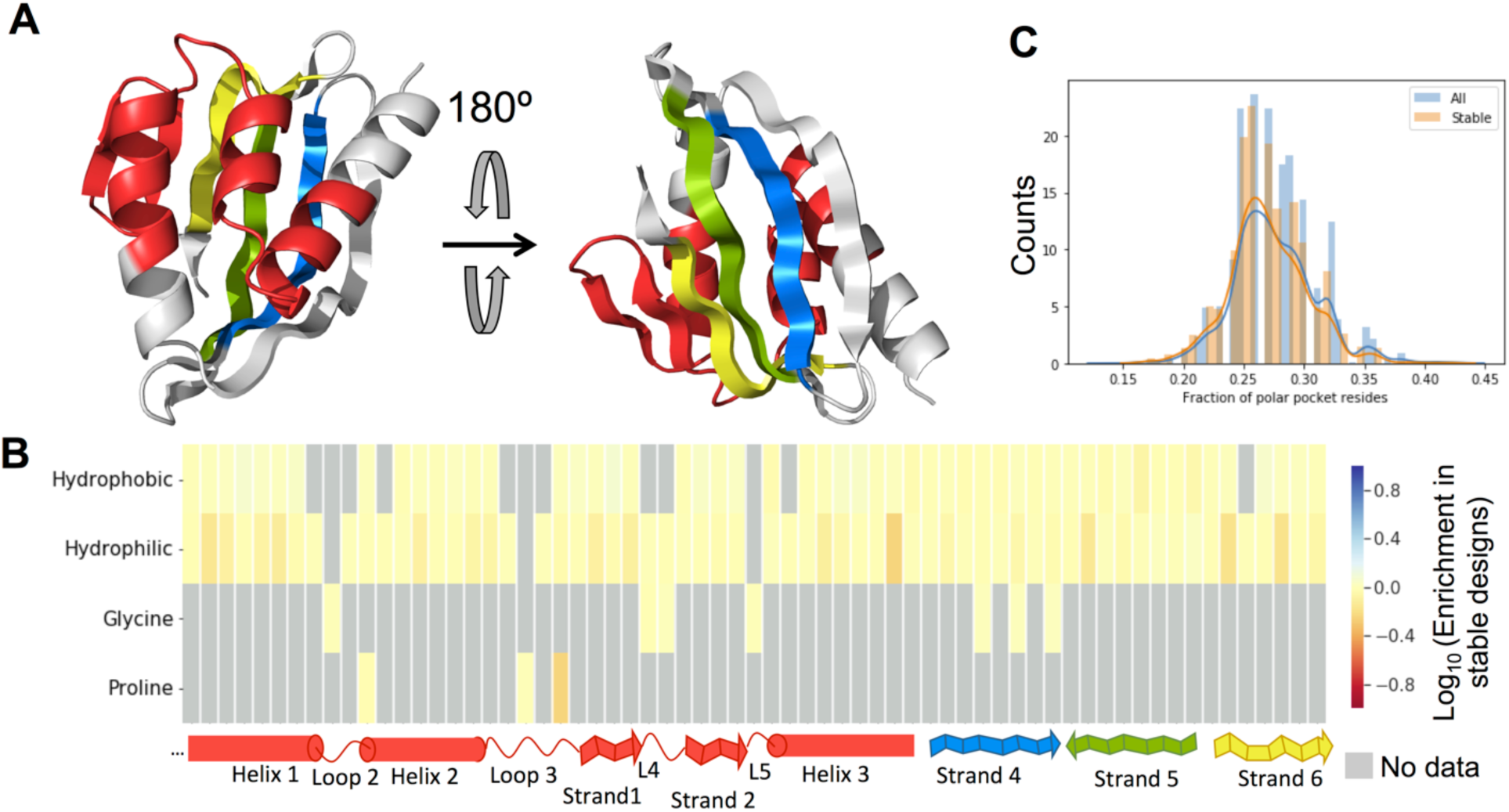
Enrichment pattern for second-round designs. **A:** Structural legend of analyzed positions, presented in the bottom of B as a continuous stretch, each color represents a separate sequence stretch. **B:** Enrichment heat map. Each column is a structurally homologous position in all tested designs. Grey cells indicate that either the amino acid was not sampled at that position, or the enrichment was not statistically significant. C: Fraction of polar pocket residues in all and stable second-round designs. These proteins have on average 24 pocket residues.

**Fig. S21.**
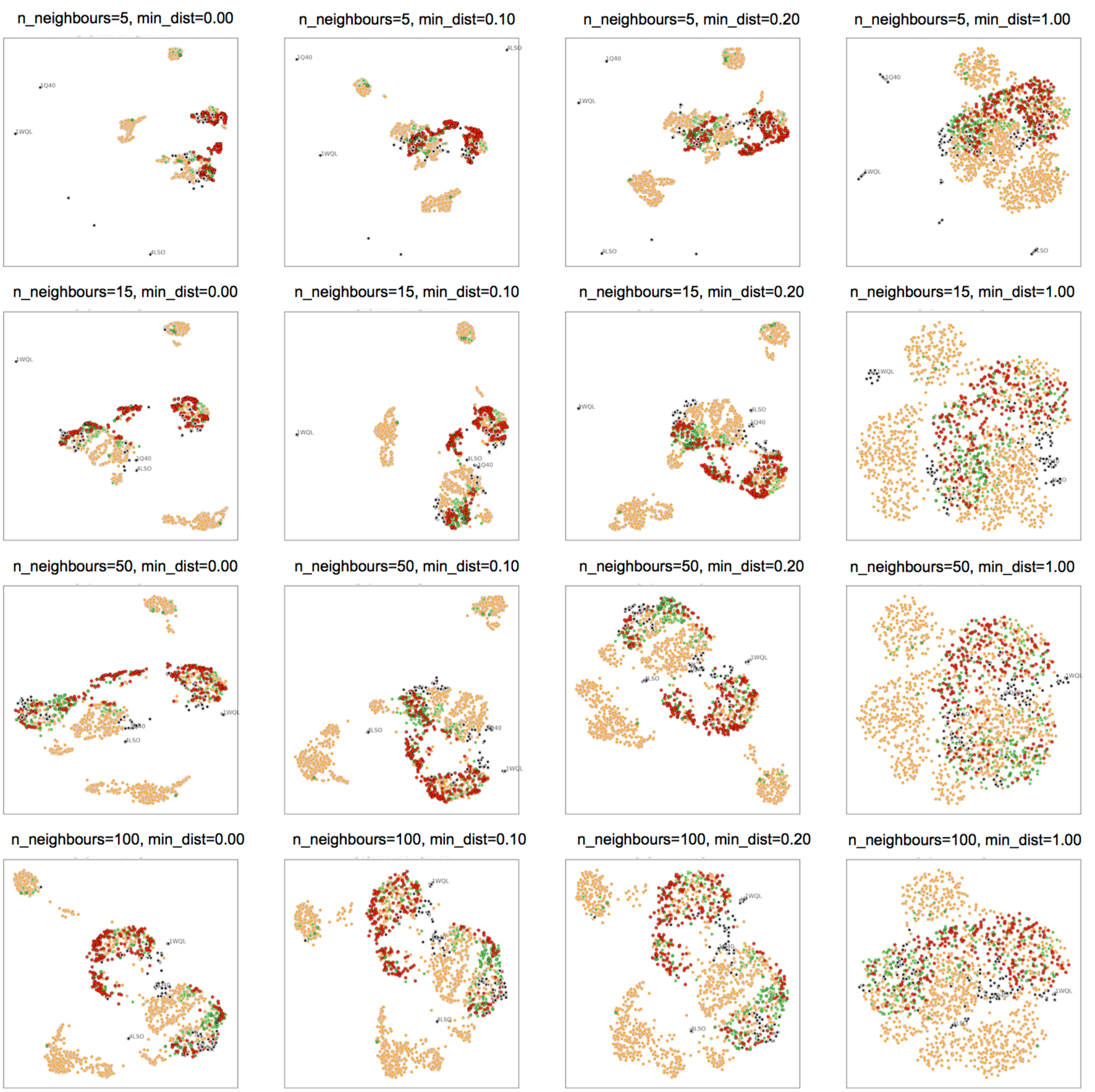
UMAP hyperparameter exploration. Color legends are the same as in Figure 3D.

**Fig. S22.**
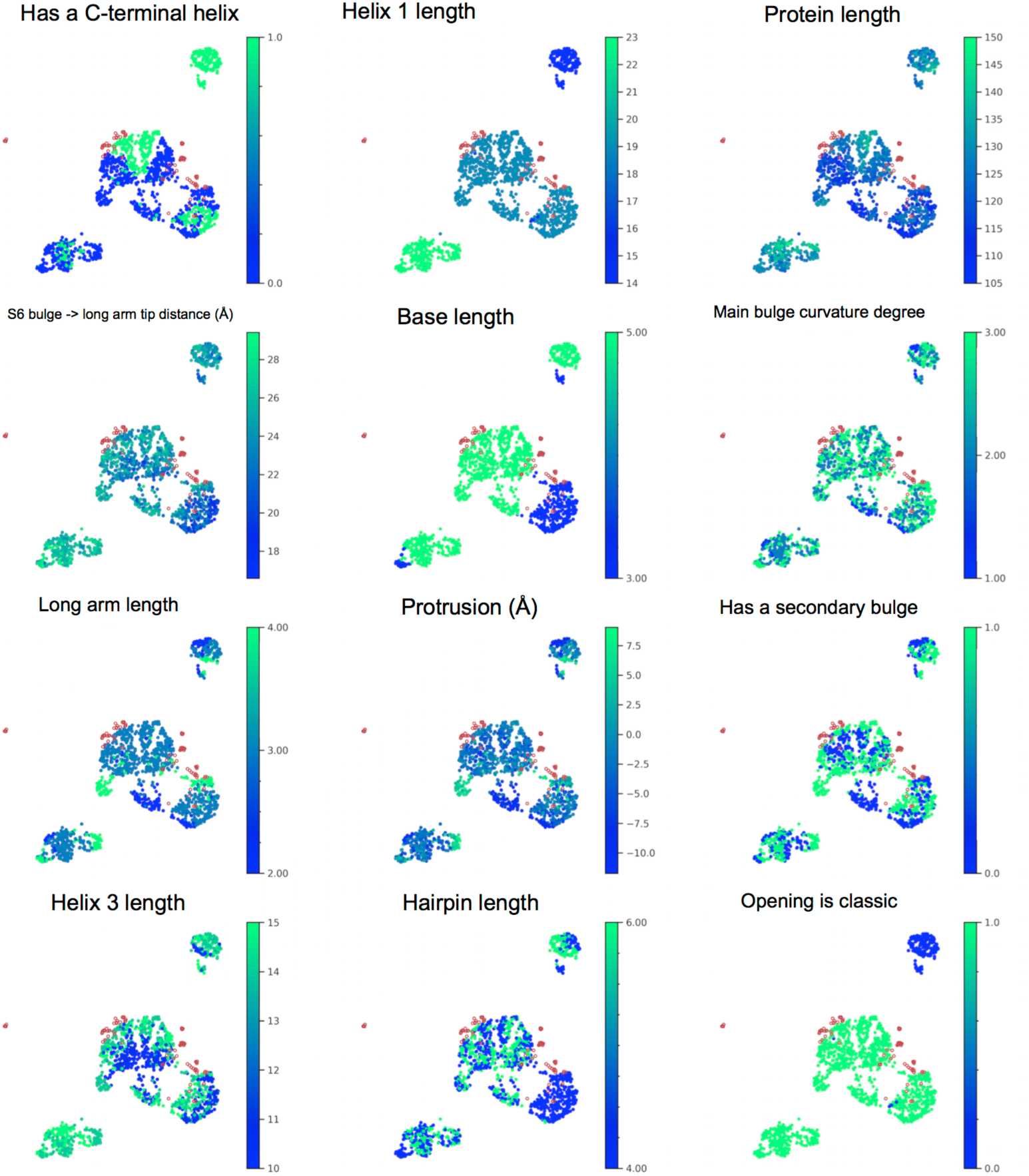
UMAP embedding colored by different generative algorithm parameters used to construct each structure. Red dots are native structures.

**Fig. S23.**
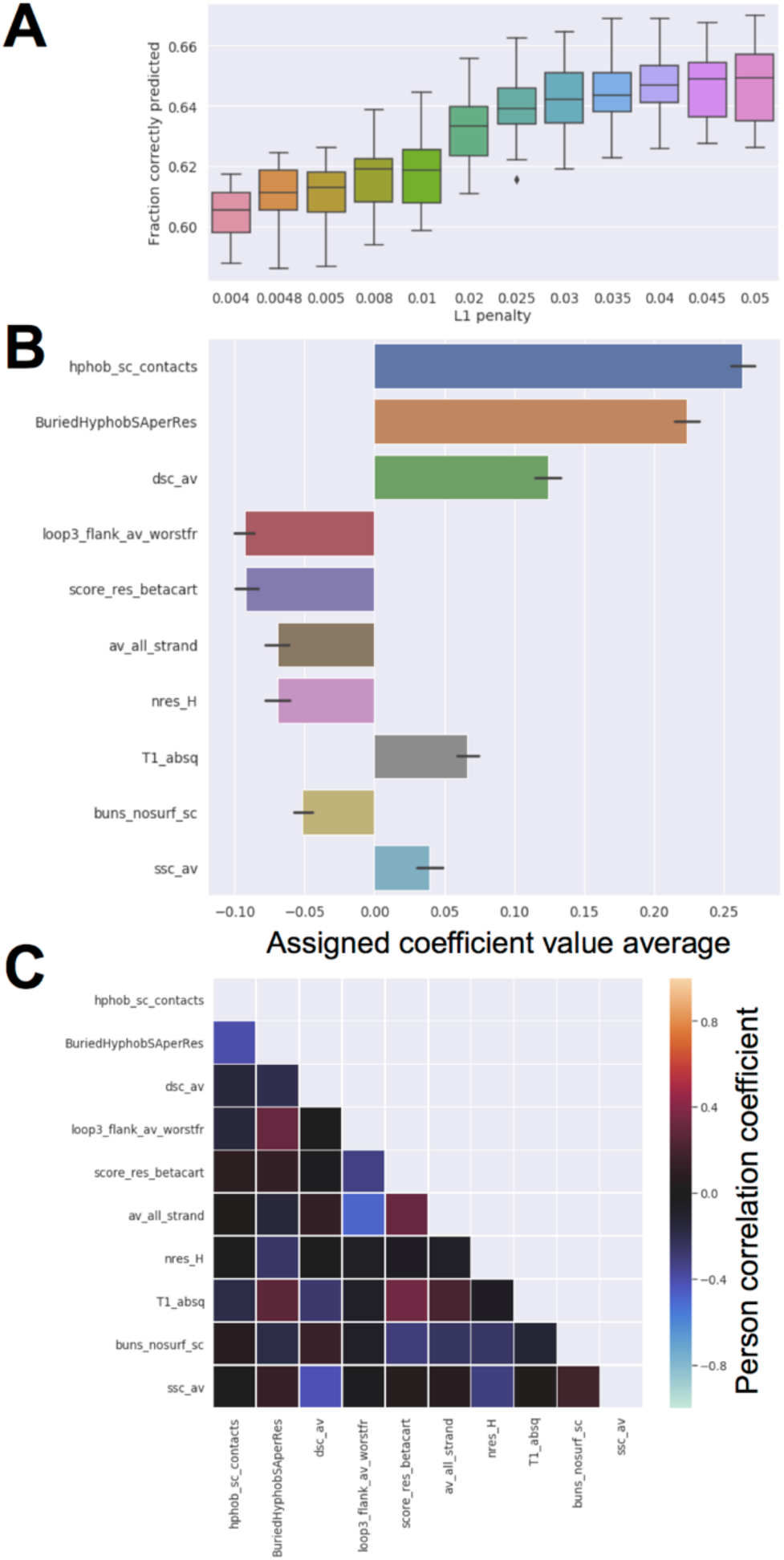
Logistic regression model from second-round designs. **A**. Boxplot of model accuracy on test set for different values of L1 penalty. Each box represents 40 different random partitions of the dataset (with replacement), with one third of it as test set in each case. **B**. Absolute weights of the 10 features with the highest average weights in the 40 dataset partitions. **C**. Correlation matrix for top 10 features.

**Fig. S24.**
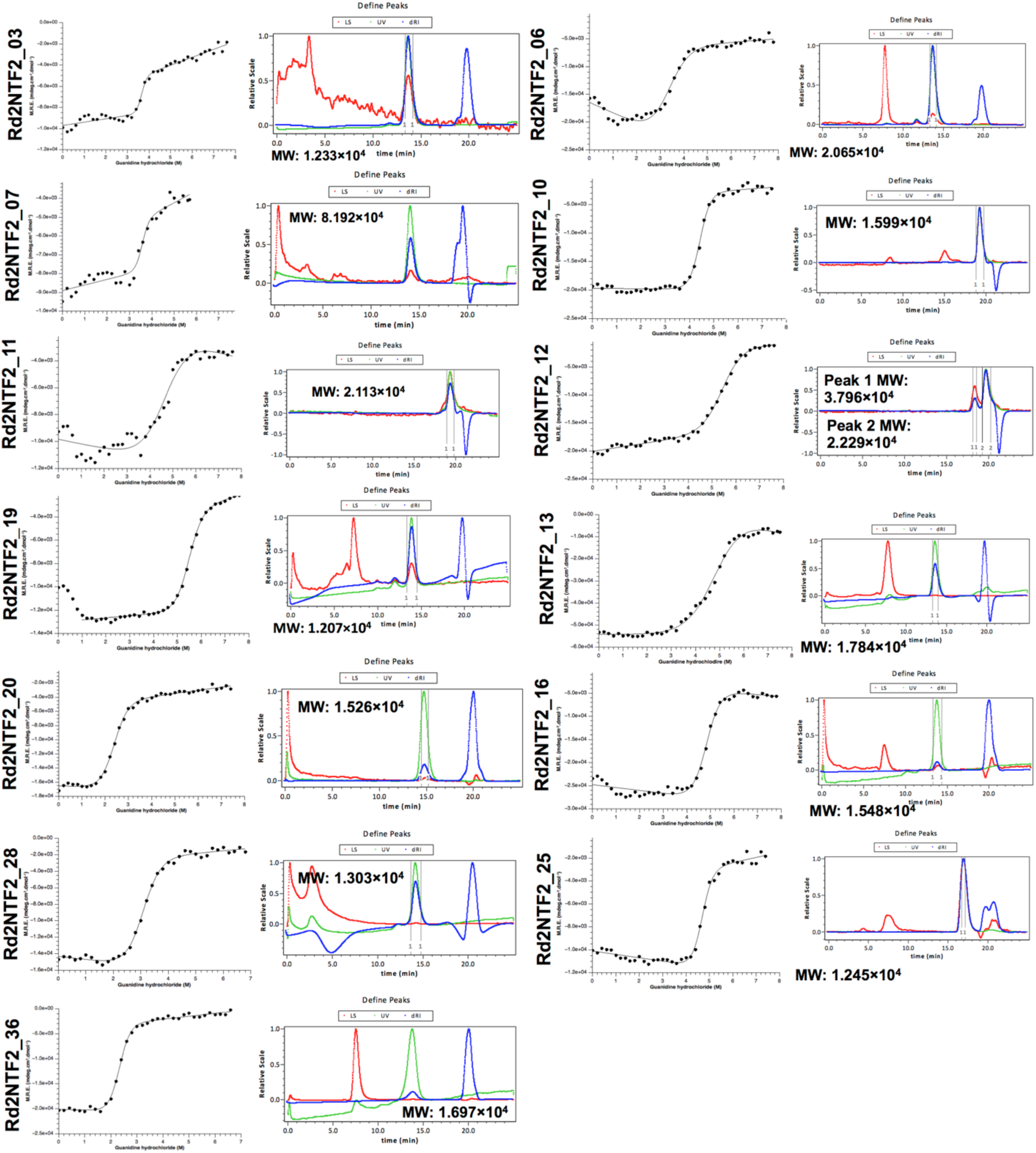
Experimental characterization of second-round designs. Column 1: Guanidine hydrochloride titrations following circular dichroism at 222nm. Column 2: Size-exclusion chromatography followed by multi-angle light scattering.

**Fig. S25.**
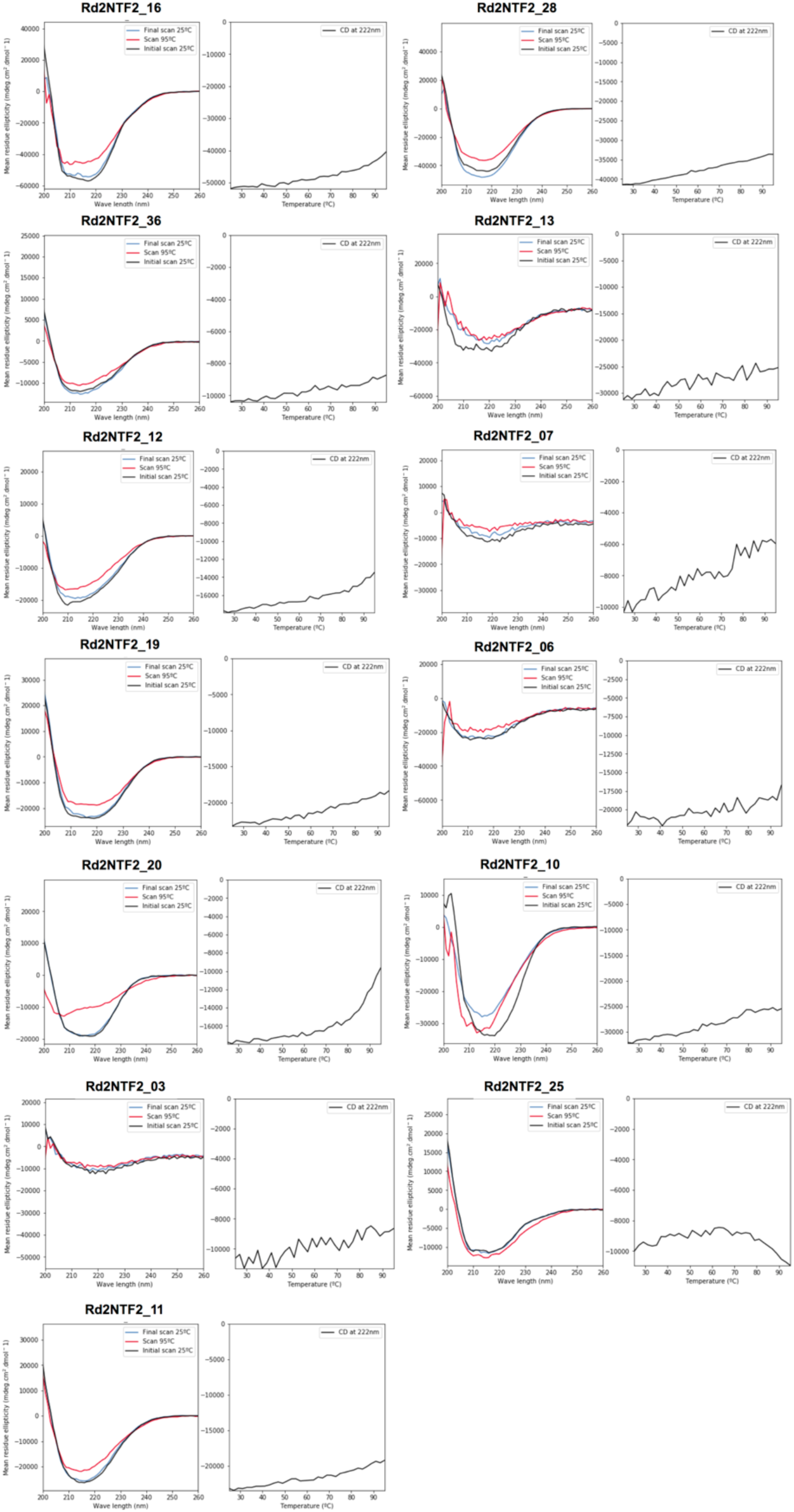
Experimental characterization of second-round designs. Circular dichroism spectra at 25°C and 95°C and temperature curves following circular dichroism at 222nm.

**Fig. S26.**
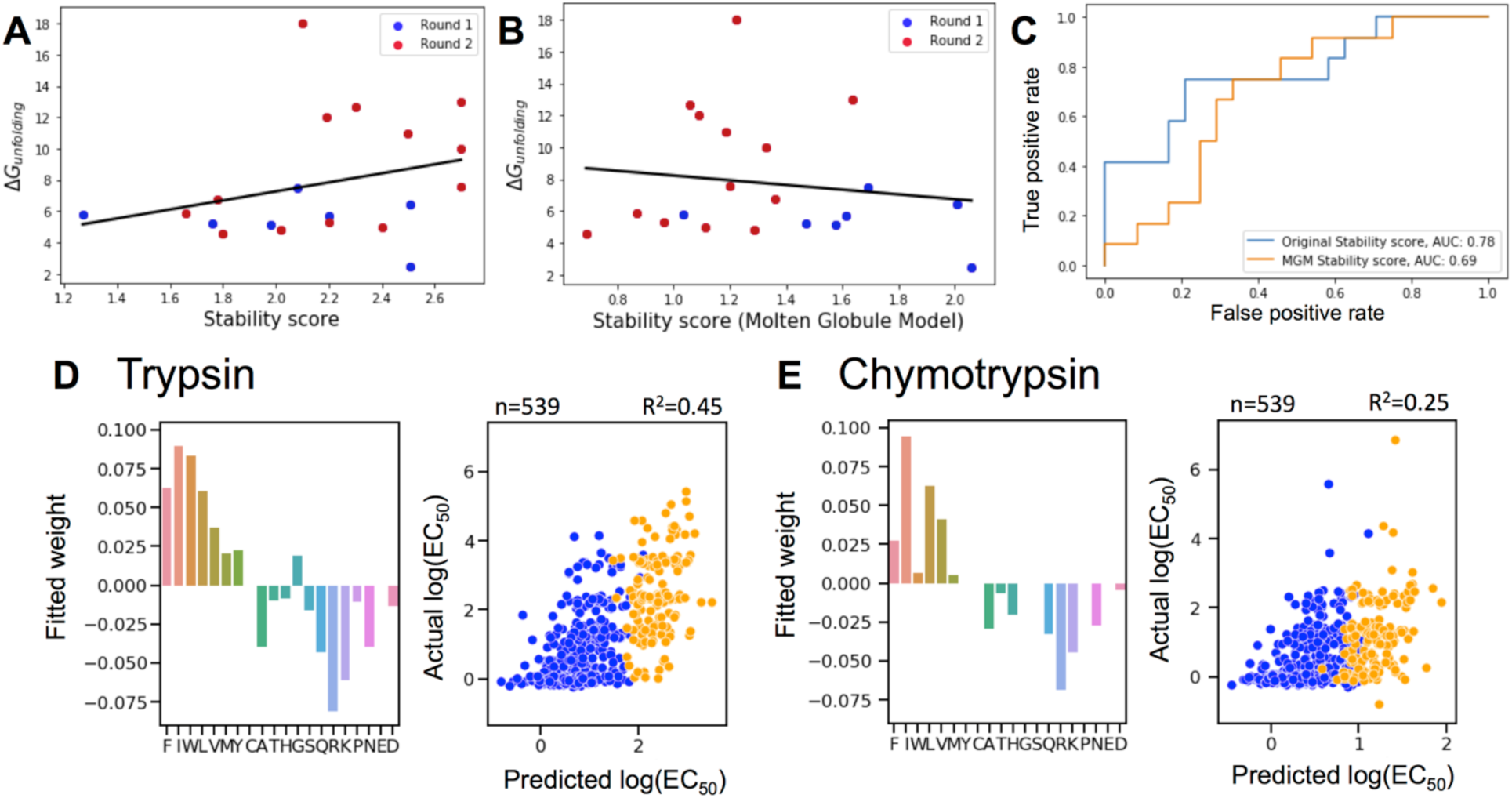
Molten globule model. **A:** ΔG of unfolding vs. stability score calculated usign the unfolded-state model, with data points colored by design round. The black line is a linear fit, with Pearson R 0.3 and p-value 0.182. **B:** ΔG of unfolding vs. stability score calculated usign the “molten globule” model, with data points colored by design round. The black line is a linear fit, with Pearson R 0.07 and p-value 0.753. **C:** Receiver operating characteristic curve of stability score as classifier for folded designs (True) and designs that do not express or not fold (False). **D:** Per-amino acid type weigths in the elastic network model for trypsin (left), and scrambled-sequence precited vs measured EC50 for a validation sequence set, blue dots are sequences from the first round, orange ones are from the second (right). **E:** Per-amino acid type weigths in the elastic network model for chymotrypsin (left), and scrambled-secquence precited vs measured EC50 for a validation sequence set, blue dots are sequences from the first round, orange ones are from the second (right).

**Fig. S27.**
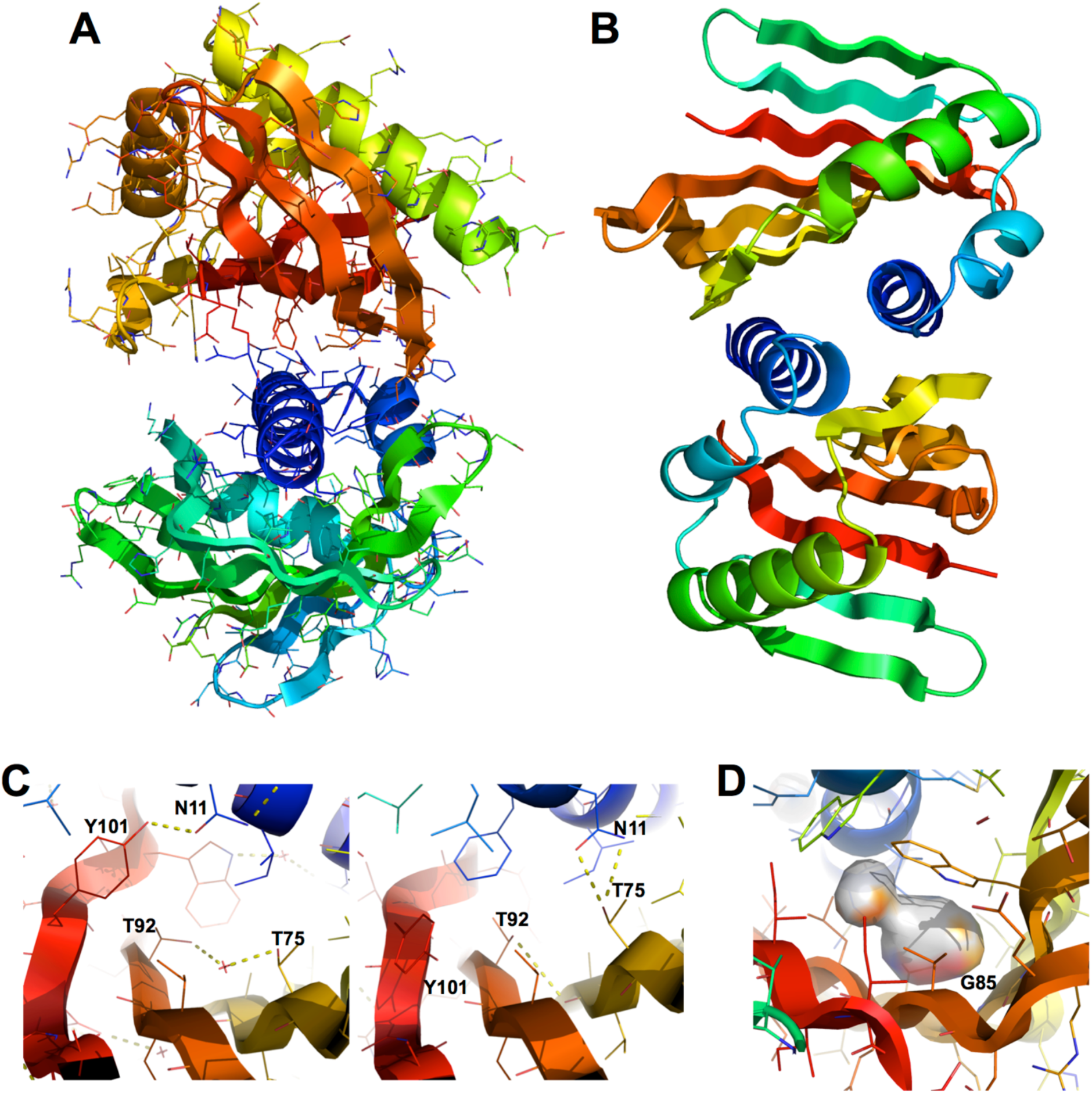
Rd2NTF2_20 and Rd2NTF2_16 crystal structures. **A:** Crystal contacts in Rd2NTF2_20 involving H1 and H2. **B:** Rd2NTF2_16 asymmetric unit, with two copies of the Rd2NTF2_16 monomer interacting through H1, which is displaced from its modeled location. **C:** Rd2NTF2_20 core packing in the crystal structure (left) and model (right). **D:** Core packing of the Rd2NTF2_16 model, showing the cavity near G85.

**Fig. S28.**
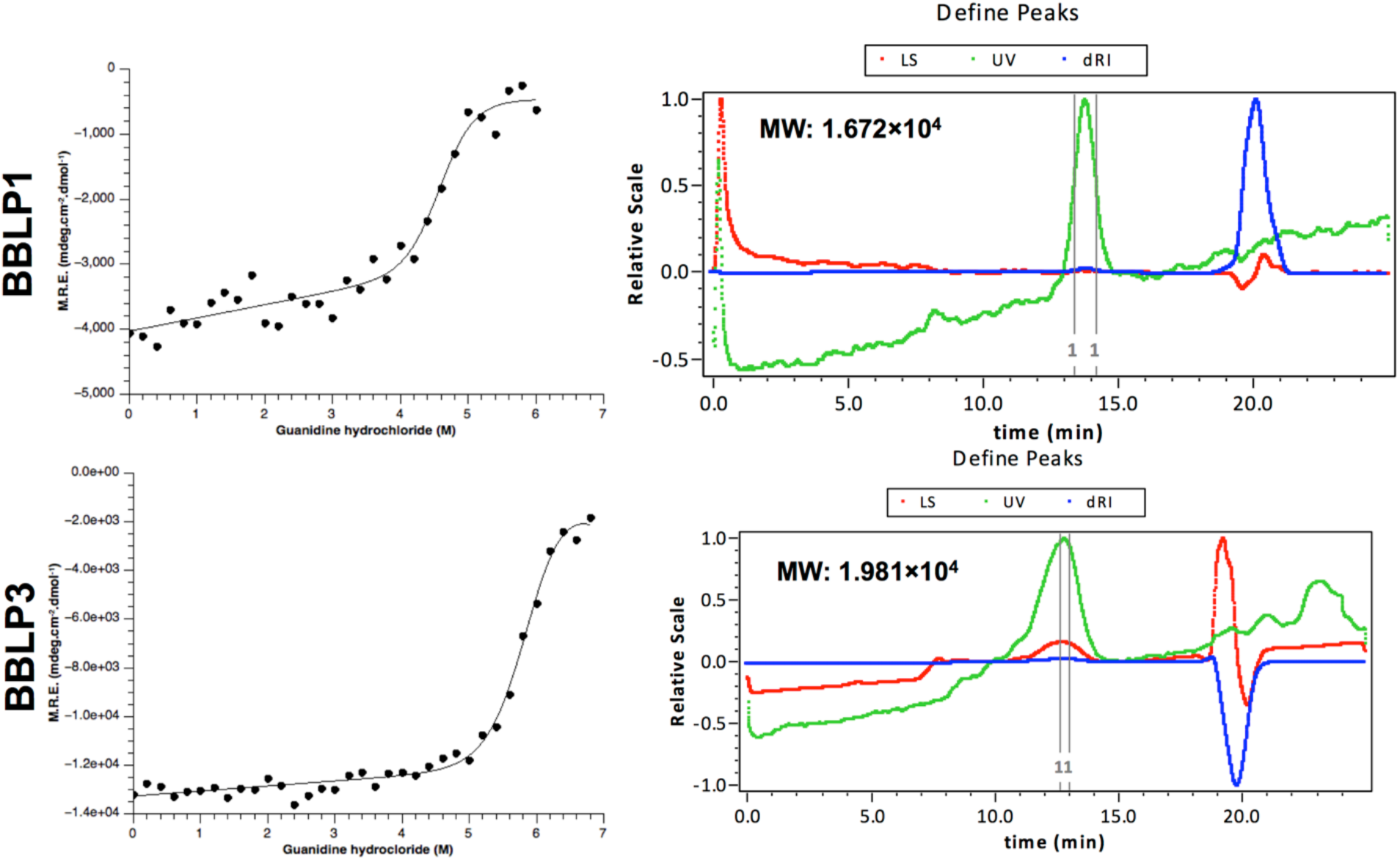
Experimental characterization of NTF2 designs longer than 120 amino-acids. Column 1: Guanidine hychloride titrations following circular dichroism at 222nm. Column 2: Size-exclusion chromatography followed by multi-angle light scattering.

**Fig. S29.**
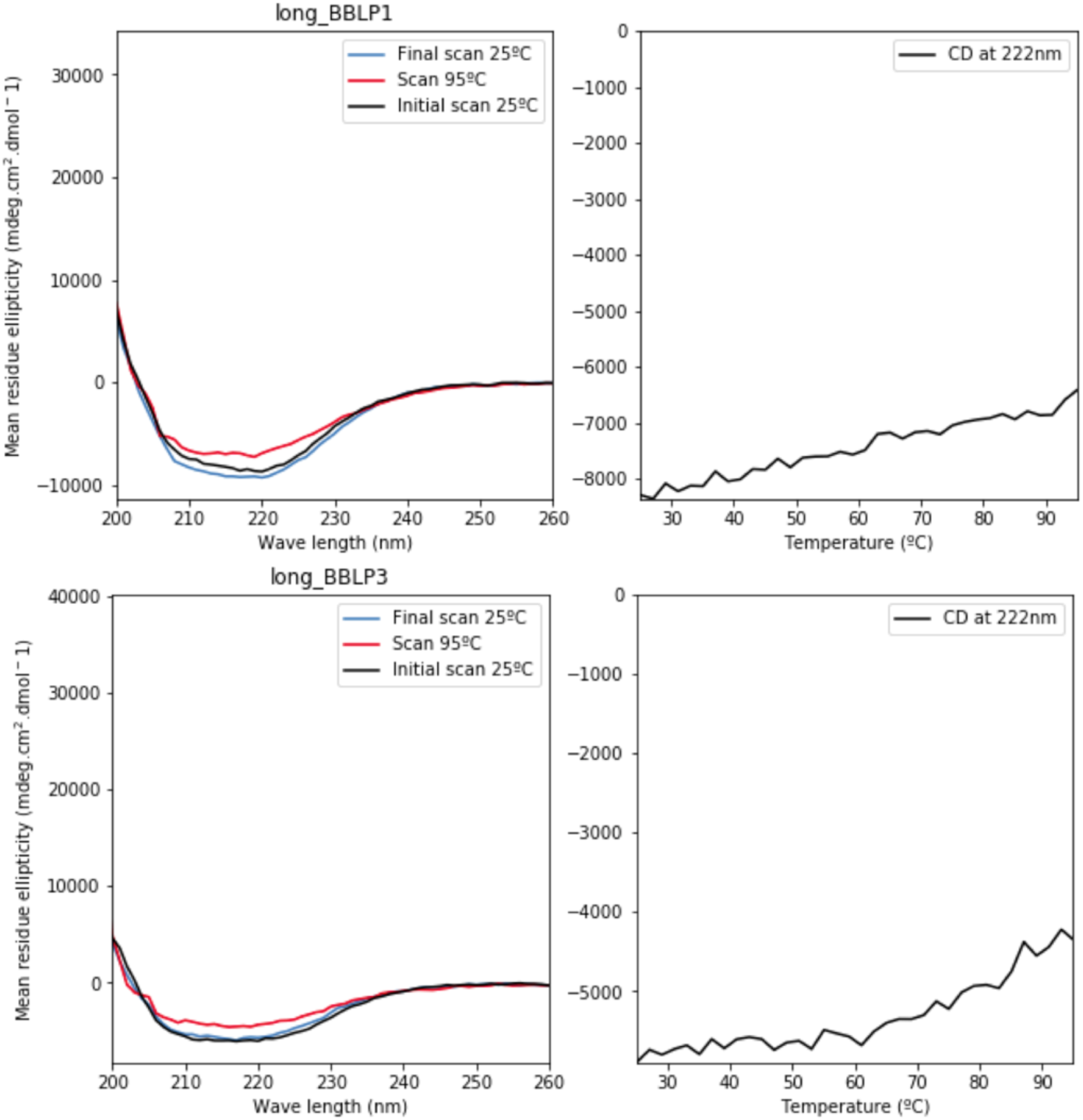
Experimental characterization of NTF2 designs longer than 120 amino-acids. Circular dichroism spectra at 25°C and 95°C and temperature curves following circular dichroism at 222nm.

**Fig. S30.**
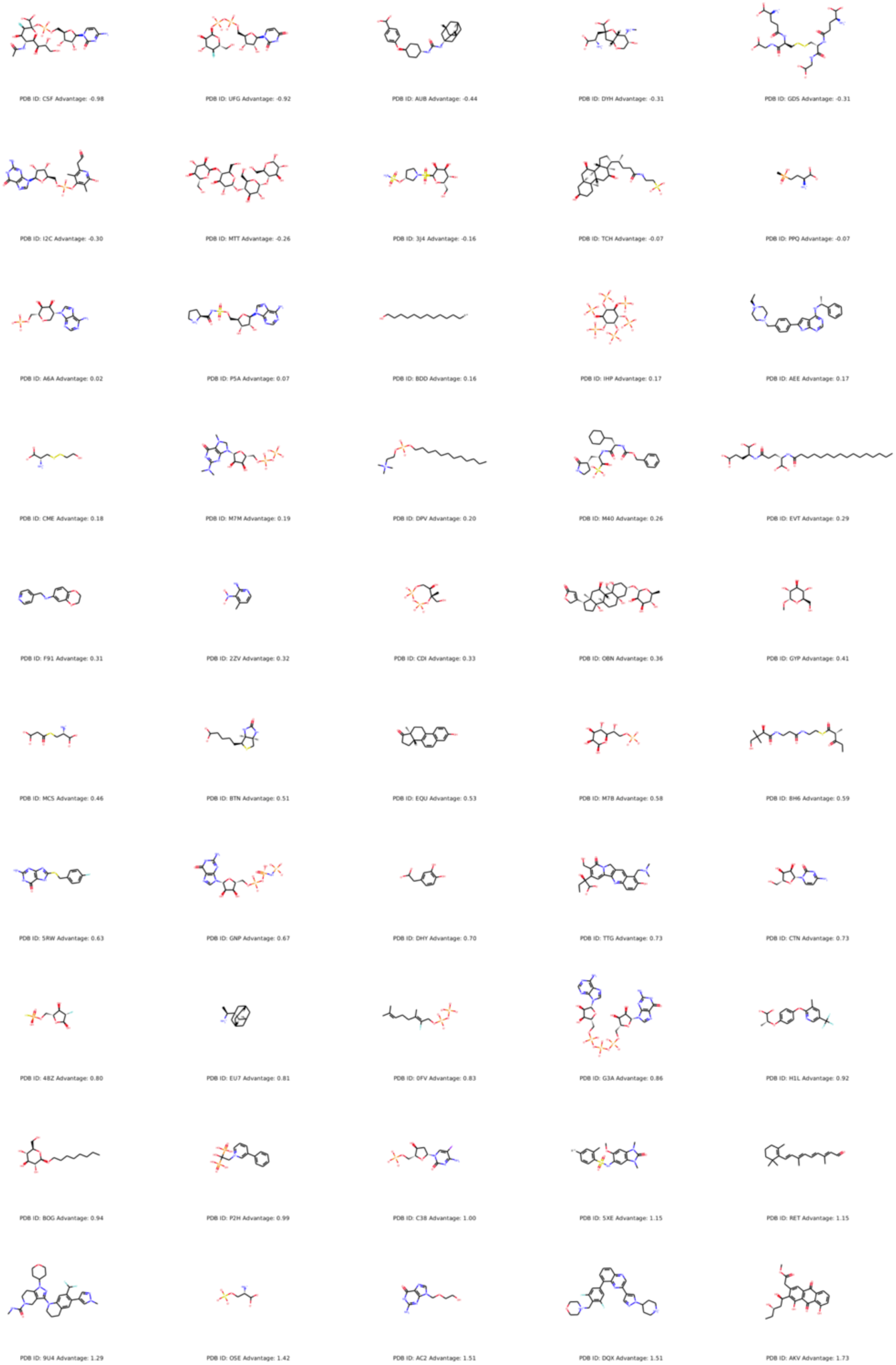
Ligand structures used in the docking benchmark, long with PDB name and advantage value (ΔZ-score).

**Fig. S31.**
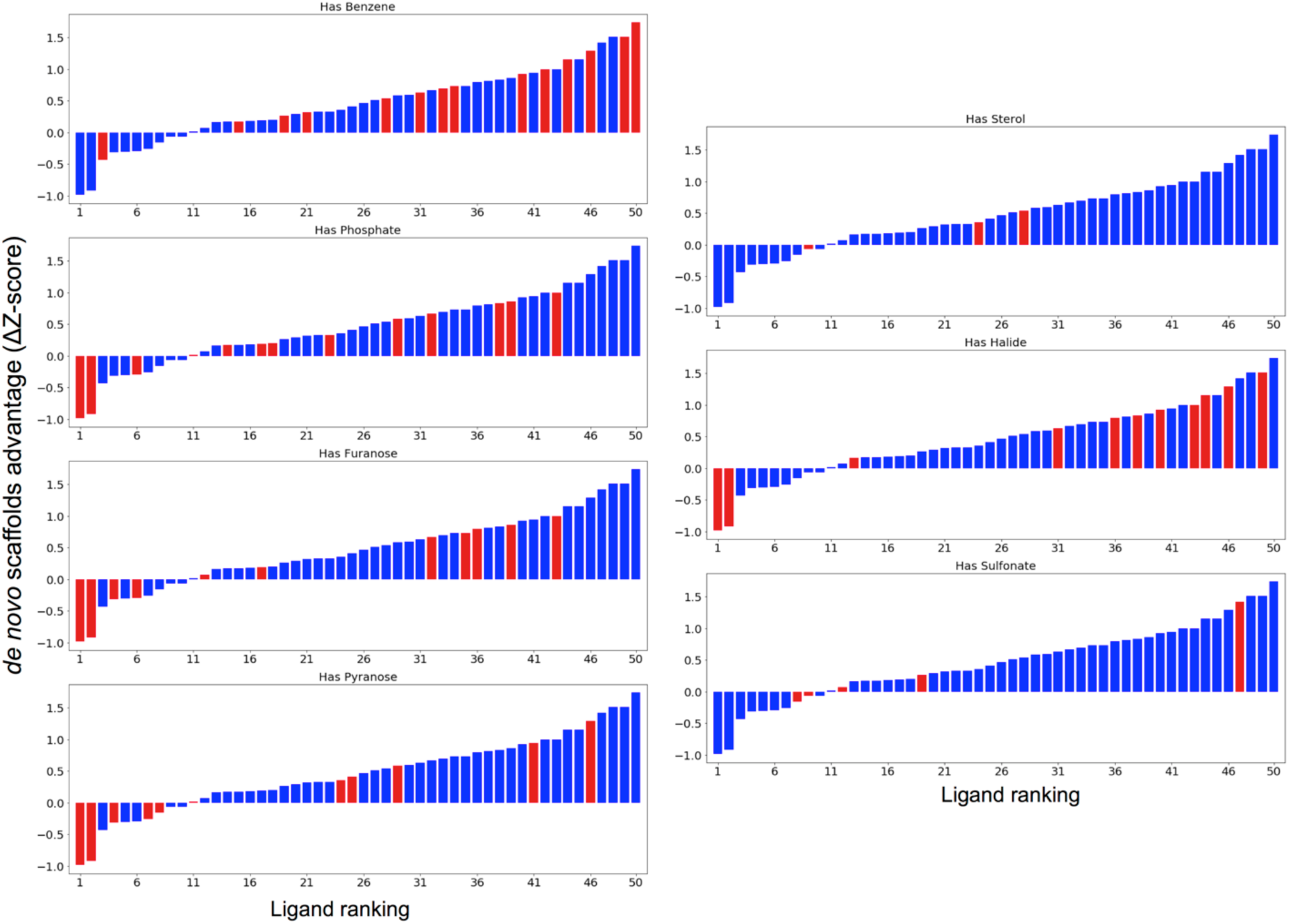
Advantage ranking, colored by presence (red) of a particular chemical moiety (plot title on top).

**Fig. S32.**
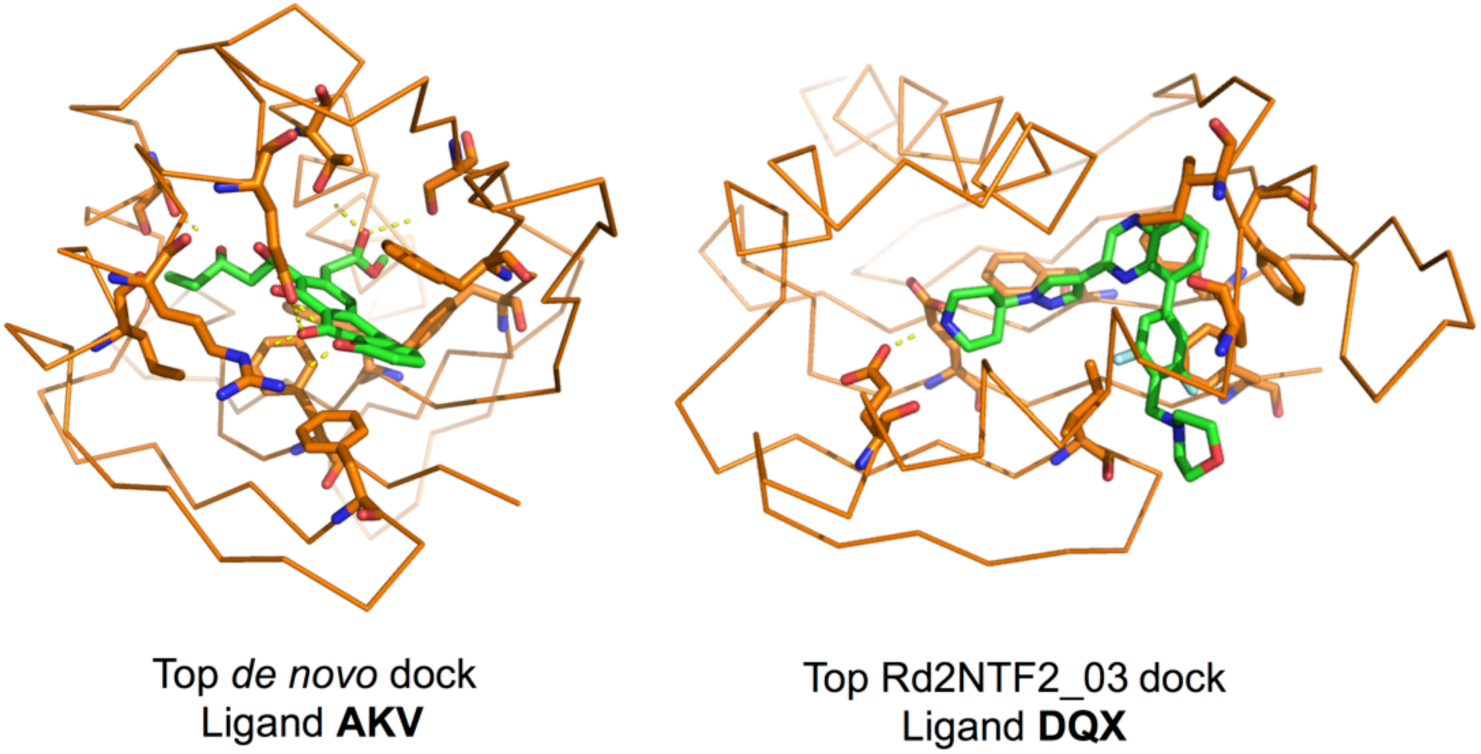
Ligand-scaffold complexes for the two top *de novo* docks (ligands AKV and DQX). Hydrogen bonds are shown as yellow dotted lines.

**Fig. S33.**
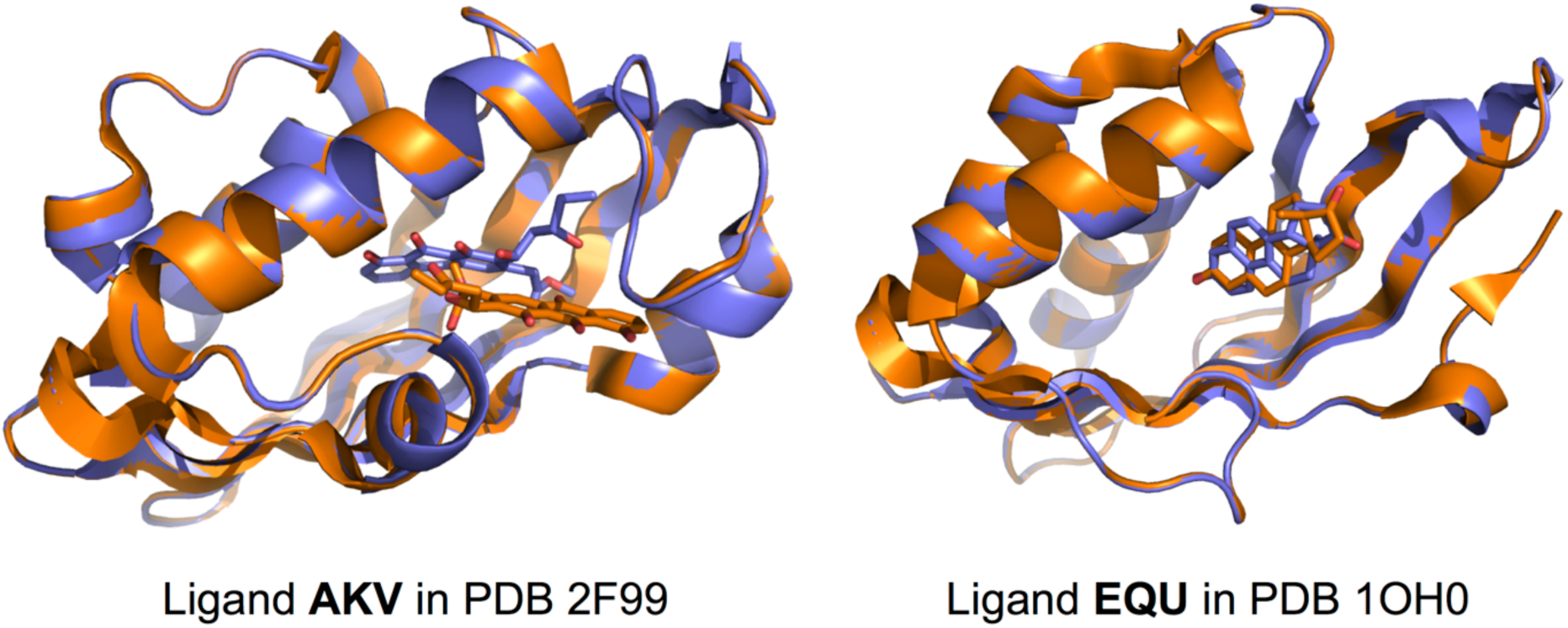
Ligand-scaffold complexes for the two control ligands (AKV and EQU), on their native scaffolds. Purple: RIF docked complex. Orange: original PDB structure. While an orientation similar to the original was recovered for EQU on the 1OH0 backbone, differences between the docked AKV conformer and the PDB conformer likely caused docks in the 2F99 backbone to be substantially different from the PDB structure. These controls suggest the proposed docking test can recover native-like pockets when the same ligand congener is used.

**Fig. S34.**
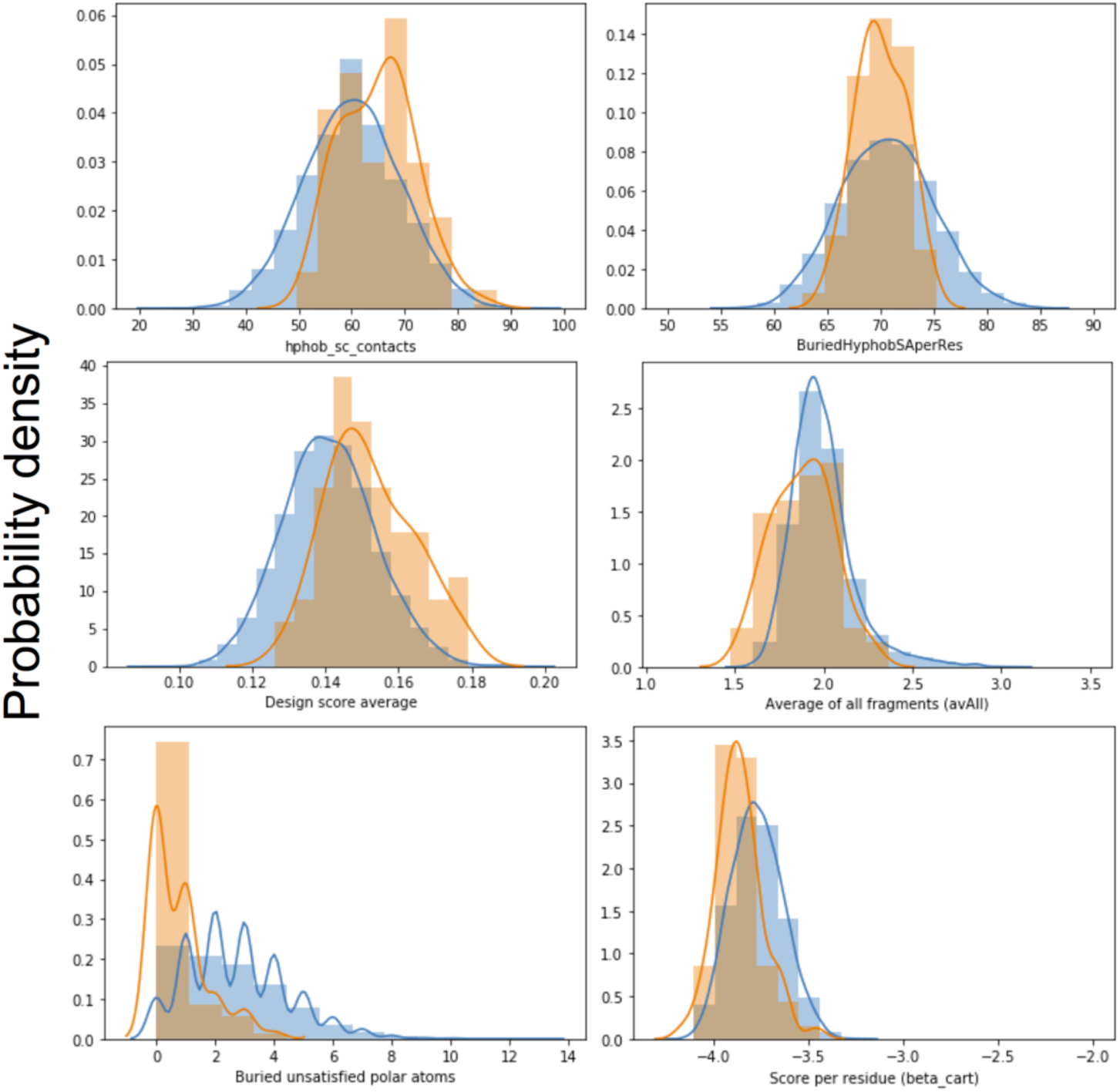
Top stability-predicting features improved in the last version of the generative algorithm. Orange: Final-round designs, blue: second-round designs.

**Table S1.** Generative algorithm parameters

| Parameter<br>(short name) | Units (Allowed<br>values) | Explanation |
| --- | --- | --- |
| **Stage 1** |  |  |
| **Base width**<br>(base_width) | Residues (3,5) | Number of residues between relative positions of main bulges on strand 3 and 6, including the B ABEGO bulge residue. |
| **Long arm length**<br>(long_arm_l) | Pairs of residues<br>(2,3,4) | Number of residue pairs between the main bulge in strand 3 and the N-terminus of strand 3, does not take into account additional bulge residues when strand 3 has additional bulges. |
| **Short arm length**<br>(short_arm_l) | Pairs of residues (1,2) | Number of residue pairs between the main bulge in strand 6 and the C-terminus of strand 6. |
| **Additional bulge on the long arm**<br>(Second_bulge_E3) | Boolean (True,False) | Presence or not of a second bulge on strand 3. |
| **Placement of second bulge on strand 3**<br>(Second_b_place) | Pairs of residues<br>(1,2,null) | Position of the second bulge on strand 3, relative to the main bulge, towards the N-terminus of strand 3. If Second_bulge_E3 is False, then Second_b_place is null. |
| **Degree of curvature angle between long arm and base**<br>(E3_MainBulgeCurve) | $(160-10 \cdot X)^\circ$, where X is one of the allowed values (1,2,3) | Average value for harmonic angle constraint centered in the strand 4 residue that is paired to the main strand 3 bulge. |
| **Degree of curvature angle centered at the E3 second bulge**<br>(E3_SecBulgeCurve) | Null or $(160-10 \cdot X)^\circ$, where X is one of the allowed values (1,2,3,null) | Average value for harmonic angle constraint centered in the strand 4 residue that is paired to the second strand 3 bulge. If Second_bulge_E3 is False, then E3_SecBulgeCurve is null. |
| **Extension of strand 6**<br>(ExtendedE6) | Boolean (True,False) | If True, strand 6 is extended by 2 residues on its C-terminus. This is only compatible with a low degree of strand curvature on the main bulge. |
| **Extension of strand 4**<br>(ExtendedE4) | Boolean (True,False) | If True, strand 4 is extended by 2 residues on its N-terminus. This is only compatible with a short arm of length 2. |
| **Small degree of curvature on the long arm**<br>(CurvedLongArm) | Boolean (True,False) | If True, impose a $150^\circ$ angle constraint on the central residues of the long arm on strands 3 and 4 to impart a small curvature in the absence of a second bulge on strand 3. |
| **Stage 2** |  |  |
| **H3 length** (h_len) | Residue number (10,11,14,15) | Length, in residues, of H3 |
| **H3-S3 loop connection type** (connection_type) | Categorical - Loops with defined ABEGO strings (See table 3) | See table 3 |
| **Frontal hairpin length** (hairpin_len) | Strand residue pair number (4,6) | Length, in residue pairs, of the hairpin formed by S1 and S2 (See Fig. S2C) |
| **Stage 3** |  |  |
| **Opening placement** (Opening) | Categorical (Classic,Alternative) | The pocket opening on NTF2s can be placed either between the frontal hairpin and H3 (Classic), or between H3 and the H1-H2 connection (Alternative). |
| **H1 length** (h1_len) | Residue number (23,19,14) | Length, in residues, of H1 |
| **H2 length** (h2_len) | Residue number (11,7) | Length, in residues, of H2 |
| **C-terminal helix** (has_cHelix) | Boolean (True,False) | Presence or not of a C-terminal helix |
| **Stage 4 (optional)** |  |  |
| **C-helix length** (h_len) | Residue number (11,8) | Length, in residues, of the C-terminal helix. |

**Table S2.** Logic check for sheet construction.

| Stage 1 logic check | Explanation |
| --- | --- |
| All variables must have allowed values | Unexpected values outside of those described in Table 1 (Stage 1) are not allowed. |
| If E3_MainBulgeCurve = 1, then ExtendedE6 must be False | As explained in Chapter 3, sheets with high degree of curvature require bulges to alleviate the clashes caused by bending. Therefore, it is only possible to extend strand 6 past the curvature center on strand 5 if the degree of curvature is low. |
| If Second_bulge_E3 = True, then Second_b_place and E3_SecBulgeCurve must be integers | If a second bulge is placed on strand 3, a position and curvature for it must be selected |
| If long_arm_l = 2, then Second_bulge_E3 must be False | An additional bulge on strand 3 cannot be placed if the long arm is not long enough. |
| If Second_bulge_E3 = True and long_arm_l = 3, then Second_b_place must be 1 | An additional bulge on strand 3 cannot be placed on a position beyond the N-terminus of strand 3 |
| If base_width = 5, and long_arm_l = 4, then Second_bulge_E3 must be True | A sheet with base width 5 and long arm length 4 (the highest lengths for both sheet components) would be impossible to connect by helix 3, even at the highest degree of main bulge curvature. It is therefore required for sheets these length to have an additional bulge on strand 3 to curve the N-terminus of strand 3 back towards the center of the sheet where it can be connected by helix 3. |
| If base_width = 3, then ExtendedE4 must be True, else, ExtendedE4 must be False | The extension of strand 4 is only compatible with base width 3, and base width 5 is only compatible with non-extended strand 4. This is a rule derived from manual inspection of native NTF2-like domains. |
| If base_width = 3, then short_arm_l must be 2 | When the base width is the shortest, the short arm must provide additional interactions with H1. |
| If long_arm_l > 3 (same as = 4), and Second_bulge_E3 = False, then E3_MainBulgeCurve must be 3 | If the length of the long arm is 4, and there is no second bulge on strand 3, then, in order to avoid making a sheet that is too elongated, the curvature degree at the main bulge must be 3. |
| If long_arm_l > 2, then E3_MainBulgeCurve must be higher than 1 | Regardless of the presence of a second bulge on strand 3, if the long arm has length 3 or 4, its degree of curvature must be 2 or 3 in order to avoid an excessively elongated sheet |
| If long_arm_l = 4, Second_bulge_E3 = True, and Second_b_place = 1, then E3_SecBulgeCurve must be exactly 2. | In the case that we have the longest possible long arm with second bulge on strand 3, and this bulge is close to the main strand 3 bulge, then the degree of curvature at the second bulge must be 2 to avoid the sheet extending too far (E3_SecBulgeCurve = 1), or folding back onto |
|  | itself (E3_SecBulgeCurve = 3). |
| If CurvedLongArm = True, then long_arm_l must be 3 or 4 | Imparting a small degree of curvature on a long arm without a second bulge requires it to be a minimal length of 6 residues, otherwise constraint vertices will target atoms outside the logical range. |
| If Second_bulge_E3 = True, then CurvedLongArm must be False | When a second bulge is present on the long arm, curvature is dictated solely by E3_SecBulgeCurve. |

**Table S3:** H3-S3 connection types and features.

| Short name | Length (# residues) | ABEGO string | Compatible H3 lengths |
| --- | --- | --- | --- |
| **BA** | 2 | BA | 10,14 |
| **GBA** | 3 | GBA | 11,15 |
| **GB** | 2 | GB | 11,15 |
| **ClassicDirect** | 0 | - | 10,14 |
| **BulgeAndB** | 4 | GBAB | 11,15 |
| **BBGB** | 4 | BBGB | 10,14 |

**Table S4.** Description of previously published blueprints and variants, in terms of parameters of the generative algorithm described in this work.

| Subfamily | Base width | Long arm length | Short arm length | C-helix | Opening |
| --- | --- | --- | --- | --- | --- |
| **Mk1** | 5 | 4 | 2 | No | Classic |
| **Mk1.PaCH** | 5 | 4 | 2 | Yes – parallel to long arm | Classic |
| **Mk1.PeCH** | 5 | 4 | 2 | Yes – perpendicular to long arm | Classic |
| **Mk1.TP** | 5 | 4 | 2 | Yes – Occludes classic pocket entrance | Alternative |
| **Mk2** | 3 | 6 | 4 | No | Classic |
| **4B.5** | 7 | 4 | 2 | No | Classic |
| **4B.5.CH** | 7 | 4 | 2 | Yes – parallel to long arm | Classic |
| **4B.7** | 5 | 6 | 2 | No | Classic |
| **4B.7.CH** | 5 | 6 | 2 | Yes – parallel to long arm | Classic |

**Table S5.**
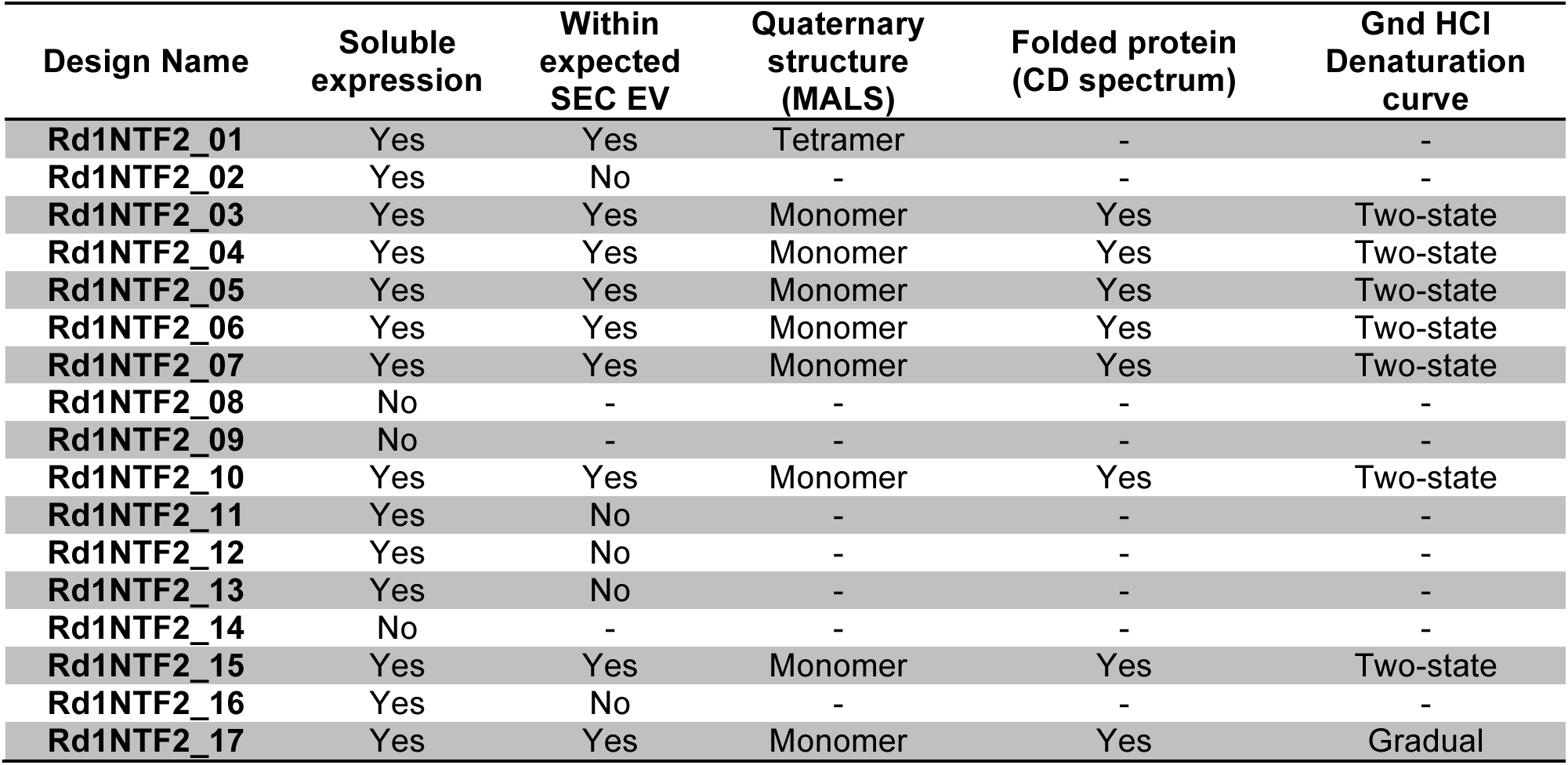
Experimental characterization of first-round designs.

**Table S6.** Biochemical characterization second-round designs.

| Design Name | Soluble expression | Within expected SEC EV | Quaternary structure | Folded protein | Denaturation curve |
| --- | --- | --- | --- | --- | --- |
| Rd2NTF2_01 | No | - | - | - | - |
| Rd2NTF2_02 | No | - | - | - | - |
| Rd2NTF2_03 | Yes | Yes | Monomer | Yes | Two-state |
| Rd2NTF2_04 | No | - | - | - | - |
| Rd2NTF2_05 | No | - | - | - | - |
| Rd2NTF2_06 | Yes | Yes | Monomer | Yes | Two-state |
| Rd2NTF2_07 | Yes | Yes | Monomer | Yes | Two-state |
| Rd2NTF2_08 | Yes | No | - | - | - |
| Rd2NTF2_09 | No | - | - | - | - |
| Rd2NTF2_10 | Yes | Yes | Monomer | Yes | Two-state |
| Rd2NTF2_11 | Yes | Yes | Monomer | Yes | Two-state |
| Rd2NTF2_12 | Yes | Yes | Monomer | Yes | Two-state |
| Rd2NTF2_13 | Yes | Yes | Monomer | Yes | Two-state |
| Rd2NTF2_14 | No | - | - | - | - |
| Rd2NTF2_15 | Yes | Yes | Dimer | - | - |
| Rd2NTF2_16 | Yes | Yes | Monomer | Yes | Two-state |
| Rd2NTF2_17 | No | - | - | - | - |
| Rd2NTF2_18 | No | - | - | - | - |
| Rd2NTF2_19 | Yes | Yes | Monomer | Yes | Two-state |
| Rd2NTF2_20 | Yes | Yes | Monomer | Yes | Two-state |
| Rd2NTF2_21 | Yes | No | - | - | - |
| Rd2NTF2_22 | Yes | No | - | - | - |
| Rd2NTF2_23 | Yes | Yes | Dimer | - | - |
| Rd2NTF2_24 | Yes | No | - | - | - |
| Rd2NTF2_25 | Yes | Yes | Monomer | Yes | Two-state |
| Rd2NTF2_26 | Yes | No | - | - | - |
| Rd2NTF2_27 | Yes | No | - | - | - |
| Rd2NTF2_28 | Yes | Yes | Monomer | Yes | Two-state |
| Rd2NTF2_29 | Yes | No | - | - | - |
| Rd2NTF2_30 | Yes | No | - | - | - |
| Rd2NTF2_31 | Yes | No | - | - | - |
| Rd2NTF2_32 | No | No | - | - | - |
| Rd2NTF2_33 | Yes | No | - | - | - |
| Rd2NTF2_34 | Yes | No | - | - | - |
| Rd2NTF2_35 | Yes | No | - | - | - |
| Rd2NTF2_36 | Yes | Yes | Monomer | Yes | Two-state |
| Rd2NTF2_37 | Yes | No | - | - | - |

**Table S7.** Biochemical characterization of *de novo* NTF2 designs longer than 120 amino acids.

| Design Name | Soluble expression | Within expected SEC EV | Quaternary structure | Folded protein | Denaturation curve |
| --- | --- | --- | --- | --- | --- |
| **BBLP1** | Yes | Yes | Monomer | Yes | Two-state |
| **BBLP2** | Yes | Yes | Dimer | - | - |
| **BBLP3** | Yes | Yes | Monomer | Yes | Two-state |
| **BBLP4** | No | - | - | - | - |
| **BBLP5** | No | - | - | - | - |
| **BBLP6** | No | - | - | - | - |
| **BBLP7** | Yes | No | - | - | - |
| **BBLP8** | Yes | No | - | - | - |
| **BBLP9** | Yes | No | - | - | - |
| **BBLP10** | No | - | - | - | - |

**Table S8:** Design features based on Rosetta filters with score function ref2015

| Feature name | Explanation |
| --- | --- |
| **Holes** | Rosetta filter “Holes”, described in (22), using default values. |
| **HolesCorSCN** | Rosetta filter “Holes”, but only for core atoms, with core defined by the number of side-chain neighbors. |
| **HolesCorSCNnBB** | Rosetta filter “Holes”, but only for core atoms, with core defined by the number of side-chain neighbors, and not taking into account backbone atoms |
| **HolesCorSAS** | Rosetta filter “Holes”, but only for core atoms, with core defined by the solvent accessible surface area of side-chains. |
| **HolesCorSASnBB** | Rosetta filter “Holes”, but only for core atoms, with core defined by the solvent accessible surface area of side-chains, not including backbone atoms. |
| **nres** | Length of the protein in amino-acids |
| **cavity_vol** | Rosetta “CavityVolume” filter with default values |
| **BuriedHyphobSA** | Buried surface area of all atoms in hydrophobic residues (FAMILYVW) as calculated by the “BuriedSurfaceArea” Rosetta filter. |
| **BuriedHyphobSA_H** | Buried surface area of all atoms in hydrophobic residues (FAMILYVW) as calculated by the “BuriedSurfaceArea” Rosetta filter. In helices only. |
| **BuriedHyphobSA_E** | Buried surface area of all atoms in hydrophobic residues (FAMILYVW) as calculated by the “BuriedSurfaceArea” Rosetta filter. In strands only. |
| **BuriedHyphobSA_L** | Buried surface area of all atoms in hydrophobic residues (FAMILYVW) as calculated by the “BuriedSurfaceArea” Rosetta filter. In loops only. |
| **BuriedHyphobSA2_H** | Buried surface area of all atoms as calculated by the “BuriedSurfaceArea” Rosetta filter. In helices only. |
| **BuriedHyphobSA2_E** | Buried surface area of all atoms as calculated by the “BuriedSurfaceArea” Rosetta filter. In strands only. |
| **BuriedHyphobSA2_L** | Buried surface area of all atoms as calculated by the “BuriedSurfaceArea” Rosetta filter. In loops only. |
| **nres_aro** | Number of aromatic residues in the protein |
| **nres_aro_E** | Number of aromatic residues in the protein strands |
| **nres_aro_H** | Number of aromatic residues in the protein helices |
| **nres_aro_L** | Number of aromatic residues in the protein loops |
| **nres_H** | Number of residues in the protein helices |
| **nres_E** | Number of residues in the protein strands |
| **nres_L** | Number of residues in the protein loops |
| **nres_aro_per_res** | Number of aromatic residues in the protein, divided by its length |
| **nres_charge** | Number of charged residues in the protein |
| **nres_hydrophob** | Number of hydrophobic residues in the protein |
| **nres_hydrophob_noA** | Number of hydrophobic residues in the protein, not counting alanine |
| **nAla** | Number of alanine residues in the protein |
| **nres_H_per** | Fraction of residues in helices |
| **nres_E_per** | Fraction of residues in strands |
| **nres_L_per** | Fraction of residues in loops |
| **nres_charge_per** | Number of charged residues in the protein divided by its length *100 |
| **nres_hydrophob_per** | Number of hydrophobic residues in the protein divided by its length*100 |
| **nres_hydrophob_noA_per** | Number of non-alanine hydrophobic residues in the protein divided by its length*100 |
| **nAla_per** | Number of alanine residues in the protein divided by its |
|  | length*100 |
| **BuriedHyphobSAperRes** | Buried surface area of all atoms in hydrophobic residues (FAMILYVW) as calculated by the “BuriedSurfaceArea” Rosetta filter, divided by protein length |
| **total_score** | Total Rosetta score (calculated by ref2015) |
| **scoreRes** | Total Rosetta score (calculated by ref2015) divided by protein length |
| **ramaRes** | Total rama Rosetta score term (calculated by ref2015) divided by protein length |
| **fa_atr** | Total fa_atr Rosetta score term (calculated by ref2015) |
| **fa_atrRes** | Total fa_atr Rosetta score term (calculated by ref2015) divided by protein length |
| **fa_repRes** | Total fa_rep Rosetta score term (calculated by ref2015) divided by protein length |
| **charge** | Absolute protein charge (Assuming typical amino-acid behavior at pH7) |
| **hx_sc** | Shape complementarity between helices and the rest of the protein (See SSShapeComplementarityFilter filter in Rosetta) |
| **longestPS** | Length of the longest continuous stretch of polar amino-acids |
| **longestPS_H** | Length of the longest continuous stretch of polar amino-acids in helices |
| **longestPS_E** | Length of the longest continuous stretch of polar amino-acids in strands |
| **longestPS_L** | Length of the longest continuous stretch of polar amino-acids in loops |
| **exposedHyphob** | Number of solvent-exposed hydrophobic residues (See ExposedHydrophobics filter in Rosetta) |
| **SSmismatch** | Nth root of the product of all residue probabilities of NOT being in the modeled secondary structure state, as calculated by PSIPRED (23), where N is the length of the protein. See the SSPrediction filter in Rosetta. |
| **hb_lr_bb_per_res** | hb_lr_bb Rosetta score term, divided by protein length |
| **hb_lr_bb_per_sheet** | hb_lr_bb Rosetta score term, divided by the number of residues in sheets |
| **hb_sr_bb_per_helix** | hb_sr_bb Rosetta score term, divided by the number of residues in helices |
| **av_loop_rama_prepro** | rama_prepro Rosetta score term in loops, divided by the number of residues in loops |
| **av_loop_p_aa_pp** | p_aa_pp Rosetta score term in loops, divided by the number of residues in loops |
| **av_rama_pp_loop** | av_loop_rama_prepro+ av_rama_pp_loop |
| **geom_res** | Number of residues with large deviations of Omega angle from planarity |
| **AvDeg** | Average number of residues in contact with each position in the protein (See AverageDegree filter in Rosetta) |
| **arom_in_core_SS_SCN** | Number of aromatic amino-acids in core positions of non-loop secondary structure elements, with core defined by the number of side-chain neighbors |
| **arom_in_core_SS_SASA** | Number of aromatic amino-acids in core positions of non-loop secondary structure elements, with core defined by solvent accessible surface area |
| **hyphob_in_core_SS_SCN** | Number of hydrophobic amino-acids in core positions of non-loop secondary structure elements, with core defined by the number of side-chain neighbors |
| **hyphob_in_core_SS_SASA** | Number of hydrophobic amino-acids in core positions of non-loop secondary structure elements, with core defined by solvent |
|  | accessible surface area |
| **core_SCN** | Number of core residues, with core defined by the number of side-chain neighbors |
| **core_SASA** | Number of core residues, with core defined by solvent accessible surface area |

**Table S9:** Design features based on Rosetta filters with score function beta_nov16.

| Feature name | Explanation |
| --- | --- |
| **score_res** | Total Rosetta score divided by protein length. |
| **score_res_betacart** | Total Rosetta score divided by protein length, with score calculated taking into account deviations from ideal covalent bonds angles and lengths. |
| **hyphob_contact** | Number of carbon-carbon atomic contacts between hydrophobic residues. |
| **hphob_sc_contacts_rta** | Number of carbon-carbon atomic contacts between hydrophobic residues, not counting alanine. |
| **hyphob_Aro_contact** | Number of carbon-carbon atomic contacts between aromatic residues. |
| **hyphob_contact_norm** | Number of carbon-carbon atomic contacts between hydrophobic residues divided by protein length |
| **hyphob_Aro_contact_norm** | Number of carbon-carbon atomic contacts between aromatic residues divided by protein length |

**Table S10:** Design features based on Rosetta filters related to burial of unsatisfied polar atoms.

| Feature name | Explanation |
| --- | --- |
| **buns_all** | Total number of residues with at least one buried polar unsatisfied atom |
| **buns_nosurf_all** | Total number of residues with at least one buried polar unsatisfied atom, except in exposed residues |
| **buns_nosurf_sc** | Total number of residues with at least one buried polar unsatisfied side-chain atom, except in exposed residues |
| **buns_nosurf_bb** | Total number of residues with at least one buried polar unsatisfied backbone atom, except in exposed residues |

**Table S11:** Design features based on CLIPPERS pocket detection and inventory software.

| Feature name | Explanation |
| --- | --- |
| pckt_vol | Volume of main detected pocket in Å<sup>3</sup> |
| mouth_n | Number of mouths or openings of the main detected pocket |
| mouth_area | Area of the largest mouth of the main detected pocket |
| pckt_maxTD | Shortest distance (Å) between a point in the outer hull and the deepest part of the main detected pocket. |

**Table S12:**
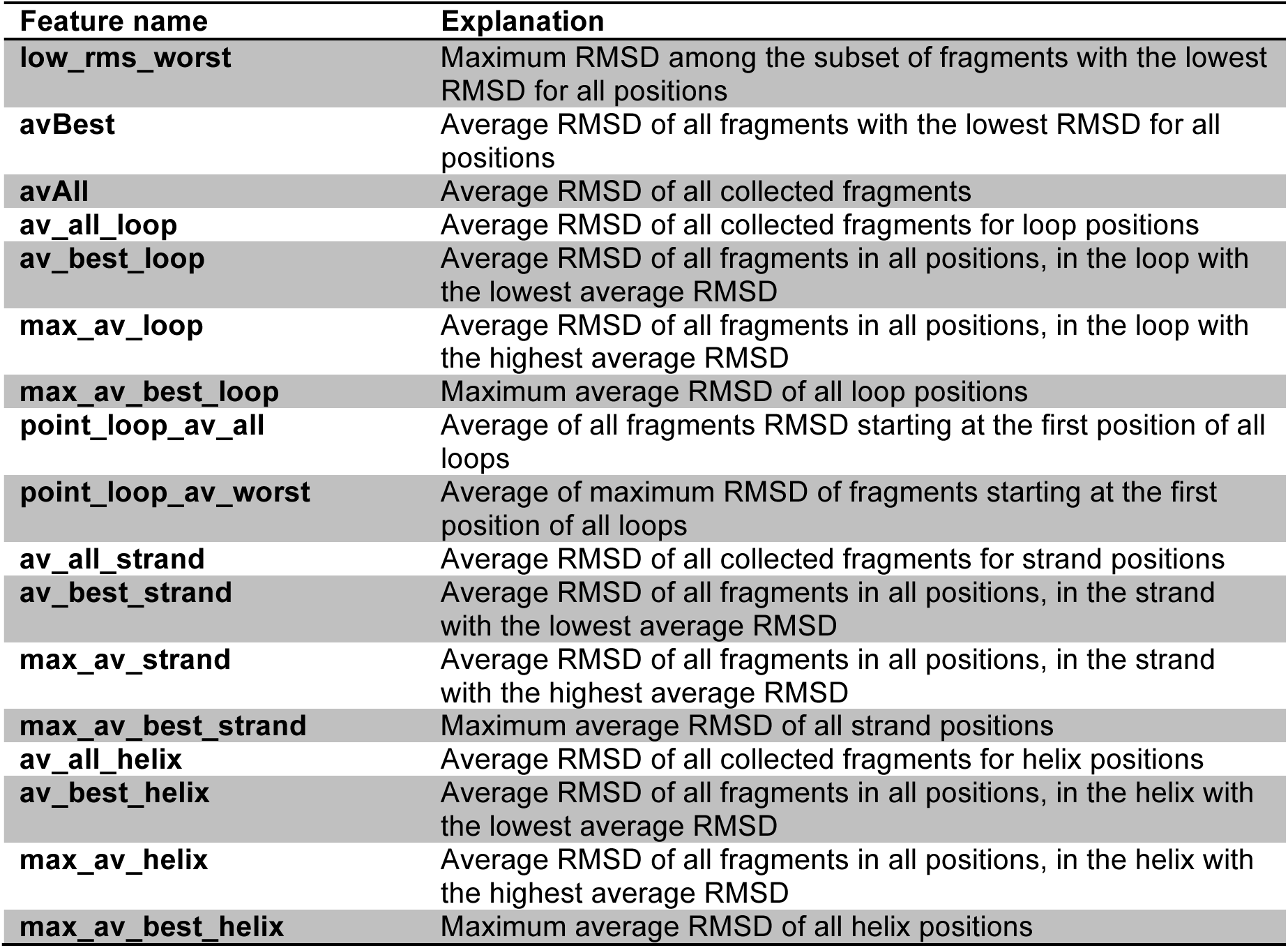
Protein-wide fragment-related features.

| **Feature name** | **Explanation** |
| --- | --- |
| **low_rms_worst** | Maximum RMSD among the subset of fragments with the lowest RMSD for all positions |
| **avBest** | Average RMSD of all fragments with the lowest RMSD for all positions |
| **avAll** | Average RMSD of all collected fragments |
| **av_all_loop** | Average RMSD of all collected fragments for loop positions |
| **av_best_loop** | Average RMSD of all fragments in all positions, in the loop with the lowest average RMSD |
| **max_av_loop** | Average RMSD of all fragments in all positions, in the loop with the highest average RMSD |
| **max_av_best_loop** | Maximum average RMSD of all loop positions |
| **point_loop_av_all** | Average of all fragments RMSD starting at the first position of all loops |
| **point_loop_av_worst** | Average of maximum RMSD of fragments starting at the first position of all loops |
| **av_all_strand** | Average RMSD of all collected fragments for strand positions |
| **av_best_strand** | Average RMSD of all fragments in all positions, in the strand with the lowest average RMSD |
| **max_av_strand** | Average RMSD of all fragments in all positions, in the strand with the highest average RMSD |
| **max_av_best_strand** | Maximum average RMSD of all strand positions |
| **av_all_helix** | Average RMSD of all collected fragments for helix positions |
| **av_best_helix** | Average RMSD of all fragments in all positions, in the helix with the lowest average RMSD |
| **max_av_helix** | Average RMSD of all fragments in all positions, in the helix with the highest average RMSD |
| **max_av_best_helix** | Maximum average RMSD of all helix positions |

**Table S13:**
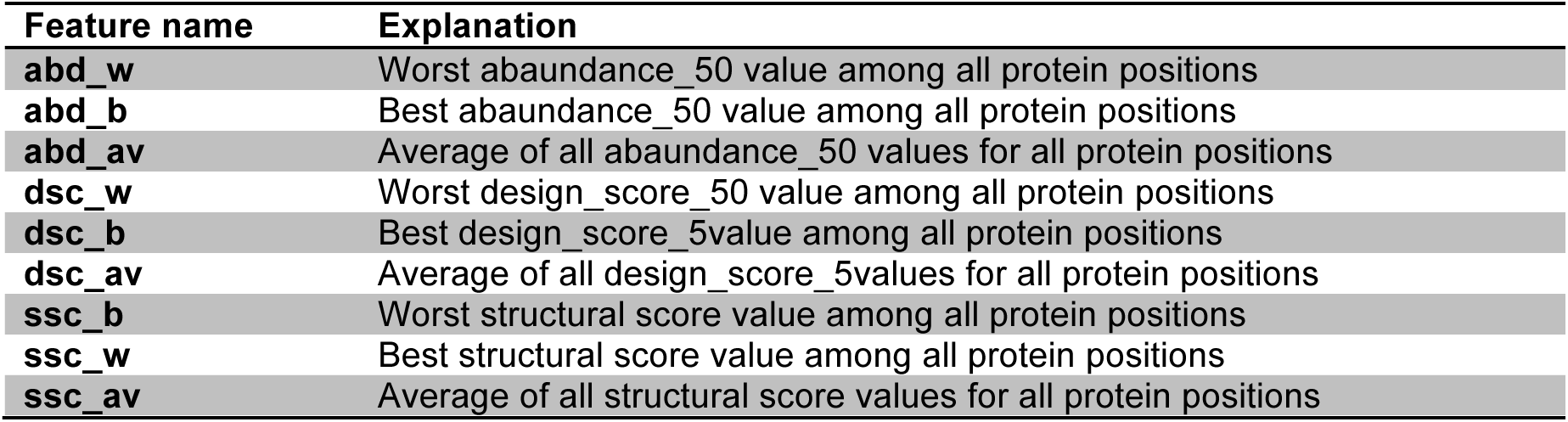
Protein-wide TERM-related features.

**Table S14:** Different local structural domains for TERM and fragment local feature calculation.

| Structure stretch name | Explanation |
| --- | --- |
| N-term_helix | Residues from the N-terminus up to the second to last helix 1 turn |
| H1H2_link | Last H1 turn, loop connection to H2 and first 4 residues of H2 |
| loop3_flank | Loop 3 and flanking residues |
| hairpin | S1 and 2, and connections |
| H3_n | 4 residues of N-terminus of H3 and 3 previous residues |
| H3 | All of H3 |
| H3C_str3 | Last 4 residues of H3, connection to S3 and 5 first residues of S3 |
| str3_4 | 5 last residues of S3 and 5 first residues of S4 |
| str4_5 | 5 last residues of S4 and 5 first residues of S5 |
| str5_6 | 5 last residues of S5 and 5 first residues of S6 |
| str6c_ch | C-terminus of S6 to the C-terminus of the protein, when a C-terminal helix is present. |

**Table S15:** Different ways of calculating TERM and fragment local features.

| Metric name | Explanation |
| --- | --- |
| **av_allfr** | Average of all fragments at all positions |
| **av_bestfr** | Average of only the fragments with the lowest RMSD at all positions |
| **av_worstfr** | Average of only the fragments with the highest RMSD at all positions |
| **best_at_worstfr** | Highest RMSD among the lowest RMSD fragments of all positions |
| **abd50_av** | Average of all abundance_50 values |
| **dsc50_av** | Average of all design_score_50 values |
| **ssc50_av** | Average of all structure score values |
| **abd50_worst** | Worst abundance_50 value among all positions |
| **dsc50_worst** | Worst design_score_50 value among all positions |
| **ssc50_worst** | Worst structural score value among all positions |

**Table S16:** Data collection and refinement statistics. Statistics for the highest-resolution shell are shown in parentheses.

|  | Rd1NTF2_05 | Rd1NTF2_04 | Rd1NTF2_05_I<br>64F_A80G_T9<br>4P_D101K_L1<br>06W | Rd2NTF2_16 | Rd2NTF2_20 |
| --- | --- | --- | --- | --- | --- |
| **Wavelength** | 1 | 1 | 0.9786 | 0.97741 | 0.97741 |
| **Resolution range** | 28.27 - 1.38<br>(1.429 - 1.38) | 44.33 - 1.62<br>(1.678 - 1.62) | 44.72 - 1.83<br>(1.896 - 1.83) | 43.03 - 2.49<br>(2.59 - 2.49) | 42.31 - 1.55<br>(1.58 - 1.55) |
| **Space group** | C 1 2 1 | P 21 21 21 | P 31 | P 21 21 21 | P 41 21 2 |
| **Unit cell** | 60.076 30.498<br>61.099 90<br>97.837 90 | 32.578 36.814<br>177.303 90 90<br>90 | 38.51 38.51<br>134.148 90 90<br>120 | 37.53 73.74<br>86.06<br>90 90 90 | 48.86 48.86<br>84.58<br>90 90 90 |
| **Total reflections** | 100427 (9900) | 132857<br>(13262) | 81860 (8151) | 61433 (7080) | 213398<br>(10368) |
| **Unique reflections** | 22794 (2258) | 28087 (2762) | 19589 (1956) | 8867 (964) | 15581 (781) |
| **Multiplicity** | 4.4 (4.4) | 4.7 (4.8) | 4.2 (4.2) | 6.9 (7.3) | 13.7 (13.3) |
| **Completeness (%)** | 99.65 (99.78) | 93.63 (79.12) | 88.44 (66.92) | 100 (100) | 100 (99.9) |
| **Mean I/sigma(I)** | 19.06 (0.87) | 19.37 (0.96) | 11.07 (0.68) | 9.2 (1.8) | 13.7 (0.8) |
| **Wilson B-factor** | 20.24 | 25.13 | 29.86 | 46.1 | 22.1 |
| **R-merge** | 0.0366 (1.774) | 0.03966<br>(1.586) | 0.06516 (2.12) | 0.133 (1.005) | 0.112 (4.34) |
| **R-meas** | 0.04167 (2.02) | 0.04474<br>(1.779) | 0.07492<br>(2.439) | 0.155 (1.167) | 0.121 (4.68) |
| **R-pim** | 0.01965<br>(0.9543) | 0.02027<br>(0.7912) | 0.03643<br>(1.194) | 0.058 (0.427) | 0.044 (1.74) |
| **CC1/2** | 1 (0.319) | 1 (0.322) | 0.999 (0.432) | 0.996 (0.722) | 0.999 (0.423) |
| **CC\*** | 1 (0.695) | 1 (0.698) | 1 (0.777) | 0.999 (0.898) | 1 (0.788) |
| **Reflections used in refinement** | 22779 (2254) | 26376 (2187) | 17345 (1309) | 8801 (1283) | 15519 (1234) |
| **Reflections used for R-free** | 1997 (196) | 1897 (156) | 1702 (122) | 879 (142) | 1553 (137) |
| **R-work** | 0.1825<br>(0.3084) | 0.2109<br>(0.3592) | 0.2163<br>(0.3734) | 0.2371<br>(0.2712) | 0.1951<br>(0.3692) |
| **R-free** | 0.2158<br>(0.3547) | 0.2403<br>(0.3462) | 0.2546<br>(0.4206) | 0.2910<br>(0.3240) | 0.2279 (4240) |
| **CC(work)** | 0.952 (0.690) | 0.958 (0.647) | 0.951 (0.722) | 0.905 (0.802) | 0.936 (0.666) |
| **CC(free)** | 0.932 (0.612) | 0.947 (0.681) | 0.955 (0.650) | 0.854 (0.684) | 0.925 (0.543) |
| **Number of non-hydrogen atoms** | 1008 | 1879 | 1770 | 2004 | 984 |
| **macromolecules** | 951 | 1776 | 1715 | 1969 | 899 |
| **solvent** | 57 | 103 | 55 | 35 | 77 |
| **Protein residues** | 116 | 217 | 229 | 231 | 107 |
| **RMS(bonds)** | 0.019 | 0.005 | 0.007 | 0.002 | 0.004 |
| **RMS(angles)** | 1.78 | 0.63 | 0.84 | 0.425 | 0.633 |
| **Ramachandran favored (%)** | 100.00 | 99.04 | 100.00 | 94.37 | 98.10 |
| **Ramachandran allowed (%)** | 0.00 | 0.96 | 0.00 | 4.76 | 1.90 |
| **Ramachandran outliers (%)** | 0.00 | 0.00 | 0.00 | 0.87 | 0.00 |
| **Rotamer outliers (%)** | 0.00 | 0.00 | 0.00 | 3.24 | 0.00 |
| **Clashscore** | 8.59 | 3.21 | 5.76 | 6.32 | 4.45 |
| **Average B-** | 29.64 | 41.31 | 48.40 | 52.50 | 29.83 |
| **factor** |  |  |  |  |  |
| **macromolecules** | 29.12 | 41.11 | 48.55 | 52.50 | 28.96 |
| **solvent** | 38.31 | 44.71 | 43.69 | 52.30 | 36.84 |
| **Number of TLS groups** |  | 13 | 12 | 0 | 0 |

**Table S17:**
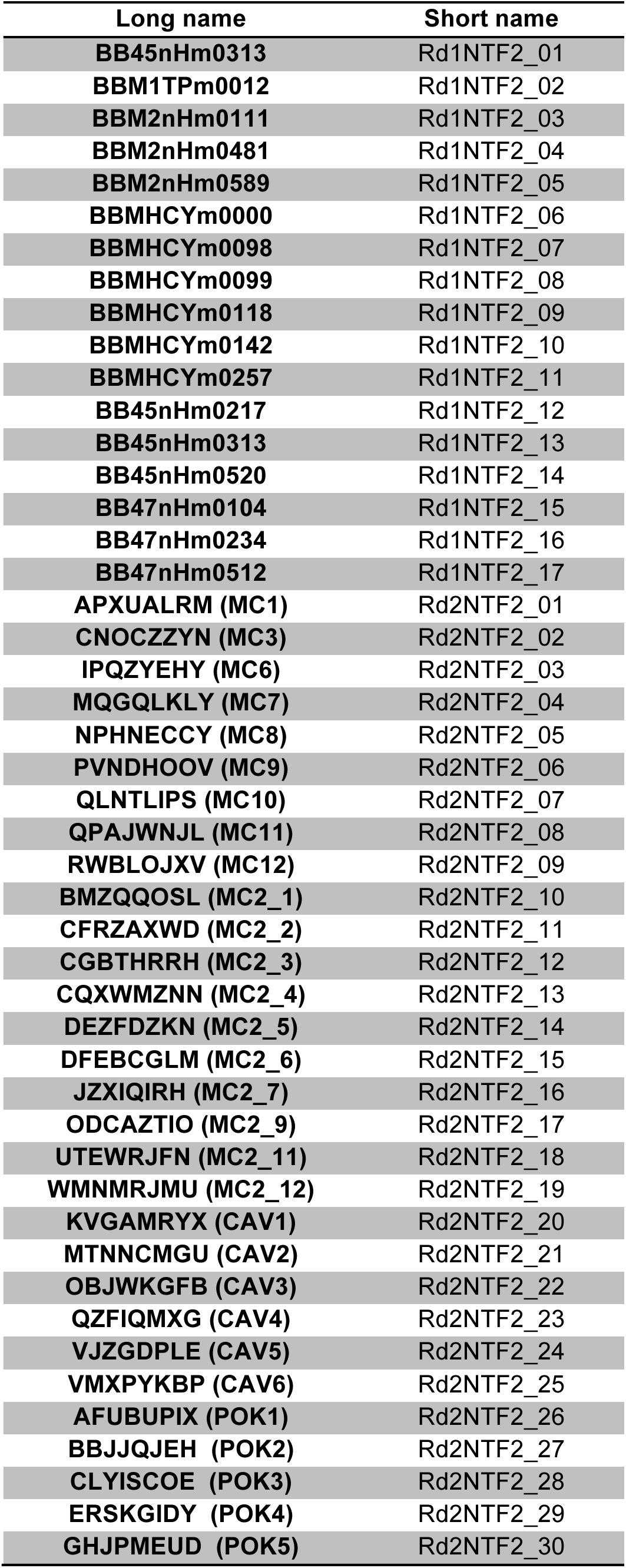
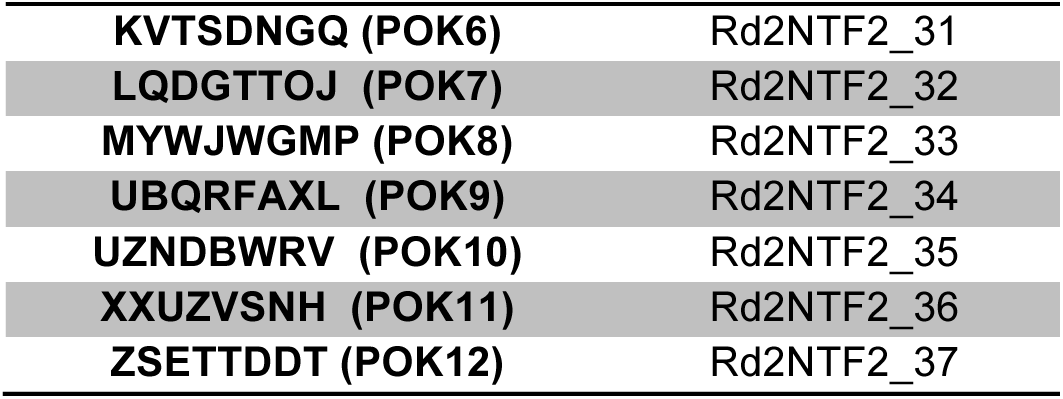
Design name mapping.

